# Co-fluctuations of genes in single cells predict transcriptome-wide outcomes to perturbations

**DOI:** 10.1101/2025.06.27.661814

**Authors:** Benjamin Kuznets-Speck, Nitu Kumari, Irish Senthilkumar, Jaekwon Jung, Hector Manuel Lopez Rios, Leon Schwartz, Hanxiao Sun, Vidyesh Rao Anisetti, Changhu Wang, Qingyang Wang, Pornchanan Pholraksa, Emanuelle I. Grody, Madeline E. Melzer, Benjamin Haley, Ekta Prashnani, Jingyi Jessica Li, Akshit Goyal, Suriyanarayanan Vaikuntanathan, Yogesh Goyal

## Abstract

Pooled single-cell perturbation screens represent powerful experimental platforms for functional genomics, yet interpreting these rich datasets for meaningful biological conclusions remains challenging. Most current methods fall at one of two extremes: either opaque deep learning models that obscure biological meaning, or simplified frameworks that treat genes as isolated units. As such, these approaches overlook a crucial insight: gene co-fluctuations in unperturbed cellular states can be harnessed to model perturbation responses. Here we present CIPHER (Covariance Inference for Perturbation and High-dimensional Expression Response), a conceptual framework leveraging ideas from linear response in statistical physics to model transcriptome-wide perturbation outcomes using gene co-fluctuations in unperturbed cells. We validated our approach on synthetic regulatory networks before applying it to 29 large-scale single-cell genetic perturbation datasets covering **19,003** perturbations and over **6.26**M cells. Our work robustly recapitulated genome-wide responses to single and double perturbations by exploiting baseline gene covariance structure. Importantly, eliminating gene-gene covariances, while retaining gene-intrinsic variances, i.e. “mean-field” conditions, dramatically reduced model performance by several folds across multiple metrics, demonstrating the rich information stored within baseline fluctuation structures. Benchmarked against recent deep learning and linear baselines, CIPHER matched or exceeded the best-performing approaches with fitting a single parameter. Moreover, gene-gene correlations transferred successfully across independent studies of the same cell type, revealing stereotypic fluctuation structures. We further extended CIPHER to the inverse problem of identifying true driver perturbations, where it achieved high performance across both genetic and chemical perturbation screens through uncertainty-aware Bayesian inference. We further used CIPHER to nominate drivers of therapy resistance in melanoma and pancreatic cancer, validating its top predictions experimentally. Finally, most genome-wide responses propagated through the covariance matrix along approximately 1-3 independent and global gene modules, consistent with a low-rank structure of the underlying gene regulatory network, which we show can enable the framework’s success. We have also created a package called `cipher-perturb`, available on PyPI, to apply the framework to any dataset, accompanied by a detailed website (https://goyallab.github.io/CIPHERWebsite/). Our study underscores the importance of theoretically-grounded models in capturing complex biological responses, highlighting fundamental design principles encoded in cellular fluctuation patterns.

## INTRODUCTION

Single-cell profiling technologies enable measurements of cell and tissue molecular makeup at an unprecedented resolution and scale^1–3^. Reference collections of the molecular profiles of individual cells across tissues, organisms, and conditions (“atlases”) comprise multimodal measurements, documenting variability within and across cell types through global consortia. Moving beyond the baseline states catalogued by these atlases, recent breakthroughs in experimental platforms, such as Perturb-seq and CROP-seq^4–11^, have enabled the investigation of genetically-perturbed cells by integrating single-cell RNA sequencing with CRISPR screens. Such approaches are transforming functional genomics by providing a systematic, scalable, and high-resolution view of how genetic perturbations drive phenotypic changes. These large-scale perturbation datasets lay the groundwork for mechanistic models that move beyond correlation, toward truly causal maps of gene regulation in single cells^12,13^.

While powerful, high-content pooled single-cell perturbation screens face significant challenges, including few cells per perturbation in addition to the noise and sparsity inherent in single-cell data^2,12,14,15^. These experimental limitations impede reliable detection of genome-wide perturbation effects, especially when using conventional differential expression analyses and significance testing^12,16^. A suite of computational methods have been developed to tackle these limitations^12^. These approaches broadly fall into two categories: interpretable statistical frameworks that analyze existing perturbation data, and predictive models that forecast cellular responses to unseen perturbations. Statistical frameworks typically estimate the average impact of each perturbation by fitting linear regression models^6,17^. Other interpretable methods use matrix factorization (e.g., SVD, NMF, SMAF) to perform dimensionality reduction and identify gene programs or modules that respond coherently to perturbations^18,19^. A notable recent interpretable framework is TRADE^16^, which models true differential expression distributions while accounting for noise, using a “transcriptome-wide impact” metric to quantify perturbation effects including weak signals missed by conventional significance testing. At the same time, traditional linear regression models have some limitations: they typically treat genes as independent, cannot explicitly identify the genes driving the response (or the degree of impact per driver gene), and, with the exception of TRADE^16^, produce only point estimates of the regression coefficients and are not uncertainty-aware. Deep learning frameworks ^20–22^ model perturbation responses in latent spaces and can predict the outcomes of perturbations. These methods often present “black-box” solutions with complex non-linear mappings, limiting interpretability^23^ of the underlying biological mechanisms, and typically require substantial amounts of training data.

Despite progress, we still lack interpretable, physically-grounded approaches with the ability to decipher the underlying mechanistic basis of genome-wide responses. Nevertheless, techniques from statistical physics help provide a universal framework to model the effect of perturbations from baseline fluctuations in other contexts^24–27^. If similar physically-grounded models could explain genome-wide perturbation responses, what might that imply about the underlying design principles of biological responses?

We present CIPHER (Covariance Inference for Perturbation and High-dimensional Expression Response). CIPHER is a simple linear response framework that can *capture and interpret* genome-wide responses to perturbations across a broad range of experimental and biologically-validated synthetic systems. Specifically, these responses can be *recapitulated* using only the information contained in gene-gene covariances in unperturbed cells, which can be readily estimated from large-scale single-cell RNA sequencing datasets. We first tested our framework in stochastic linear and nonlinear models of gene regulation, including several biologically validated regulatory networks, where the underlying ground truth was known.

Although gene regulatory networks are nonlinear and out of equilibrium, we show theoretically^28–30^—combining statistical physics, nonlinear dynamics, and random matrix theory—that this linear relationship emerges when the network is approximately low-rank, dominated by one or a few global modes (**Box 1**). The theory is robust to intrinsic expression noise and yields independent predictions that we test in what follows.

We next applied CIPHER to large single-cell perturbation sequencing datasets^16,31–35^ and one imaging Perturb-FISH dataset^10^, including 22 CRISPR interference (CRISPRi) gene knockdown and 7 CRISPR gene activation (CRISPRa) datasets, covering **19,003** perturbations, **6,265,800** cells, and 15 cell types (**Table 1**). Importantly, in all these cases, unperturbed gene-gene covariances were essential to successfully recapitulating perturbation responses: eliminating off-diagonal covariances in a “mean-field” model—shuffling expression values across cells independently for each gene, which preserves each gene’s marginal distribution but disrupts gene-gene coupling—dramatically decreased model performance. These correlation structures also transferred robustly across independent studies of the same cell type. Together, these findings demonstrate the rich information contained within baseline gene-gene fluctuation architectures. Benchmarked against five recent deep learning and linear methods across these datasets, CIPHER matched or exceeded the best-performing approaches while fitting a single parameter per perturbation. Moreover, our framework exhibited high performance for identifying true causal drivers given the genome-wide change upon perturbation by leveraging Bayesian inference alongside gene-gene covariances, across both genetic and chemical compound screens. We further used CIPHER to nominate drivers of therapy resistance—to BRAF/MEK inhibitors in melanoma and to KRAS inhibitors in pancreatic cancer—validating its top-ranked genes experimentally. CIPHER also successfully predicted lineage outcomes in hematopoietic differentiation from single-cell clonal barcoding dataset. Together, our results highlight the power of simple theoretical models inspired by statistical physics in probing and dissecting biological complexity.

**Table 1:**
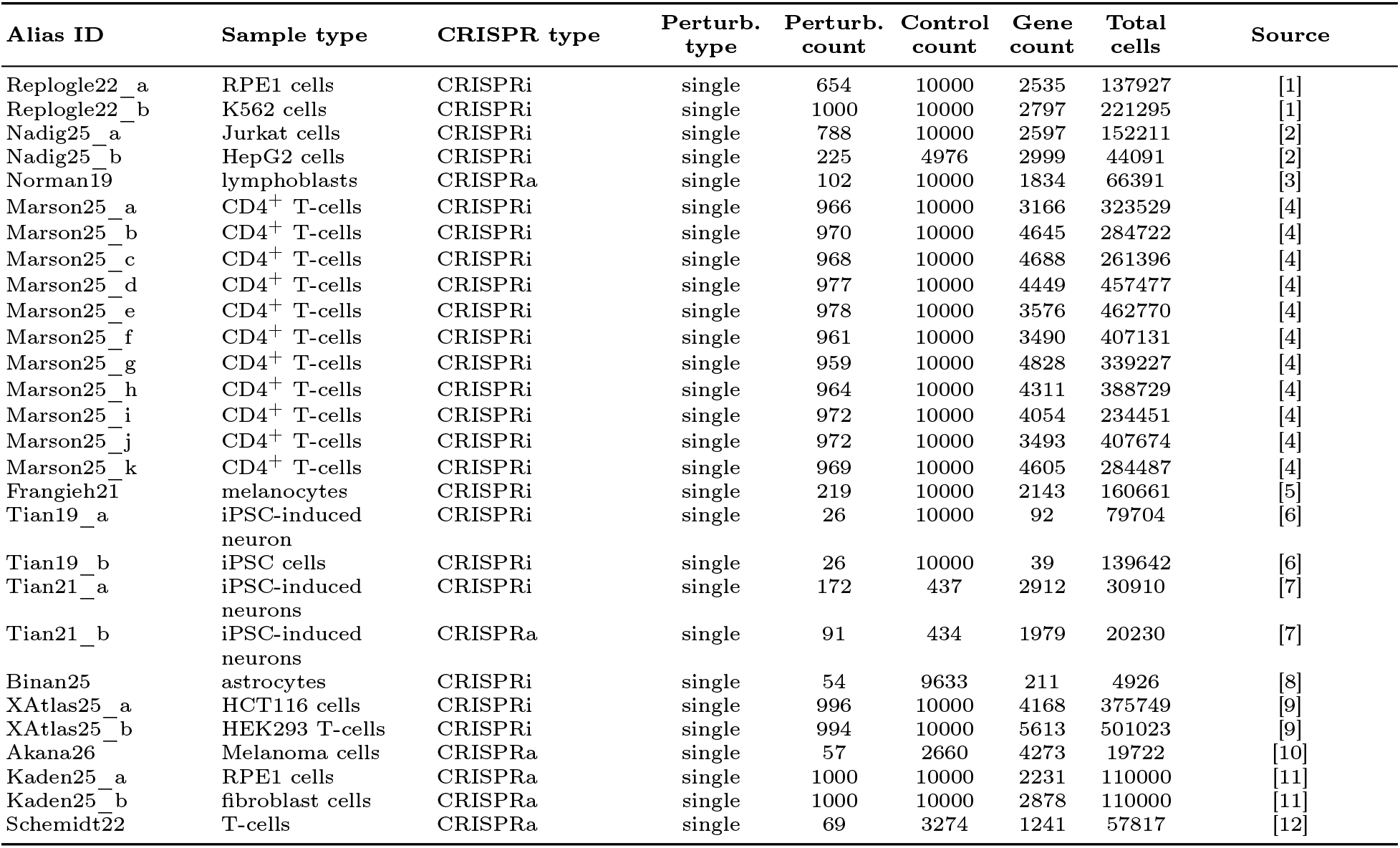
Specifications for single-perturbation subset of the single-cell perturbation datasets used in this study.

## RESULTS

### Overview of CIPHER

Gene correlations have long served as proxies for gene-gene interactions^36–38^. Recent work reveals these correlations contain meaningful information about gene regulatory network dynamics, particularly during bifurcations^39^. Our framework builds on these insights, using correlations to fit perturbation-induced expression changes. We named our framework as CIPHER, noting that the primary goal of our study is to mechanistically interpret perturbation-response rather than predict the response to unmeasured perturbations per se.

CIPHER is based on a simple relationship between gene-gene covariances in unperturbed cells and the mean genome-wide response to a perturbation. Let *X* denote the vector of gene expression levels. We define the response to a perturbation as the difference between the mean expression levels in the perturbed and unperturbed populations, 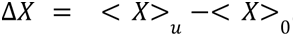. Now let 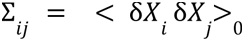 for 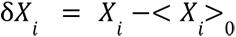 denote the gene-gene covariance matrix estimated across unperturbed cells. Throughout the paper, we denote averages of gene expression *X* over cells as < *X* > and distinguish between the perturbed and control ensembles with subscript *u* and 0 respectively.

CIPHER relates the response Δ*X* to the unperturbed covariance matrix Σ through the following simple equation:

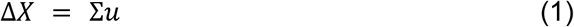

where *u* is the effective perturbation vector that determines the strength with which each gene was perturbed. Equation (1) is the central relationship underlying CIPHER: it states that a genome-wide perturbation response can be represented as a linear combination of gene-gene covariance patterns measured entirely in unperturbed cells.

In a variety of gene expression networks, experimental Perturb-seq datasets, and biological trait experiments (**Fig. 1**), we investigated the utility of this framework using a three-pronged approach. First, we considered the forward problem of fitting perturbation responses given true single or multi-gene perturbations and fluctuation structure Σ,

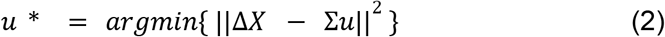

**Fig. 1:**
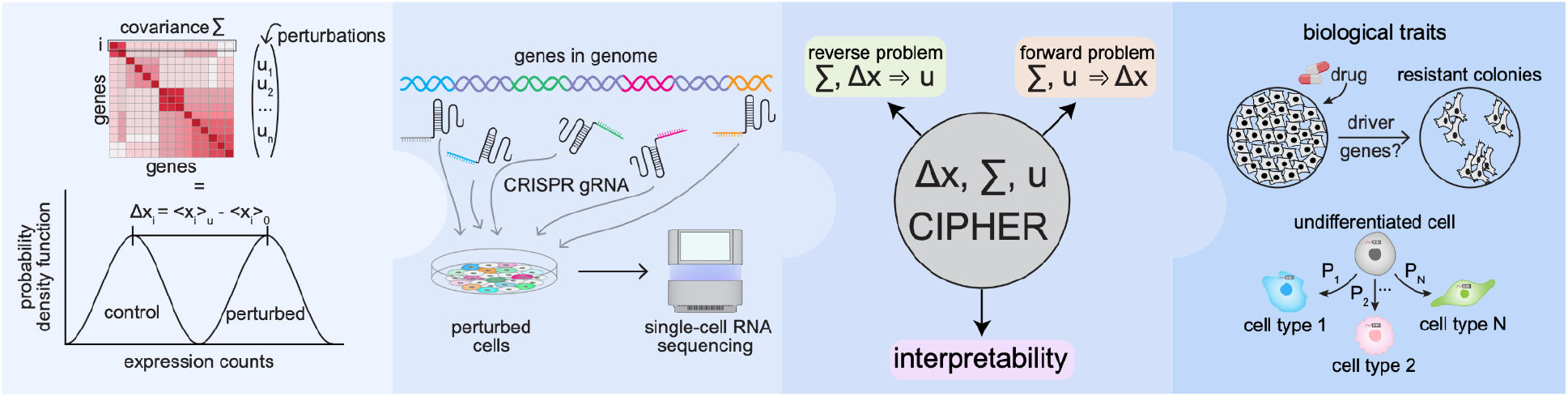
Schematic of the workflow and applications to synthetic and real experimental datasets. (column 1) The gene-gene covariance matrix informs on changes to gene expression upon perturbation. (column 2) Perturb-seq interrogates thousands of perturbations by combining RNA-seq with CRISPR screens. (column 3) Given covariance and expression changes, CIPHER has three complementary modalities. The forward problem fits changes in expression, the reverse problem predicts the driving perturbation(s) and the framework is fully interpretable by telling us 1) how each genes change is made up of contributions from both itself and every other gene and 2) how response is propagated along dominant or subdominant modes of the covariance. (column 4) CIPHER can be readily applied to biological traits. In cancer drug resistance, CIPHER can identify resistance drivers from the fluctuations in naive cancer cells. In differentiation, CIPHER can infer cell fate decisions from the fluctuations of undifferentiated cells.

Equation (2) obtains u* by minimizing the residual ||Δ*X* − Σ*u*|| with respect to *u*. For a single perturbation to gene i, Eq. (2) reduces to fitting a global scalar, that projects the response onto the direction of the i-th covariance column:

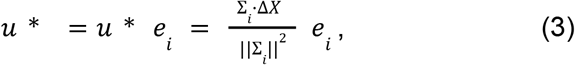

where, *e_i_* is the unit vector for gene i (**Methods 1**). Fitting u* from the observed ΔX in Eq. (3) cannot supply the prediction’s directional content, which comes from the covariance alone. As we detail below, Δ*X* may comprise all cells and genes, or a subset of them, holding a subset out for testing.

Next, we tackled the inverse problem of predicting driver perturbations (direction and magnitude) from Σ and Δ*X*. As detailed in **Methods 1**, we used the central limit theorem to model the noise in the empirical average Δ*X* as a zero-mean Gaussian with covariance 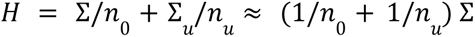, where *n_0/u_* denotes control/perturbation sample size and Σ is the covariance of the perturbed cells. The likelihood over responses is then

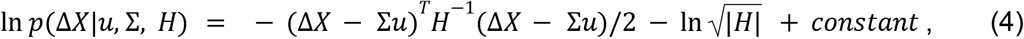

from which we rank perturbations according to their posterior

ln *p*(*u*|Δ*X*, Σ, *H*) = *constant* + ln *p*(Δ*X*|*u*, Σ, *H*) + ln *p*(*u*). We subsequently examined eigengenes, or principal components—groups of co-expressed genes that captured dominant expression patterns—to interpret response as propagating broadly on the gene level but localized to a handful of underlying modes or regulatory modules. Detailed description of our framework as well as corresponding derivations are provided in **Methods 1**. We have also created a package called `cipher-perturb`, available on PyPI, to apply the framework to any dataset, accompanied by a detailed website (https://goyallab.github.io/CIPHERWebsite/).

### Implementation on synthetic toy regulatory networks

We initially validated our framework CIPHER through proof-of-concept testing on a suite of synthetic gene networks (see **Methods 2**). These toy networks provided controllable architectures and interaction parameters with known ground truth for benchmarking. We began with linear systems of random gene regulatory networks (dx/dt = Ax + noise, where A is a matrix of gene-gene interactions and x is a vector of gene expression). We reasoned that our framework should accurately recapitulate perturbation responses within the linear regime (Case 1, see **Methods 2.1**). As Robert May demonstrated in their seminal study^40^, such networks remain stable for N genes when coupling strength 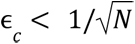, beyond which they bifurcate to become globally unstable. Consistently, increasing coupling strength (**Fig. 2A**) caused gene-gene correlations to become modular and fractal-like (**Fig. 2B**). The transition from the subcritical to supercritical regime is marked by a striking increase in the percent variance explained by the first principal component. Accordingly, a collapse of the dynamics onto a low-dimensional subspace (with low participation ratio) where “team”^41,42^ structure emerges in the covariance as measured by variance explained and participation ratio—a measure of how many components actively contribute to the response (**Fig. 2C,D**). Biologically, this corresponds to the dynamics passing through an unstable fixed point, as would happen, for example, during a differentiation trajectory^43^. Importantly, the predicted responses to genetic perturbations exhibited relatively low coefficient of variation within the linear regime, but diverged at and beyond the critical coupling threshold (**Fig. 2E**).

**Fig. 2:**
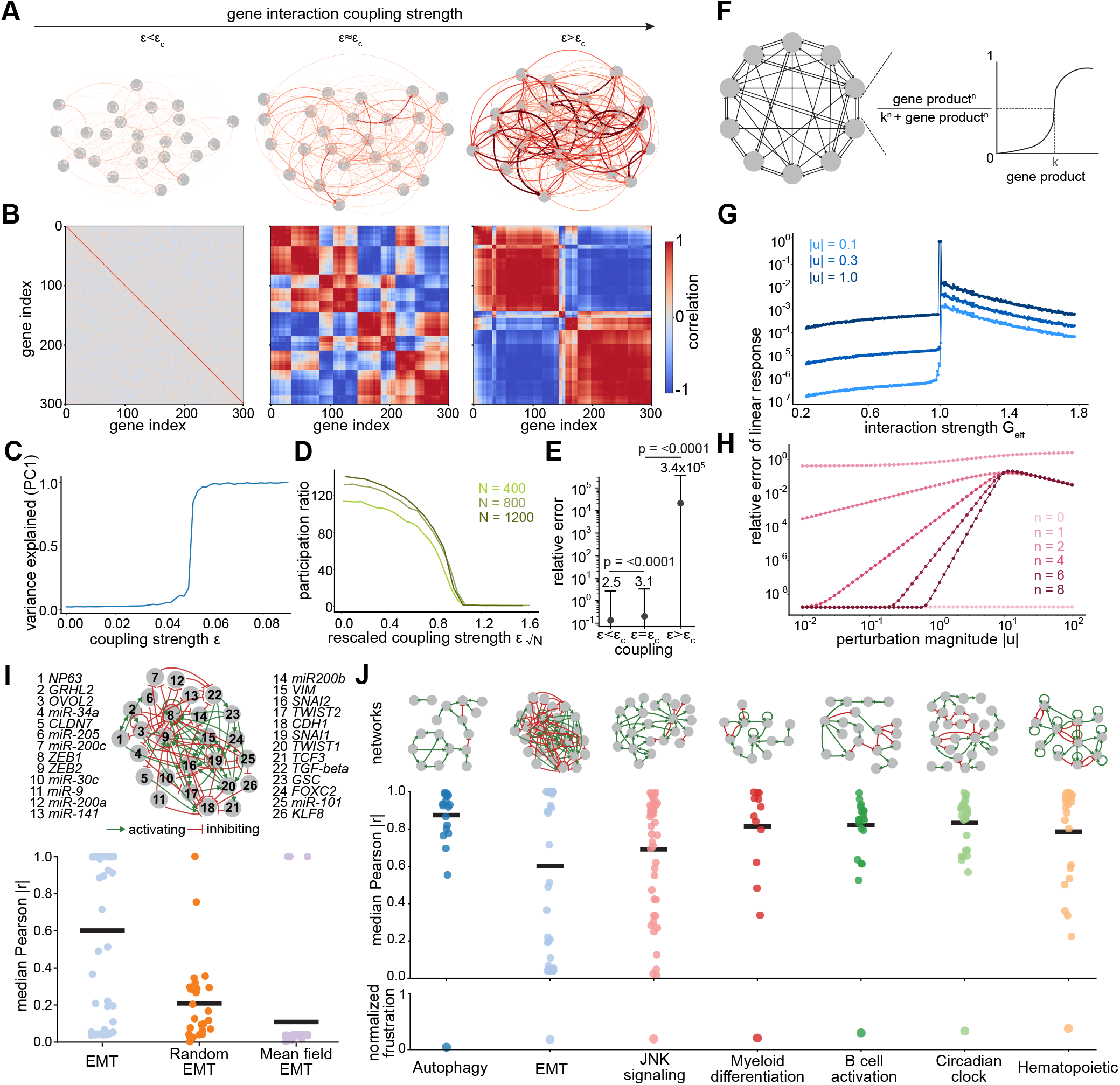
Genome-wide response to perturbations in synthetic regulatory networks. A) Random linear regulatory networks of N genes. Nodes represent genes and edge thickness represents the (absolute) strength of interactions between genes. Subcritical, critical and supercritical networks are shown from left to right. B) Steady-state gene-gene correlations for the networks in A). C) Fraction of variance explained by the first principal component as a function of the gene-gene interaction parameter. D) Participation ratio (effective dimension) as a function of the scaled interaction parameter for different sized networks (N = 400, 800, 1,200). E) Points are the typical error in linear response, ||Δ*X* − Σ*u*||^2^/||Δ*X*||^2^, across all genes in the three-regimes, averaged over 500 trajectories. Standard error bars shown. (One-sided Wilcoxon signed-rank tests: subcritical error < critical error, p-value < 0.0001 ; critical error < supercritical error, p-value < 0.0001). F) Non-linear networks with activating Hill function interactions, n = 10. G) Error in linear response as a function of effective interaction strength, G_eff_ at different perturbation magnitudes (|u| = 0.1, 0.3, 1.0). Points represent individual simulation runs and lines guide the eye. H) Error in linear response as a function of perturbation magnitude |u| for several different Hill coefficients (n), n = 10. I) EMT regulatory network (top), in which genes interact by inhibitory (red) and activating (green) edges. Bottom, median absolute Pearson correlation (|r|) between predicted and observed covariance under knockdown of each gene for the EMT network, a randomized version and a mean-field covariance for the EMT network. Each point is one gene; black bars indicate the absolute Pearson correlation mean across all genes in a network. J) Seven biological regulatory networks including EMT (top; edges coloured as in I), the median absolute Pearson correlation for knockdown of each gene (middle) and the normalized mean frustration from Boolean simulations (bottom). Each point in the middle row corresponds to one gene; black bars indicate the mean across all genes in a network.

We next tested CIPHER on a prototypical nonlinear dynamical gene expression system consisting of a network of Hill functions (Case 2; **Fig. 2F**; **Methods 2.2**). By perturbing a single gene in this nonlinear network, we could directly compare the observed response with our predictions. First, we fixed the Hill-function parameters and varied the gene-gene interaction parameter G, finding a sharp transition at G_eff_ = G/G* = 1 where G* is the critical interaction above which the trivial homogeneous state (zero expression for all genes) becomes unstable (see **Methods 2.2** for derivation and **Fig. S3A,B** for additional examples under different parameter choices) (**Fig. 2G**). Increasing the perturbation magnitude (|u|) corresponded to a subsequent increase in the relative mean squared error of linear response (||Δ*X* − Σ*u*|| ^2^ /||Δ*X*|| ^2^, **Fig. 2G and Fig. S3A,B**). We next fixed G and varied the Hill coefficient, n. As expected, CIPHER performed optimally under linear conditions (Hill coefficient, n=0), yielding the lowest error between actual and predicted values (**Fig. 2H**). However, as nonlinearity increased (Hill coefficient >1), error rates peaked during transition periods but remained low otherwise—a pattern consistent with the characteristic response curves of Hill function dynamics (**Fig. 2H**). We hypothesized that our framework’s prediction accuracy should decline under more challenging perturbation conditions. Specifically, we tested whether increasing either the perturbation magnitude (|u_i_|) for single gene knockouts or the total number of simultaneously perturbed genes would degrade performance. Indeed, the performance systematically decreased as we increased both the magnitude (|u_i_|) and total number of simultaneous perturbations (u_i_, where i ∈ {1,2,3,4,5}) (**Fig. 2F-H and S3C**), confirming that the method’s linear assumptions become limiting under extreme perturbation conditions. CIPHER also performed well for recently described “teams” topology implicated in cancer and developmental plasticity^41,42^(**Fig. S3I-K**).

### A theoretical framework explains why CIPHER works

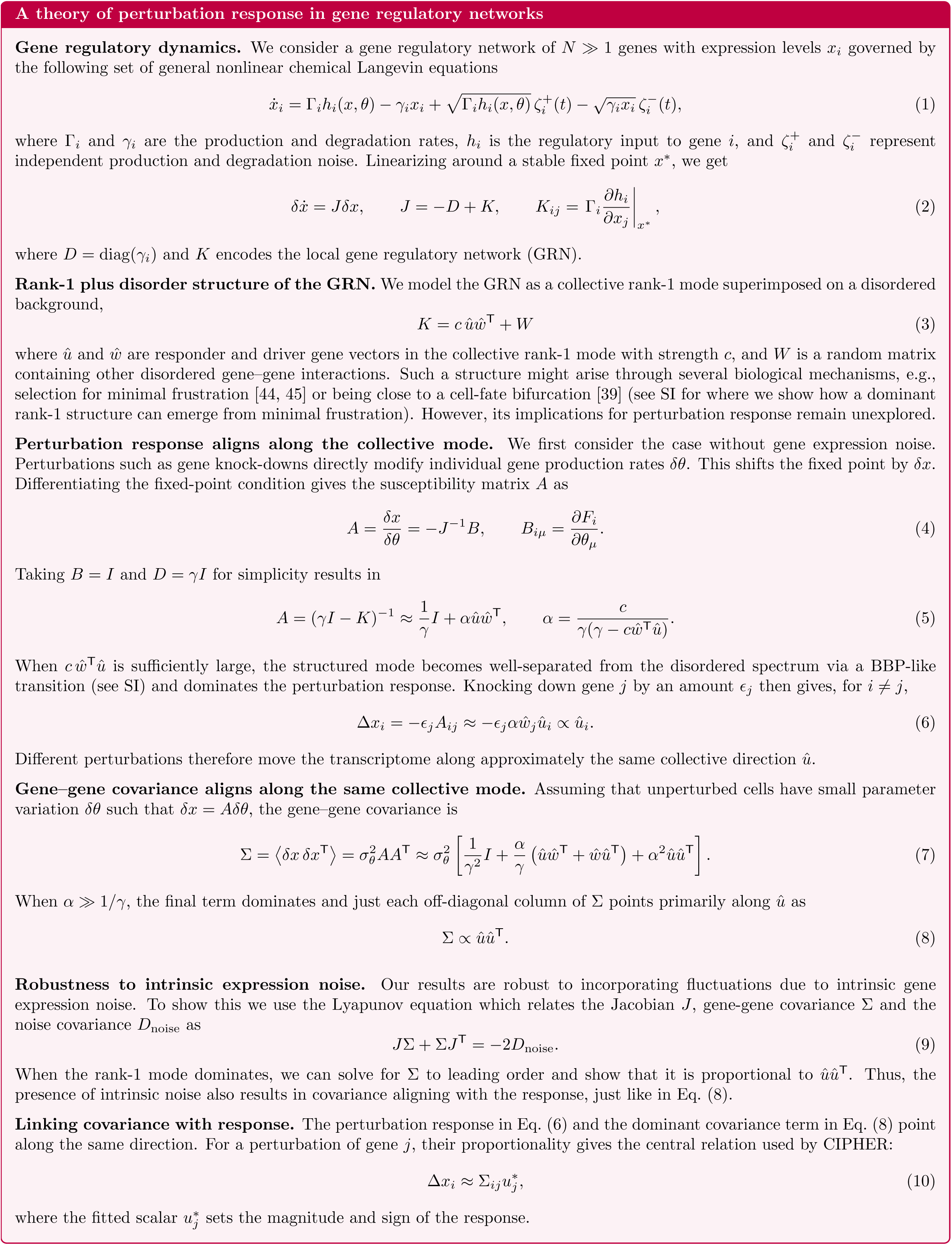

The success of CIPHER, even on toy networks, is relatively unexpected and raises a natural question: why should gene-gene covariances in unperturbed cells predict responses in noisy, nonlinear, and nonequilibrium gene regulatory networks? To address this question, we developed a theoretical framework in which gene-gene interactions contain a dominant rank-1 mode atop a heterogeneous, disordered background (**Box 1** and **Methods 1.1-1.8**). We found that when this mode is sufficiently strong, perturbations to different genes drive changes along a common dominant direction in the gene expression space (**Fig. S1**). This leads to gene-gene covariances aligning with perturbation responses. The theory suggests that a simple linear response relationship can emerge despite complex non-linear dynamics if the underlying gene regulatory networks are low-rank (**Fig. S2**). Recent work has suggested that such an approximately rank-1 structure could arise through several mechanisms, such as selection for minimal frustration^44,45^, and is a hallmark of several biologically-derived networks^44,45^ we investigate below. This theory makes several predictions (detailed in **Methods 1.9**), which we test throughout the rest of this study including in biologically-inspired reaction networks.

### Implementation on biologically-informed synthetic regulatory networks

We next evaluated CIPHER’s performance on biologically-informed regulatory networks, covering a range of network sizes and topology. These networks include: the epithelial-mesenchymal transition (EMT), autophagy, a JNK regulatory module, myeloid differentiation, B-cell activation, hematopoiesis, and the circadian clock^45–48^. We used a degree-preserving randomized EMT network as control. Each signed network was modeled using continuous ordinary differential equations in which activating and inhibitory interactions were represented by shifted Hill functions (See **Methods 2.3**). Rather than assigning a single fitted parameter set to each network, we used the random circuit perturbation (RACIPE)^49,50^ framework to generate ensembles of biologically plausible kinetic parameters, thereby incorporating heterogeneity in production rates, degradation rates, regulatory thresholds, Hill coefficients, and interaction strengths. In this framework each parameter realization can therefore be interpreted as a distinct cellular or environmental context, allowing performance to be evaluated across a broad distribution of dynamical regimes rather than for one finely tuned model. We first focused on the EMT network, as it is a well-established benchmark for multistable cell-fate regulation and has previously been shown to exhibit robust, low-dimensional structure across heterogeneous parameter ensembles (**Fig. 2I and Fig. S3F**). Prior work further demonstrated that the EMT network is minimally frustrated—its regulatory interactions are largely mutually consistent, so few nodes face conflicting inputs—and its dominant expression patterns and perturbation responses are preserved under substantial network simplification^44,45^. Moreover, it was shown that a single dominant mode (PC1) emerges in such low-frustration networks, consistent with a low rank structure (**Fig. S3D-H**). This network therefore provides a biologically realistic test case in which CIPHER should be highly predictive according to our theory. We quantified CIPHER’s predictions by computing the absolute Pearson correlation between the nonlinear response and the covariance-based linear-response prediction. CIPHER resulted in high Pearson correlations for the literature-informed EMT network, but the performance dropped significantly for randomized EMT network and real EMT network with scrambled gene-gene covariance matrix (mean-field) (**Fig. 2I**).

We next computed the absolute Pearson correlation for the remaining biologically-informed networks in **Fig. 2J**: the upper panel shows the distribution of perturbation-level correlations for each network together with network-level summaries, whereas the lower panel shows the corresponding normalized frustration value. Across all networks, CIPHER covariance structure predicted the direction of the non-linear perturbation response with high accuracy despite substantial kinetic-parameter heterogeneity (average Pearson 0.80) (**Fig. 2J**). Our theory explains that this is likely attributed to these networks having minimal frustration, which leads to a low-rank structure that is relatively robust to parameter heterogeneity (**Fig. 2J** and **Fig. S3D-H**). Together, these results demonstrate the ability of CIPHER in predicting responses for simulations of biologically-informed regulatory networks across contexts.

### Gene-gene correlations capture genome-wide perturbation responses in real datasets

Complex gene regulatory networks can be viewed as out-of-equilibrium systems with underlying correlations between genes. We tested whether CIPHER could recapitulate genome-wide responses to perturbations, which are typically a single gene knockdown or activation at a time. To address this question, we first systematically collected and performed quality control on existing single-cell perturbation datasets (see **Methods 3.4**). In total, we analyzed 27 high-throughput genetic perturbation datasets, containing 21 CRISPRi and 6 CRISPRa datasets and covering 18,075 perturbations, 6,078,966 cells, and 14 cell types (**Table 1**, **Fig. 3A**). We calculated covariance matrices for unperturbed single cells from each experimental gene count matrix (raw absolute counts for each gene with a minimum average expression cutoff). As a null model, we constructed a “mean-field” covariance by shuffling expression values across cells independently for each gene. This procedure preserves each gene’s marginal expression distribution but disrupts gene-gene covariances, thereby removing off-diagonal covariance structure while retaining gene-wise count statistics (**Fig. 3B**). We then implemented CIPHER on the forward problem, fitting the measured transcriptome-wide response with a single parameter *u*\* that rescales the covariance column of the true perturbed gene i. Because this scalar sets only magnitude and sign, the predicted direction is determined by the covariance alone, and we assessed how well it matched the data by calculating ΔR^2^ = 1-||Δ*X* − Σ*u*||^2^/||Δ*X*||^2^, Pearson correlation, and Spearman correlation values (**Fig. 3C-E**). A ΔR^2^ = 1 implies that we could perfectly transport the initial vector of gene expression < *X*>_0_ to the final measured target < *X*>*_u_*. We found that CIPHER alone was able to significantly recapitulate transcriptome-wide responses as compared to the mean-field model for both activating and interfering perturbations across multiple metrics including ΔR^2^, Pearson correlation, and Spearman correlation (**Fig. 3C-E**). We have illustrated gene-wise concordance between our predictions and experimental data for a few high performing test cases in **Fig. S4A,B**. While we systematically compare different transformation and normalization schemes in a later section, log1p transformation with appropriate change of variables did not affect the conclusions across all metrics (**Fig. S4F**). For perturbations with different representative ΔR^2^ and Pearson, we visualized the control to perturbed cell state transition with UMAP, and derive an effective CIPHER UMAP flowfield indicating the effects of perturbation on the control cell manifold, providing a direct comparison to the average perturbed control cell (**Fig. S6)**. We explored the mechanistic underpinnings of variability in ΔR^2^ values across datasets later in the results section. We note, however, that the per-gene values of ΔR^2^ had little dependence on the mean expression counts (**Fig. S4C**). To further test the dependence of perturbation response on gene expression level, we systematically removed the top and bottom 10%, 20%, and 30% of genes ranked by expression count (see **Methods 3.4, Fig. S5**). Mean Pearson correlation remained essentially unchanged from the full gene set across all six filters, for both CRISPRi and CRISPRa perturbations (**Fig. S5A,B**). Predictive performance therefore does not depend on any particular expression stratum, nor much on the most variable genes (**Fig. S7C**), but draws on covariance structure distributed across the full expression range. We also implemented CIPHER on recently proposed optical screens like Perturb-FISH^10^, with similarly high-confidence results (**Table 1 Binan25, Fig. S4D**). CIPHER also outperformed mean-field when using gene-gene correlations, but drops compared to covariance coordinates (**Fig. S7A,B**), which is expected from theory (See **Methods 1.9**).

**Fig. 3:**
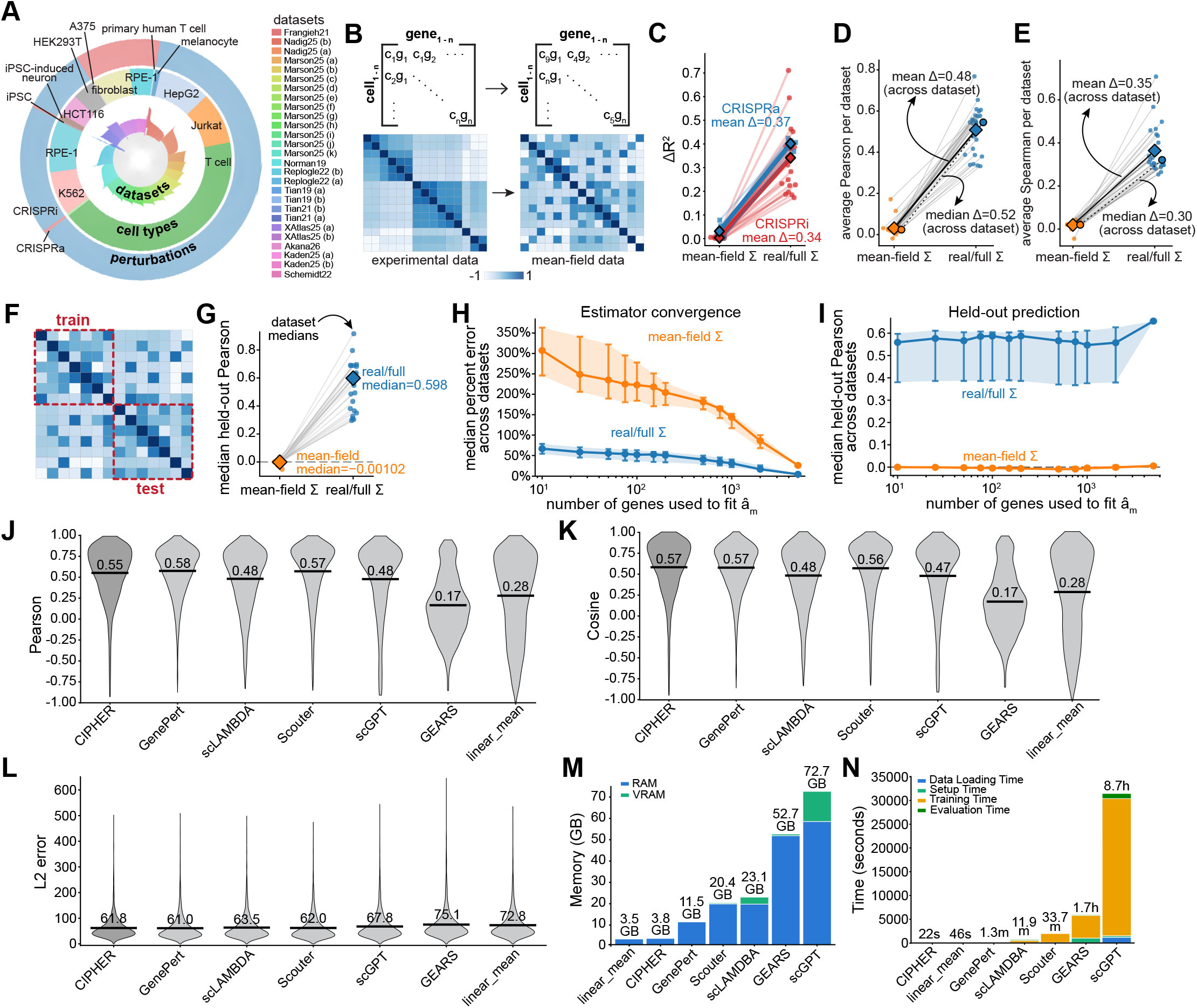
The forward problem: to what extent can covariance structure explain response to known gene perturbations from Perturb-seq. A) We consider 27 Perturb-seq datasets, comprising CRISPRi (21 datasets) and CRISPRa (6 datasets) gene perturbations (outermost ring) and a variety of different cell types (middle ring). The innermost ring shows the log-number of control (gray) and perturbed (colored) samples for each perturbation in each dataset. B) Experimental (real) and ‘mean-field’ (shuffled over cells, not genes) gene expression count matrices, and corresponding covariance matrices. C) Improvement in transcriptome-wide prediction accuracy obtained by using the experimentally measured covariance matrix instead of the mean-field covariance. Each line corresponds to one dataset and connects the performance using the mean-field covariance (left) to that obtained using the real covariance matrix (right). The vertical axis shows the increase in coefficient of determination (ΔR^2^). Red points correspond to CRISPRi datasets and blue points to CRISPRa datasets, with mean improvements indicated for each perturbation modality. Large diamonds and circles denote the mean and median across datasets. D) Average Pearson correlation between predicted and observed perturbation responses for each dataset using the mean-field covariance (left) and the real covariance (right). Each connected pair represents one dataset. Large diamonds denote the mean across datasets, while annotations indicate the mean and median improvements. E) Same analysis as in (D), but using Spearman correlation as the performance metric. Individual datasets are connected by gray lines, with mean values shown as diamonds and overall mean and median improvements annotated. F) Schematic of the train-test evaluation used to assess generalization across included genes. A block of the covariance matrix including the perturbed gene is estimated, while prediction accuracy is evaluated on held-out test genes (lower-right block), as indicated by the dashed partition. G) Median held-out Pearson correlation obtained using covariance matrices estimated from the training subset consisting of 50% of genes selected randomly. Each point corresponds to one dataset, comparing predictions using the mean-field covariance and the real covariance. Dataset medians are indicated by diamonds. H) Dependence of fitting parameter error â_m_ on the number of genes used to estimate it. The horizontal axis denotes the number of genes included during estimation, while the vertical axis shows the median percent error across datasets. Blue curves correspond to real covariance matrices and orange curves correspond to mean-field covariance estimates. Error bars denote variability across datasets. I) Held-out prediction performance as a function of the number of genes used to estimate the response fitting parameter â_m_. Median Pearson correlation across datasets is plotted for real covariance matrices (blue) and mean-field covariance matrices (orange), with error bars indicating variation across datasets. J) Distribution of Pearson correlations between predicted and experimentally observed perturbation responses for CIPHER and other state-of-the-art methods. Violin plots summarize the distribution across perturbations, horizontal bars indicate medians and median values are labeled numerically above each distribution. K) Same comparison as in (J), using cosine similarity between predicted and observed perturbation responses. Median values are indicated for each method. L) Distribution of transcriptome-wide L2 prediction error for each method. Violin plots summarize the distribution across perturbations, with horizontal bars indicating median errors and numerical median values shown above each violin. M) Peak memory requirements during model execution. Blue bars indicate peak RAM usage and green bars indicate peak GPU memory (VRAM) usage for each method. Memory consumption is reported in gigabytes (GB). N) Runtime comparison across methods. Stacked bars partition total execution time into data loading (blue), setup (green), training (orange), and evaluation components (dark green). Total runtimes are annotated above each bar, with times reported in seconds, minutes, or hours as appropriate.

### A single scaling factor predicts responses of held-out genes

In constructing CIPHER, we found that the fitting parameter remains constant for every gene. We reasoned that because a single scaling factor applies to every gene, a value fit on one subset of genes should predict the responses of genes never used in the fit. This gives a direct test: hold genes out of the fitting procedure, then compare their predicted responses against ground truth. To test this idea, we evaluated held-out-gene prediction, in which the scaling factor was fit on a subset of genes and the performance was evaluated on genes not used for fitting **(Fig. 3F)**. Across all datasets, CIPHER produced high Pearson correlation values for held-out-genes, a performance that fell to zero for the mean-field covariance control (**Fig. 3G).** We further examined how many genes were required to estimate the scalar perturbation amplitude. Across all datasets, the real covariance model showed low amplitude-estimation error (**Fig. 3H**) and high held-out Pearson correlation (**Fig. 3I**) across a broad range of training-genes, both in terms of raw number and identity of training genes, whereas the mean-field model remained near zero in held-out performance (orange curve in **Fig. 3I**). Thus, CIPHER’s performance is not an artifact of fitting on the full response vector, but persists when evaluated on held-out genes.

### Benchmarking CIPHER against other methods

We benchmarked against five recent approaches spanning distinct methodological classes—GenePert^51^, scLAMBDA^21^, Scouter^52^, scGPT^20^, and GEARS^53^—as well as a linear mean baseline, motivated by recent concerns that simple linear models can match far more complex architectures on this task^54^. To evaluate performance broadly, we assembled 27 single-gene perturbation datasets covering 14 cell types (Jurkat, melanoma, primary T cells, CD4+ T cells, T cell leukemia, HepG2, HCT116, K562, HEK293T, fibroblasts, retinal pigment epithelial cells, iPSC, and iPSC-derived neurons), with 26 to 1,000 unique perturbations (after quality control steps, see **Methods 3.4 and 3.5; Fig. S8A**) per dataset—expanded from the 17 datasets used in the most recent systematic benchmark study^55^. We scored each method on six metrics capturing complementary aspects of prediction quality^55–57^: Pearson correlation, Spearman correlation, cosine similarity, mean squared error, L2 error, centroid accuracy (Systema^57^), alongside training/evaluation time and compute cost (**Fig. 3J-N; S8**). For each dataset, we split perturbations 50/50 into training and test sets, fitting each method on the training half and evaluating on held-out in-distribution test perturbations. To ensure a fair comparison across methods, we applied appropriate changes of variables so that all methods were evaluated on metrics in a common expression space (see **Methods 3.5**).

Across all datasets, CIPHER performed at or near the top on the majority of metrics (**Fig. 3J-L; S8E-G**). It achieved the highest cosine similarity (0.57) and lowest mean squared error (1.91) of any method tested, and the second-lowest L2 error (61.8), marginally behind GenePert (**Fig. 3K,L; S8G**). On Pearson correlation, CIPHER (0.55) was competitive with the leading methods GenePert (0.57) and Scouter (0.57), and exceeded scLAMBDA and scGPT (**Fig. 3J**). On centroid accuracy, a Systema metric designed to isolate perturbation-specific effects from systematic variation, CIPHER (0.71) again ranked among the leading methods, just behind Scouter (0.77) and well above GEARS (0.54) and the linear mean baseline (0.50) (**Fig. S8F**). The linear mean baseline in turn outperformed GEARS, consistent with recent reports that added architectural complexity does not reliably improve prediction^54^. CIPHER was also the most efficient method by a wide margin, completing in 22 seconds per dataset using 3.8 GB of memory, compared with 1.7 hours and 52.7 GB for GEARS and 8.7 hours and 72.7 GB for scGPT (**Fig. 3M,N; S8B,C**). This gap predominantly reflects the absence of a training phase: whereas the deep learning methods spend the bulk of their runtime on parameter optimization, CIPHER fits only a single scaling parameter and requires no GPU. Indeed, the deep learning methods use roughly 10⁶-10⁷ parameters, compared with CIPHER’s single scalar (**Fig. S8D**).

### Covariances are preserved across studies of same cell type

We tested whether the gene-gene co-fluctuations in unperturbed cells from one dataset or experiment can be informative in fitting perturbation response in another independent study of the same cell type or cell line. If so, it would suggest a universal underlying structure of covariance that carries essential information for perturbation response. Conversely, gene-gene covariances in each experiment—regardless of whether cell type is shared—are sensitive to specific experimental conditions. To test this idea, we focused on three unrelated datasets with a shared cell type (neuron) (**Table 1**, **Fig. 3A**). We calculated covariances from two of the datasets and used CIPHER to fit responses in the third “host” dataset (Tian21_a, **Fig. 4A**). Remarkably, we found strong correlation (ΔR^2^=0.73) between transcriptome-wide responses calculated from cross-dataset covariance matrices of the same cell type (**Fig. 4B**) and typically much poorer performance from cross-dataset of a different cell type (**Fig. 4C; Fig. S4E)**. Similar analysis across distinct cell types/lines failed to recapitulate transcriptome-wide responses (**Fig. 4C**). However, real covariances of unrelated cell types outperformed their corresponding mean-field models (shuffling the gene counts column-wise, which retains the original mean and distribution for each gene individually), suggesting the presence of certain universal, albeit weakly-contributing, fluctuation structures that inform outcomes (**Fig. 4C**). To test whether cross-experiment covariance matrices retain predictive ability in primary tissue beyond the single-cell perturbation datasets, we analyzed three single-cell atlases from CellxGene^58^: human embryonic meninges (5-13 weeks post conception, 58 cell types), human intestine organoid (22 cell types), and human lung organoid (9 cell types)^59,60^. To compute the covariance matrices, we used as many cells as there were neurons in the meninges dataset. The meninges dataset contained many neuron-related cell types, while lung and intestine datasets both predominantly contained non-neuron cell types. Most cell types in the embryonic meninges dataset better recapitulated cell culture perturbation responses than their intestine and lung counterparts, with the only neuron cell type in the intestine dataset performing best among all intestine cell types (**Fig. 4D; Fig. S4G-I**). We further tested these findings for other datasets including primary T Cells and Retinal pigment epithelium (RPE) cell line (**Fig. 4E,F**). In each case, the performance did not drop significantly even when we used covariance from a different dataset to compute responses. We note, however, that the performance dropped when we broke the covariance with the mean-field count matrices (**Fig. 4E,F**).

**Fig. 4:**
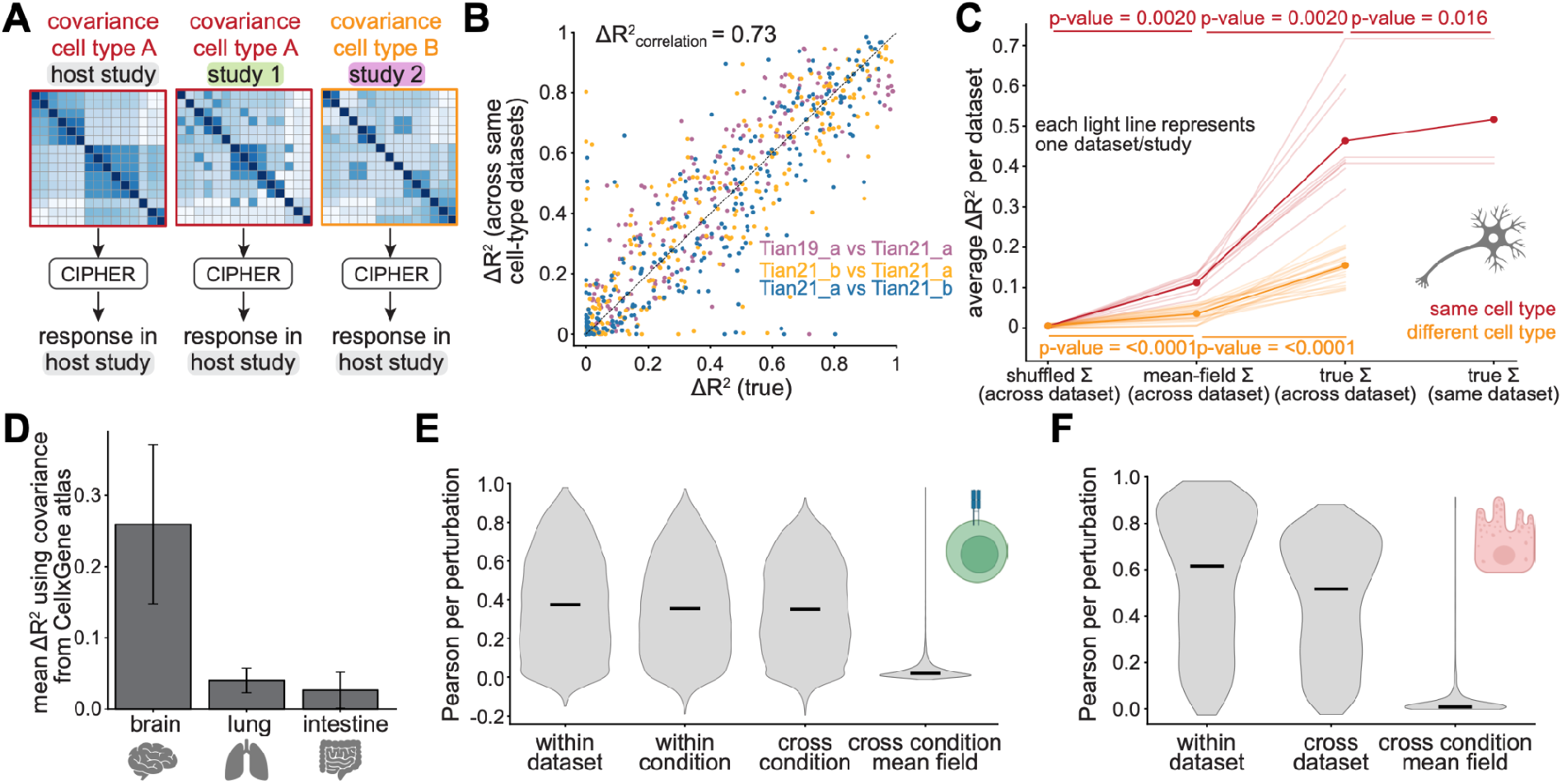
CIPHER transfers across datasets and cell atlases. A) Responses in a host study from inter-study covariances either of the same or different cell type. B) ÄR^2^ values across neuron datasets. ΔR^2^ values using the true covariance matrix from the host study compared to those from a different dataset but same cell type (ΔR^2^_correlation = 0.73). Each point is a perturbed gene. C) Average ΔR^2^ values across all pairs of datasets and conditions, using a covariance matrix from either a shuffled X_0_ (across datasets), the mean-field approximation (across datasets), the true covariance (true Σ, across datasets), or the true covariance from the same dataset. Thin lines correspond to individual dataset pairs (averages over all perturbations in the host dataset with a Σ of the same cell type in red or different cell types in orange) and points are averages over the typical ΔR^2^ values for these pairs of datasets. Note that there are only three lines connecting the across and within dataset real covariance for the same cell type. This is because we are comparing the three datasets shown in B, and that there is no true covariance from the same dataset and different cell type. (Same cell type Shuffled vs. mean-field: Wilcoxon Signed-Rank Test, p-value = 0.0020; Same cell type mean-field vs. cross-dataset covariance: p-value = 0.0020; Same cell type cross-dataset covariance vs. same dataset covariance: p-value = 0.016. Different cell type Shuffled vs. mean-field: p-value < 0.0001; Different cell type mean-field vs. true: p-value < 0.0001). D) Mean improvement in prediction accuracy (ΔR^2^) obtained by transferring covariance matrices derived from the CellxGene atlas to external perturbation datasets. Bars show the average increase in ΔR^2^ relative to predictions using the corresponding mean-field covariance for brain, lung, and intestine datasets. Error bars denote variability across cell-types within each tissue. E) Distribution of Pearson correlations between predicted and real perturbation responses for T-cell perturbation datasets under different covariance transfer settings. Violin plots summarize prediction accuracy using covariance matrices estimated within the same dataset, within the same experimental condition, across different experimental conditions, and using a cross-condition mean-field covariance. Horizontal black bars indicate the mean Pearson correlation across perturbations. F) Distribution of Pearson correlations between predicted and real perturbation responses for RPE-1 perturbation datasets under different covariance transfer settings. Violin plots compare prediction accuracy using covariance matrices estimated within the same dataset, transferred across datasets (and labs) of the same cell-line, and using a cross-dataset mean-field covariance. Horizontal black bars indicate the mean Pearson correlation across perturbations.

Collectively, these results suggest the presence of cell-type specific strongly-contributing and universal weakly-contributing latent fluctuation structures which can be harnessed to predict transcriptome-wide responses.

### Predictions of true driver perturbations from genetic screens

Having tested CIPHER in the forward direction on capturing genome-wide response to single perturbations, we next investigated the inverse problem: given a covariance matrix and a transcriptome-wide change in gene expression, what gene (or genes) most likely induced the change (**Fig. 5A**)? To address this question, we solved for a full combinatorial gene perturbation *u* (a potentially nonzero element for each gene). We argued that ordering the resulting *u_i_* by magnitude should, in principle, provide an effective ranking of *gene_i_* and a point estimate of whether it was likely knocked-down or activated. High-throughput Perturb-seq data provides an excellent platform to solve this inverse problem since the ground-truth perturbations are known *a priori*. To incorporate the possibility of error in the estimates of the averages, covariances, or deviations from linearity, we framed the question as a Bayesian linear regression problem for *u*: Δ*X* = Σ*u* + ɛ, where ɛ is a normally distributed random variable (**Fig. 5B**). Concretely, we sample from the posterior distribution *p*(*u*|Δ*X*, Σ) assuming a Gaussian prior on perturbations which shrinks effect sizes. We employ a Central Limit Theorem informed noise metric (**Methods 5, Eq. S138-S139)**, which outperformed isotropic Gaussian noise both on null perturbation tests and in picking the top-k inferred true perturbations (**Fig. S9A,B)**. This framework is analytically tractable and enables us to put error bars on estimates of perturbation effects, and assign each gene a posterior inclusion probability (PIP) that it has true nonzero effect (**Fig. 5B**). Together, the Bayesian approach allowed CIPHER to provide uncertainty-aware estimates of the perturbation effect sizes that the naive point estimates like *u* = Σ ^−1^ could not. We also performed power analysis using the Central Limit Theorem and from linear response theory arguments, finding that typically perturbations require ^2 4^ cells to accurately estimate the average perturbed mean (**Fig. S9H, Methods 5**).

**Fig. 5:**
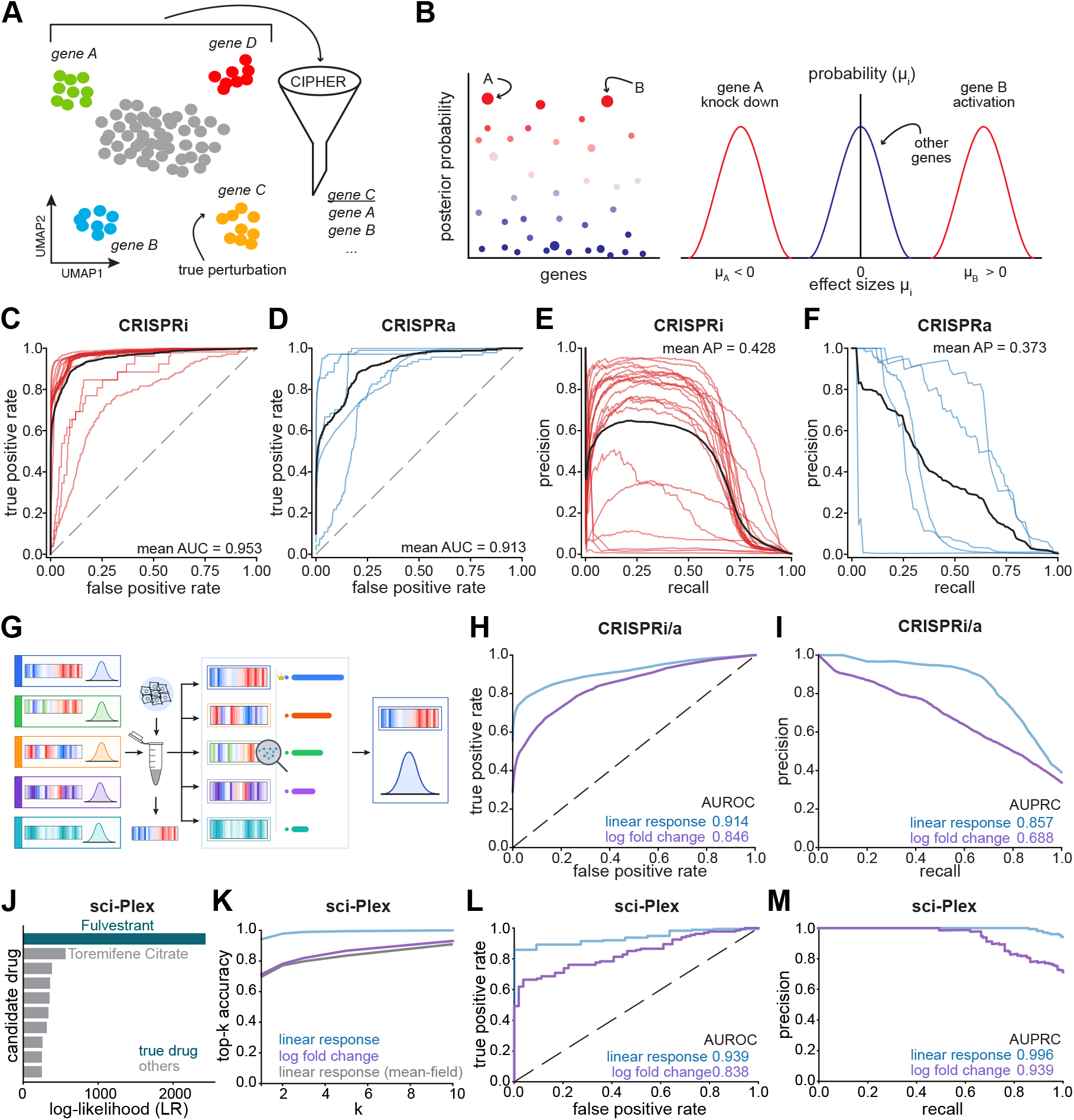
The inverse problem: predicting causal drivers of perturbation response in transcriptome-wide screens. A) Schematic depicting how CIPHER ranks genes responsible for a measured perturbation using knowledge of the control-cell gene expression fluctuations. Each point corresponds to a transcriptome from a single cell and is colored by the perturbed gene. B) Schematic of example data and interpretation. Left: Posterior probability of a nonzero perturbation effect across genes (points). True perturbed genes A and B have high posterior probabilities. Right: Effect size distributions for different perturbations. Distributions of effect size for genes A and B as well as a composite distribution over other unperturbed genes. C) Receiver-operator characteristic curves for each CRISPRi dataset (colored curves, mean AUC = 0.953). Black curve indicates average across datasets. D) Receiver-operator characteristic curves for each CRISPRa dataset (colored curves, mean AUC = 0.913). Black curve indicates average across datasets. E) Precision-recall curves for each CRISPRi dataset (colored curves, mean AP = 0.428). Black curve indicates average across datasets. F) Precision-recall curves for each CRISPRa dataset (colored curves, mean AP = 0.373). Black curve indicates average across datasets. G) Schematic of the likelihood-based CIPHER inverse ranking of transcriptional perturbation signatures. CIPHER identifies that the true signature is blue. H) Receiver-operator characteristic curves composite for all perturbations across CRISPR datasets for CIPHER (blue, linear response; 0.914) and log-fold-change ranking (purple, 0.846). I) Precision-recall curves for CIPHER (blue, 0.857) and log-fold-change ranking (purple, 0.688) across CRISPR datasets. J) CIPHER log-likelihood ranking for one held-out drug in the sci-Plex^61^ experiment. True compound is shown in blue. K) Top-k accuracy for identifying the true sci-Plex^61^ compound from heldout cells with CIPHER (blue) and log-fold-change ranking (purple). Mean-field linear response is shown in gray. L) Receiver-operator characteristic curves composite for all perturbations across sci-Plex^61^ for CIPHER (blue, 0.939) and log-fold-change ranking (purple, 0.838). M) Precision-recall curves for CIPHER (blue, 0.996) and log-fold-change ranking (purple, 0.939) across sci-Plex^61^ perturbations.

For each dataset, we compared the ranking metrics (PIP,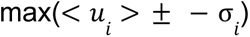) with receiver-operator characteristic (ROC) curves, finding that ranking the perturbations by the maximum spread 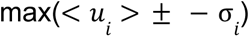 proved most informative of the true perturbation (Fig. 5C-F). We found that regardless of the dataset, cell type, or perturbation type (CRISPRi or CRISPRa), our approach, which relies on gene-gene interactions, achieved high values on receiver-operator characteristic curves (CRISPRi mean AUC = 0.953 vs CRISPRa mean AUC = 0.913) (**Fig. 5C,D**) as well as precision-recall analysis (CRISPRi mean AP = 0.428; CRISPRa mean AP = 0.373) (**Fig. 5E,F**). CIPHER also exhibited high performance for double perturbation datasets (**Table 2**; **Fig. S9C,D**). Ranking the true perturbation across the full gene set, the median rank of the true target was 2 and it was ranked first in 47% of cases (**Fig. S9E-G**).

**Table 2:**
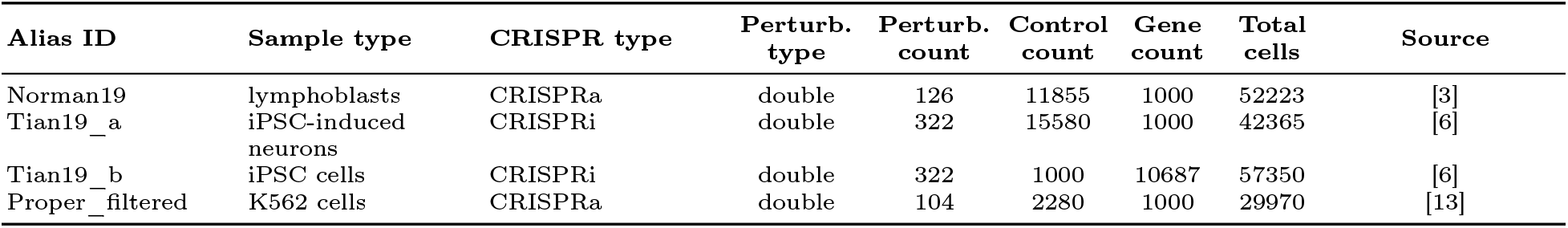
Specifications for the double-perturbation of the single-cell peturbation datasets used in subset.

We extended our analysis from target-gene identification to perturbation-identity classification. In this setting, transcriptional profiles were split into training and testing sets (**Methods 5.1**). A response signature was inferred for each candidate perturbation in the testing set, and held-out test responses were scored against all candidate signatures using the covariance-aware linear-response likelihood (**Fig. 5G**). We compared covariance-aware linear-response scoring with baselines on log-fold-change similarity. In the held-out perturbation identity recovery, linear-response scoring was evaluated by ROC and precision-recall curves alongside this log-fold-change baseline (**Fig. 5H,I**). CIPHER returned values higher than log-fold change for all metrics (AUROCCIPHER = 0.914, AUROCLFC = 0.846; AUPRCCIPHER = 0.857, AUPRCLFC = 0.688) (**Fig. 5H,I**).

### Predictions of true driver drug perturbations from pharmaceutical screens

We tested our framework’s ability to identify causal perturbations from massively multiplexed single-cell chemical screens, namely sci-Plex^61^ and Tahoe-100M^62^. We first focused on the 188 pharmaceutical perturbations from the sci-Plex3 dataset, splitting the cells into a training and testing set (see **Fig. 5G, Methods 5.1**). We next fit the transcriptional perturbation signatures *u*_d_ for each drug perturbation, using Δ*X*, Σ, and *H* . We subsequently evaluated the likelihood in Eq. 4 for each drug with Xtest, and ranked the drugs by their likelihoods under the linear response model. For comparisons, we also evaluated the mean-field linear response model, as well as classified drugs by the cosine similarity of the log-fold changes, between control and perturbation, within training and test sets. CIPHER-based classification scored highly both on top-k accuracy and AUROC/AUPRC metrics (AUROCCIPHER = 0.939, AUROCLFC = 0.838; AUPRCCIPHER = 0.996, AUPRCLFC = 0.939) (**Fig. 5J-M**). We next implemented CIPHER on the recently reported Tahoe-100M dataset where we performed a more stringent test: we took two experimental replicates (plates 6 and 14), trained on one and validated on the other.

CIPHER-based classification again scored highly both on top-k accuracy (**Fig. S10A,C)** and AUROC/AUPRC metrics for within-plate (**Fig. S10B**: AUROCCIPHER = 0.850, AUROCLFC = 0.615; AUPRCCIPHER = 0.877, AUPRCLFC = 0.492) and across-plates (**Fig. S10D**: AUROCCIPHER = 0.774, AUROCLFC = 0.756; AUPRCCIPHER = 0.714, AUPRCLFC = 0.619) **(Fig. S10B,D).** These results further point to the importance of covariance structure in identifying causal pharmaceutical perturbations in chemical compound screens.

To determine whether the inferred dense intervention vectors captured shared mechanisms of drug action, we hierarchically clustered the drug-level CIPHER perturbation profiles u* from the sci-Plex screen across both compounds and genes using correlation distance and average linkage (**Methods 6.2 and Fig. S11A**). The resulting heatmap revealed several coherent mechanism-associated blocks, including a prominent cluster of 17 HDAC inhibitors, as well as contiguous groups of JAK/STAT inhibitors, RTK/multikinase inhibitors, DNA-damaging or antimetabolite compounds, and Aurora/cell-cycle inhibitors. We quantified this organization using a clustering score defined as the excess of between-class over within-class mean (cophenetic distance), with a permutation null generated by shuffling drug-class labels while preserving category sizes. The observed global clustering score was substantially above the null (permutation z-score = 12.71, empirical p-value < 0.0001), with significant category-specific clustering for HDAC, RTK/multikinase, Aurora/cell-cycle, apoptosis/BCL2, DNA-damage/antimetabolite, and JAK/STAT compounds (**Fig. S11B,C**). Same analysis resolved by cell-type revealed additional cell-type specific compound clustering (e.g. hormone-specific compounds for MCF7, a hormone-dependent breast cancer cell line) (**Fig. S11D-F**). Together, these results indicate that CIPHER perturbation profiles recover reproducible multigene intervention programs that organize compounds according to a broad mechanism of action.

### Identifying regulatory modes driving genome-wide responses to perturbations

Our theoretical framework explains that the success of gene-gene co-fluctuations in capturing transcriptome-wide responses might arise from a low-rank structure of the underlying gene regulatory networks. This low rank structure may be interpreted in terms of a few dominant regulatory modes. To find evidence of these modes, we decomposed the covariance matrix into its eigen-representation, where eigengenes constitute effective regulatory modules: combinations of multiple co-expressed genes, each capturing a different amount of variance (**Methods 7**). Unlike the additive co-expression programs enumerated by dedicated program-discovery methods such as cNMF^63^ or Spectra^64^, these covariance eigenmodes define the collective directions along which perturbation response is transduced, and are used directly in the forward and inverse problems. Given our theory, we expect that only one or a few modes should capture most of the variance. We indeed found evidence consistent with this expectation across covariance matrices collected from multiple datasets (**Fig 6A,B; Fig. S12A**).

**Fig. 6:**
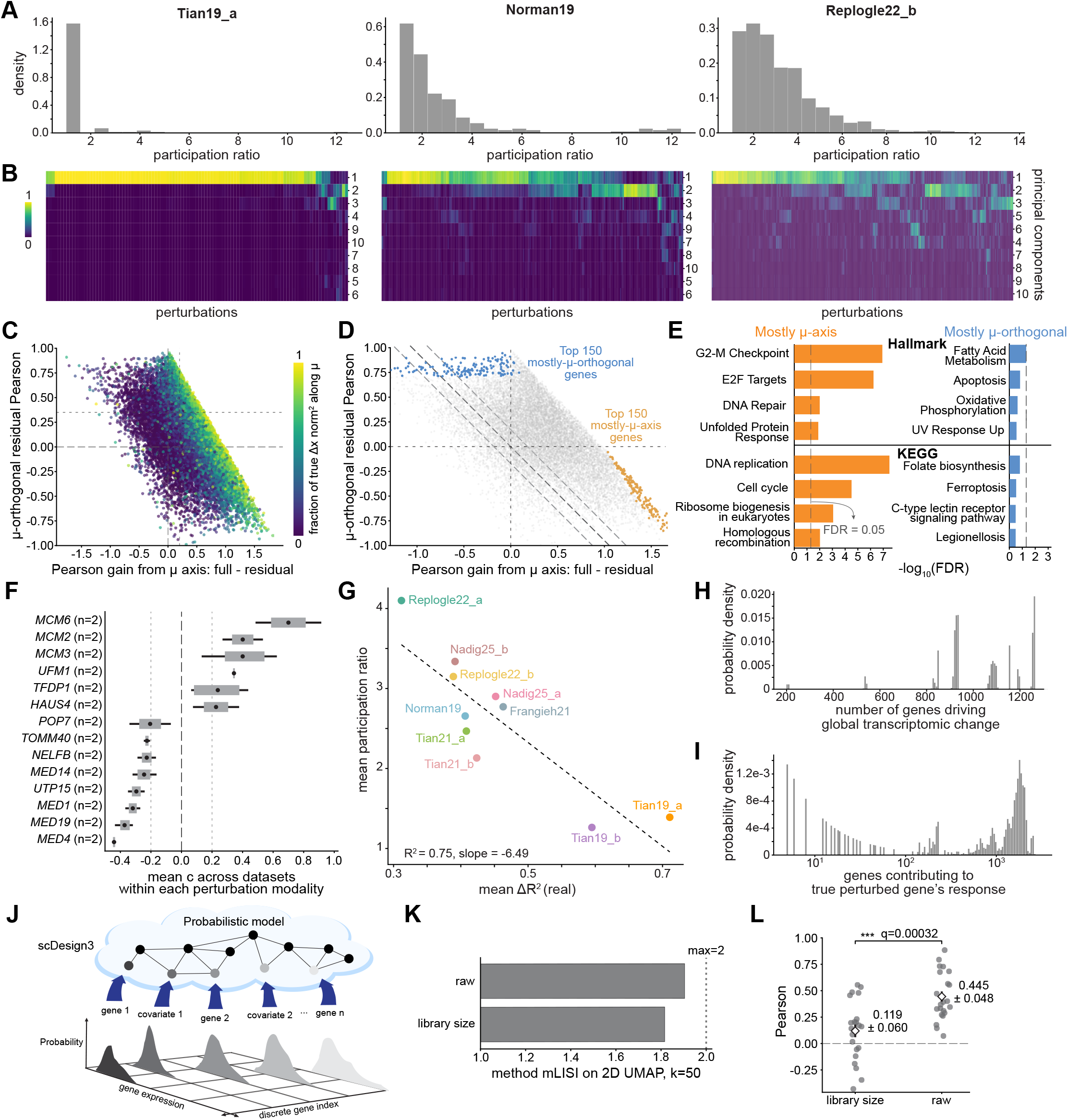
The effective dimensions of transcriptome-wide response. A) Distributions of participation ratio (i.e. effective dimension) across all perturbations for three Perturb-seq datasets. B) Clustered heatmaps of the fraction of the response that falls along each principal component for the datasets in A). C) Relationship between prediction improvements arising from transcriptome-wide (μ-axis) and orthogonal covariance modes. Each point represents a perturbation. The horizontal axis shows the increase in Pearson correlation attributable to the dominant response direction (μ-axis), while the vertical axis shows the residual improvement arising from covariance components orthogonal to this global μ direction. Points are colored by the fraction of the true response magnitude lying along the dominant response direction. Gray lines mark zero (long dashes) and threshold reference values on each axis (fine dashes). D) Same scatter as in (C), highlighting perturbations with the largest gains from different covariance components. Orange points denote the 150 perturbations with the largest improvement attributable to the dominant response direction (μ-axis), while blue points denote the 150 perturbations with the largest improvement attributable to orthogonal covariance modes. Gray points indicate all remaining perturbations. Dashed diagonal reference lines indicate constant total improvement. E) Gene set enrichment analysis of genes associated with the dominant response direction and orthogonal covariance modes. Left: Hallmark and KEGG pathways enriched among genes contributing primarily to the dominant response direction. Right: Hallmark and KEGG pathways enriched among genes contributing primarily to orthogonal covariance modes. Bar lengths indicate enrichment significance expressed as −log10(FDR). Dashed line represents FDR = 0.05. F) Mean standardized global growth scaling factor c = 1 - dX/mu. Horizontal bars show the average contribution of each gene across datasets, dots represent the mean, with error bars indicating variability across datasets modalities. The black dashed line marks c = 0 (no net change along μ); the gray dotted lines mark c = ±0.2, the threshold used to select reproducible perturbations. Each gene was identified in n = 2 datasets, as indicated in parentheses. G) Mean participation ratios vs. mean ΔR^2^ values over the CRISPRi/a Perturb-seq datasets considered. Each point corresponds to a dataset. Dashed line represents the best fit line describing the linear trend. H) The effective number of genes driving global change, calculated from the entropy of the fractional contributions of each gene’s change from every other gene. I) The effective number of genes impacting the expression change of the true perturbed gene across datasets, calculated in the same way as H). J) Schematic illustrating the scDesign3 probabilistic generative modeling of single-cell transcriptomes. A latent probabilistic model generates gene expression while incorporating marginals over measured genes and covariates, producing a distribution over gene expression values that can be sampled to generate synthetic single-cell data. K) Median mixing score (mLISI) computed on two-dimensional UMAP embeddings (k=50) comparing synthetic datasets generated using raw counts and library-size-normalized counts. The dashed vertical line denotes the maximum possible mLISI value of 2. L) Comparison of transcriptome-wide prediction accuracy obtained using raw counts and library-size-normalized counts. Each point corresponds to one perturbed gene in the Replogle22_a (RPE-1) dataset, with the overlaid diamond representing the mean and the error bar representing the standard error for each normalization. The adjusted q-value=0.00032, from two-sided paired Student’s *t*-tests across perturbations, with p-values adjusted for multiple comparisons using the Holm correction.

An expectation of our theory is that the response to perturbations should be effectively low-dimensional and dominated only by a few eigengenes (**Box 1**, **Methods 1**). To test this, we calculated a participation ratio for each perturbation in each dataset. The participation ratio—a measure of how many eigengenes actively contribute to the response—of a vector is a measure of how many nonzero elements it has: it evaluates to 1 when there is one nonzero and to the length of the vector when all elements have equal size. Specifically, we evaluated the participation ratio of the squared overlap (dot product) of each eigengene with the total response vector Δ*X*, yielding an estimate of the effective dimensionality of response. Indeed, as expected, the participation ratio was generally low relative to the dimension of the datasets (mean = 3.07 eigengenes over all perturbations) with notable variation between datasets (**Fig. 6A and S12B,E,F**). Interestingly, a majority of responses in Tian19_a (**Table 1**)—which exhibited the highest ΔR^2^ values—could essentially be described with a single dimension, consistent with theoretical arguments (**Box 1**). Conversely, datasets with relatively high participation ratios correspondingly exhibited lower ΔR^2^ values (correlation ΔR^2^ = 0.75; slope= -6.5) (**Fig. 6G**). We observed consistent results when comparing the normalized contribution of each eigengene for each perturbation, again illustrating that Tian19_a responses were typically well described by their first principal component, unlike the other datasets (**Fig. 6B; Fig. S12F**).

Next, we checked whether certain gene programs underlie dominant regulatory modes by calculating the dominant genes from the eigengene loadings for each principal component, and performed a gene ontology^65^ search on them (**Methods 7.2**). In line with a recent study on covariances in mammalian cells^66^, the majority of processes are related to translation and general cellular homeostasis (**Fig. S12C,G,H**), a trend that holds true over all datasets combined as well as within datasets from a single cell type (Neuron, **Fig. S12G,H**). We wondered whether this reflects a general principle of perturbation response or simply the baseline variation characteristic of growth, translation and homeostasis. We explore this in two ways: by examining the subleading covariance modes below, and by decomposing the response into growth-aligned and pathway-specific components in the next section.

While the dominant modes reflect translation and homeostasis shared across cell types, the subleading modes capture cell-type-specific programs. Beginning with the subleading modes, we examined the second and third principal components across five cell types. We found that each PC was dominated by a coherent, recognizable identity or state program (**Fig. S14D-H**). HepG2 cells showed a hepatocyte secretory program marked by ALB and a suite of apolipoproteins (*APOA1*, *APOB*, *APOE*). Activated T cells were dominated by an effector cytokine program led by *IFNG*, with *CCL4*, *GZMB*, and *IL2*. Neurons carried a neuronal identity program marked by *NEFM*, *NEFL*, and *VGF,* while fibroblasts showed a matrix and contractile state dominated by IGFBP5, and melanoma an inflammatory remodeling state marked by TIMP1 and the S100 family. Beyond the global modes that drive genome-wide response, covariance structure therefore also encodes rich, cell-type-specific regulatory organization.

### Perturbation responses separate into global growth and pathway-specific regimes

To determine the dependence of CIPHER predictability on a global expression mode, we decomposed both the observed and predicted response into components parallel and orthogonal to the control mean-expression vector in the control state, μ. The “μ-parallel” component captures coordinated scaling of the baseline transcriptome and can be interpreted as a global growth, biomass, or transcriptional-output axis. We note that such information is lost when the expression data are normalized, consistent with our theoretical arguments (**Methods 8**). We plotted the gain in Pearson correlation contributed by this axis against the correlation retained after removing it (**Fig. 6C**). Perturbations fell along a continuum, ranging from those whose apparent predictability was dominated by global expression scaling to those whose response remained strongly predictable within the μ-orthogonal subspace.

We next took the perturbations at the two extremes of this continuum (150 from each) and asked whether they were functionally distinct, performing pathway enrichment analysis using three complementary gene set collections: Hallmark, KEGG, and Reactome (**Fig. 6D,E; Fig. S13A**). Perturbations dominated by the μ-parallel component were strongly enriched for central growth programs, including E2F targets (controlling the G1-to-S transition) and G2-M checkpoint activity, DNA replication, cell-cycle progression, ribosome biogenesis, homologous recombination, and DNA strand elongation. Perturbations with predictable μ-orthogonal responses, by contrast, showed only weak and heterogeneous enrichment, scattering across unrelated regulatory, signaling, and metabolic processes. This is expected for a group defined by the absence of growth involvement rather than by any shared program. The dominant soft structure of the transcriptome therefore comprises a broad growth-related axis alongside lower-amplitude, pathway-specific directions that encode more targeted regulatory responses.

To test whether individual perturbations move cells reproducibly along this axis, we assigned each perturbation a factor 1+c, such that Δ*X*_µ_ = (1 + *c*)µ, where the sign of c represents the direction of total expression change. We identified genes appearing in at least two independent datasets whose knockdown or knockout produced a μ-parallel coefficient c of consistent sign and magnitude |c| > 0.2 in multiple datasets (**Fig. 6F**). These reproducible perturbations separated into two mechanistically coherent classes. The positive-c genes were dominated by DNA replication and cell-division machinery, including three MCM helicase subunits (*MCM2*, *MCM3*, *MCM6*) together with *TFDP1* and *HAUS4*. Disrupting S-phase progression or mitosis can leave cells accumulating RNA and biomass while failing to divide, so positive c appears to reflect globally amplified transcriptomes in enlarged or arrested cells rather than improved fitness. The negative-c genes, by contrast, included four Mediator subunits (*MED1*, *MED4*, *MED14*, *MED19*) alongside factors in rRNA processing (*POP7*), ribosome biogenesis (*UTP15*), and transcriptional elongation (*NELFB*)—a broad loss of biosynthetic capacity that contracts total RNA expression along the control mean. The fact that members of the same complexes recur on each side, *MCM* on the positive and Mediator on the negative, indicates a complex-level effect rather than gene-specific behavior. Both extremes are consistent with impaired proliferative fitness reached by different routes: positive-c perturbations uncouple biomass accumulation from division, while negative-c perturbations reduce the transcriptional and energetic capacity required for growth. We confirmed that CIPHER achieved high Pearson correlation for perturbations of these selected genes, whereas all other methods performed relatively poorly on the same set (**Fig. S13C**). None of this would be visible in normalized data, where these cell-size and growth phenotypes—carried entirely in the raw counts—are removed by construction. We next investigated the μ-orthogonal top 150 performers, contained 31 transcription factors and chromatin regulators, including ZFNs and ZBTBs, and had little effect on the total RNA expression (**Fig. S13B**). Lastly, CIPHER’s performance was largely independent of perturbation effect magnitude (ΔX) (R^2^ = 0.075), and did not depend on whether the perturbed gene was a transcription factor (**Fig. S14A-C**).

### Estimates of total genes affected during perturbations

We next sought to calculate the effective number of differentially expressed genes upon perturbation. CIPHER provides the total normalized contribution to the expression change of *gene_i_* due to correlations with *gene_j_*, and the analogous contribution of each gene to the total response. We calculated the entropy of these two probability vectors and exponentiated it to obtain an effective number of genes responsible for two kinds of estimates: 1) the expression change in the true perturbed gene, and 2) the total transcriptomic change upon perturbation (see **Methods 7**). Entropy-based estimates suggest that the transcriptomic response is distributed across a large number of genes (**Fig. 6H; Fig. S15**), consistent with prior findings^16^ that essential gene perturbations trigger widespread but diffuse expression changes. CIPHER predicts this quantity from the covariance alone, since the scaling parameter cancels in the fractional response. The effective number of responding genes predicted by CIPHER closely matched the observed value across datasets, with agreement improving steadily as forward-prediction accuracy increased (**Fig. S15A-E**). Many perturbed genes showed high self-variance contributions to their expression changes, implying that the perturbations directly affected the gene being targeted, as expected (**Fig. S12D**). Furthermore, while specific gene interactions could significantly affect perturbation responses, most essential gene perturbation effects appeared to emerge from many smaller correlations between the perturbed gene and its partners (**Fig. 6I**). Collectively, our results highlight that despite their presence, each soft mode represents a complex, yet quantifiable milieu of gene interactions.

### Normalization degrades the covariance structure underlying perturbation response

Throughout the manuscript, we use raw absolute expression counts X, motivated by our theoretical expectation of a dominant direction in raw gene expression space. Changing variables appropriately (see **Methods 8**), the log(1+X) transformation produces qualitatively similar results upon fitting single genes (**Fig. S16**). Frequency and library-size normalization, on the other hand, come with a number of unwanted consequences. First, these common normalization techniques exclude all variability due to total mRNA count, which can be considerable in perturbed cells (explored in detail in **Fig. 6C-F; Fig. S13A-C; S16**). Further, normalization also introduces spurious false correlations and can lead to a misalignment between the covariance matrix and perturbation response (**Methods 8**). Consistent with our finding that normalization discards biologically meaningful variation in total mRNA content, a recent single-cell gene expression reconstruction benchmark^67^ reported that supplying observed library size as an explicit input modestly improved gene expression reconstruction, and interpreted sequencing depth as carrying biological signal rather than being purely technical noise to be normalized away.

#### Linear response is invariant to measurement noise in raw counts

We asked how a linear response, defined in the underlying biology, could survive the noise introduced by measurement. We modeled sequencing as a two-layer process: a latent biological layer, where linear response holds, and an outer measurement layer that samples from it. When measurements are drawn from a Poisson or negative-binomial distribution and expression is kept in raw counts, we show that linear response passes through the measurement layer intact—the mean response and off-diagonal covariance transfer directly from the latent layer to the observed one (**Methods 4**). This transfer does not survive the nonlinear transformations routine in single-cell analysis (see **Methods 4**), unless appropriate change-of-variable operations are performed. Working in raw counts is therefore not merely a preprocessing choice but a condition for the framework to hold in this broad class of statistical models.

#### Normalization transformations can lead to a misalignment between perturbation and response

Our theoretical framework suggests that several commonly-used preprocessing schemes such as data normalization can lead to a misalignment between covariances and response (SI). This can happen because in the transformed units there may no longer be a dominant direction in transcriptional space. To test this idea, we turned to the eigenspectrum of the covariance matrix. Across all 27 datasets and common preprocessing schemes^68–71^ we tested—library-size normalization, log1p, PFlog, and frequency normalization (**Table 3**)—normalization suppressed the spectrum by orders of magnitude relative to raw counts (**Fig. S13D**). Frequency normalization was the most severe, compressing eigenvalues by six to eight orders of magnitude, while log1p, PFlog, and log1CP10k converged onto a common, flattened spectrum (**Fig. S13D**). The low rank modes that dominate the raw covariance are therefore distorted by transformation, and with them the directions along which perturbation response is transduced.

**Table 3:**
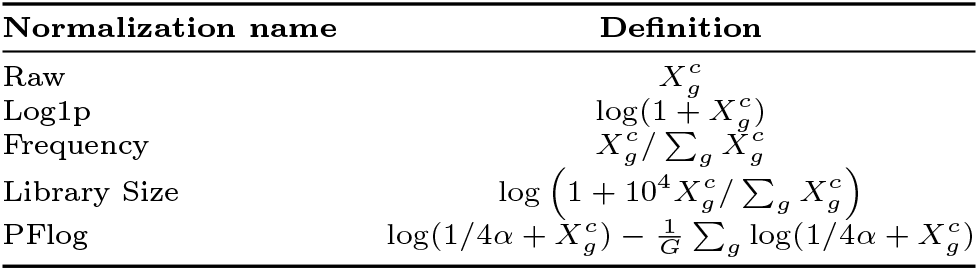
Normalization methods and their definitions.

**Table 4:**
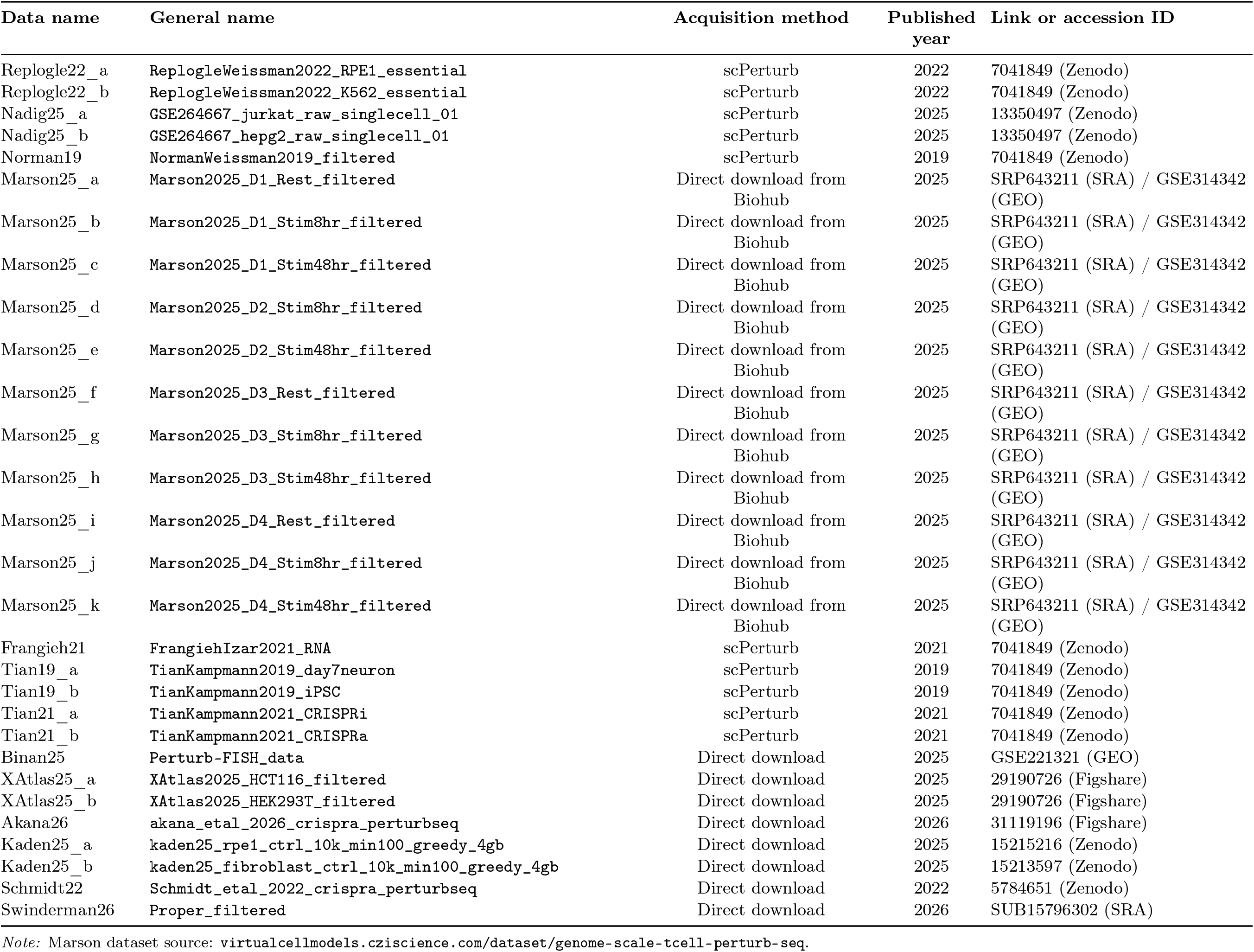
Data sources and acquisition information for the single-cell perturbation datasets used in this study.

We note however that a recently proposed variance-stabilizing PFlog^71^ normalization approach does preserve high CIPHER scores on several datasets (**Fig. S13E, S16).**

#### Synthetic data generated with scDesign3 model confirms raw counts preserve predictive signal

We tested how CIPHER’s performance depended on the exact representation of the original count matrix by generating synthetic single-cell expression data upon perturbations using the scDesign3 probabilistic model^72^ that preserves gene-wise marginal distributions and correlation structure (**Fig. 6J**). We created synthetic datasets with scDesign3 using two schemes: raw counts and library-size-normalized values. Fidelity of reconstructed embeddings from scDesign3 depended on the normalization schemes as measured by mLISI (Mean Local Inverse Simpson’s Index; mLISI of 1 and 2 indicates poor and perfect reconstruction, respectively). Raw counts performed the best for reconstructed embeddings, which was subsequently reflected in CIPHER’s performance on the Pearson correlation (**Fig. 6K,L**). Together, these analyses show that control populations’ covariance structure contains predictive information about perturbation responses.

#### Covariance from raw counts recovers validated interactions

We asked whether covariance-based predictions reflect real biological interaction rather than statistical coincidence (**Fig. S19**). Across all datasets, we compared predictions for gene pairs annotated as interacting in the BioGRID database^73^ against random pairs, testing both physical protein-protein and reported gene-gene interactions. Pearson correlation was consistently higher for annotated pairs (**Fig. S18A,B**), indicating that covariance-based predictions have potential to capture real biological signals. While covariance from raw and log-transformed counts captured validated interactions, the effect collapsed after library-size normalization, reflecting artifacts such as the spurious negative correlations, and compressed eigenspectrum that normalization introduces (**Fig. S18A,B**). This is consistent with the two-layer measurement model discussed above.

### CIPHER predictions validate experimentally on biological traits

We asked whether our framework could be applied to biological traits beyond single-cell perturbation datasets. Reasoning that targeted therapy acts as a perturbation driving cells from a therapy naive baseline to a resistant final state, we asked whether the fluctuations of the naive state could reveal which genes drive the transition. We first applied CIPHER to therapy resistance in melanoma, where we^74–76^ and others^77–82^ have demonstrated that coordinated, non-genetic fluctuations in the untreated cells can drive resistance outcome. We reanalyzed single-cell RNA sequencing data of clonal WM989-A6-G3-BRAF-V600E melanoma cells^74^ profiled for the therapy-naive state and after acquiring resistance to the BRAF/MEK inhibitors (**Fig. 7A,B**). We treated the naive and resistant populations as control and perturbed conditions, respectively. We calculated the covariance from control cells and measured the change in expression (ΔX) between control and resistant cells. Maximizing the posterior probability to rank genes by their ability to force the transition, we found *FN1* and *IGFBP7* as the top two mediators of resistance (**Fig. 7C**; **Table S1**, **Methods 9**). Some other top hits, such as *JUNB* and *JUN,* had been identified and validated previously^83^. CIPHER also recovered *ITGA3* and *AXL*, which a separate study validated as markers of states “primed” for resistance by prospective sorting and lineage tracing^84^. CIPHER’s top-ranked gene, *FN1,* was identified in that same study as a marker of essentially all primed cells but was never functionally tested. To test CIPHER’s predictions, we knocked down the two top genes *FN1* and *IGFBP7* individually by shRNA in a single-cell derived WM989 melanoma cell line (**Fig. 7D**). We found that both significantly reduced resistance to MEK inhibition (2.9 and 1.4 fold change for *FN1* and *IGFBP7* respectively; **Fig. 7E**), consistent with their high CIPHER posterior effect size ranking.

**Fig. 7:**
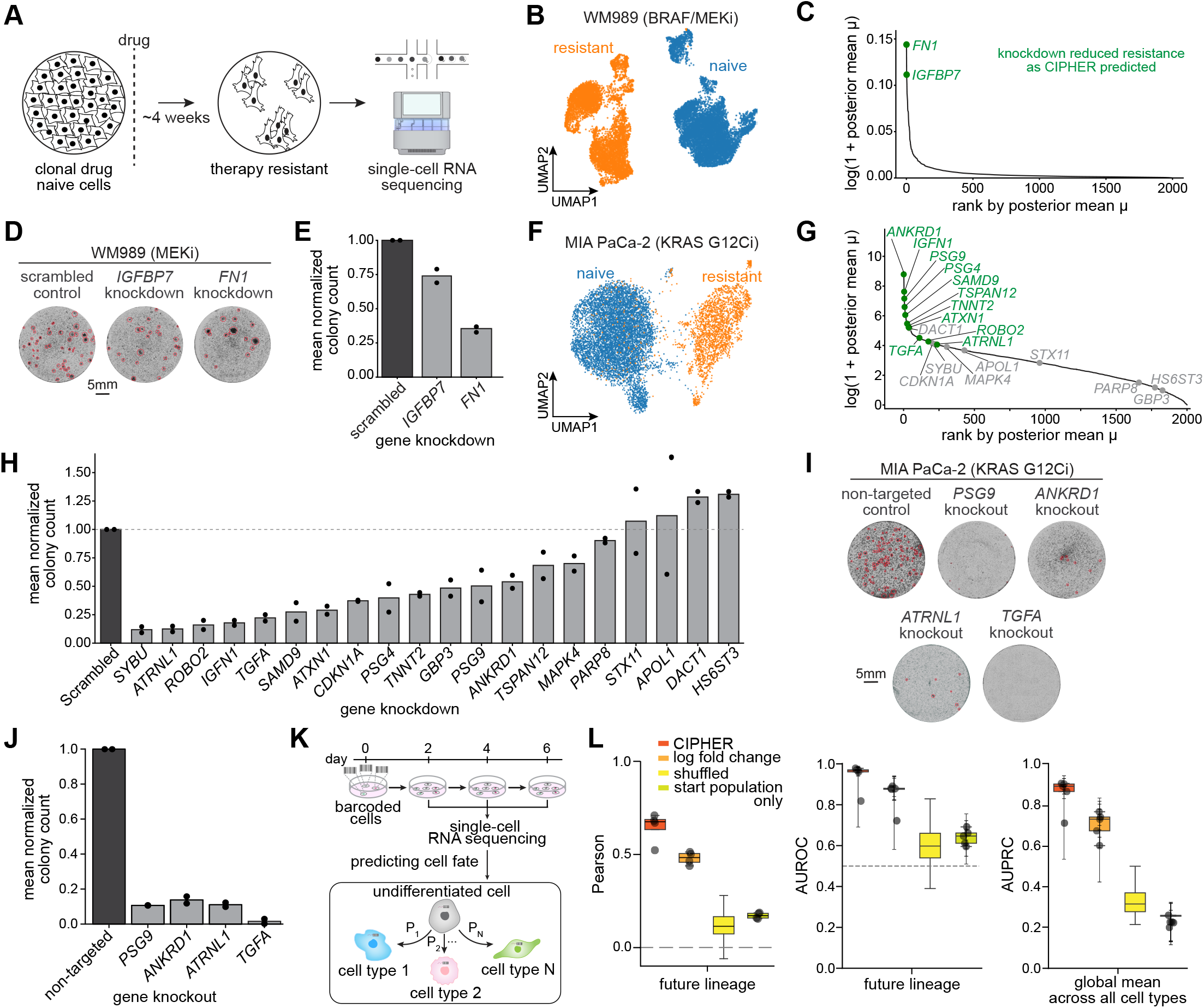
CIPHER predictions validate experimentally on therapy resistance and cell-fate traits. A) Schematic of the therapy resistance experiment. Clonal drug-naive cells were treated with targeted inhibitors to develop resistance; naive and resistant populations were profiled by single-cell RNA sequencing as control and perturbed conditions. B) UMAP of WM989 cells colored by naive (blue) and BRAF/MEK-inhibitor resistant (orange) populations. C) CIPHER ranking of candidate resistance drivers by posterior mean effect size μ. *FN1* and *IGFBP7*, the two top-ranked genes, are indicated. Green indicates genes whose knockdown reduced colony count (D, E) as predicted by CIPHER. D) Colony formation for scrambled control, *IGFBP7*, and *FN1* knockdowns in WM989 treated with MEK inhibitor. Scale bar, 5 mm. E) Mean normalized colony count for D (n = 2). Scrambled normalized to 1. F) UMAP of naive (blue) and resistant (orange) MIA PaCa-2 KRAS-G12C cells after sotorasib treatment. G) CIPHER ranking of candidate resistance drivers for MIA PaCa-2 by posterior mean μ. Tested genes labeled. Green indicates genes whose knockdown reduced colony count as CIPHER predicted. H) Mean normalized colony count after individual knockdown in MIA PaCa-2, ordered by effect (n = 2); scrambled normalized to 1 (dashed line). I) Colony formation for non-targeted control and CRISPR-Cas9 knockout of *PSG9*, *ANKRD1*, *ATRNL1*, *TGFA* in MIA PaCa-2. Scale bar, 5 mm. J) Mean normalized colony count for I (n = 2). Non-targeted normalized to 1. K) Experimental design for lineage prediction in the LARRY hematopoietic differentiation dataset. Barcoded hematopoietic progenitor cells were profiled by single-cell RNA sequencing over multiple time points (days 2, 4, and 6). Early undifferentiated cells are used by CIPHER to predict the probabilities of commitment to each terminal cell fate based on their transcriptional fluctuations. L) Performance of lineage prediction methods on the LARRY dataset. Left: Pearson correlation between predicted and observed future lineage commitment probabilities. Middle: area under the receiver operating characteristic curve (AUROC) for identifying the correct future lineage, with the dashed line indicating random performance. Right: area under the precision-recall curve (AUPRC) for the same task. Boxplots compare CIPHER scores, log fold change, shuffled covariance, and a baseline using only the starting population probabilities. Boxes indicate the interquartile range with median, whiskers extend to 1.5X the interquartile range, and individual points denote individual testing folds.

To test whether this predictive power extends beyond the (otherwise well-studied) non-genetic melanoma model of resistance, we turned to KRAS-driven cancers, where recently FDA-approved KRAS inhibitors have shown clinical promise^85,86^ but resistance remains a persistent barrier^87–90^. We clonally expanded the KRAS-G12C-mutant line MIA PaCa-2, exposed the cells to the KRAS G12C selective inhibitor sotorasib, and profiled the therapy-naive and resistant populations by single-cell RNA sequencing (**Fig. 7F**). CIPHER ranked the genes most likely to drive the naive-to-resistant transition (**Fig. 7G**; see **Methods 9**). To test this ranking, we selected 20 genes from across its full range and knocked them down individually, quantifying resistant colony formation after sotorasib treatment (**Fig. 7H**). Of the 11 tested genes within CIPHER’s top 200, 10 significantly reduced resistant colony formation, whereas genes low in the ranking largely left resistance unaffected (**Fig. 7G,H**). We then used CRISPR-Cas9 to disrupt four highly-ranked genes (*ANKRD1, PSG9, ATRNL1*, and *TGFA*) at their endogenous loci (**Fig. S17F-I**), which nearly eliminated resistance, further confirming CIPHER’s predictions (**Fig. 7I,J**).

Notably, most of these validated drivers showed relatively modest changes in expression and would have been overlooked by log-fold change, the standard approach for nominating drivers from expression data. Among genes upregulated in the resistant state, fold change was sharply bimodal: 18 exceeded a log₂ fold change of 25, while the remaining 1,874 fell below 13, with none in between (**Fig. S17A and Table S2**). Only three validated drivers belonged to the highest-fold-change mode—*ANKRD1* (CIPHER rank 1, log₂ fold change rank 1), *IGFN1* (ranks 4, 3), and *SAMD9* (ranks 12, 4). Of the remaining seven validated hits, CIPHER ranked four amongst the highest but were relatively modestly ranked in by the fold-change: *PSG9* (5, 27), *PSG4* (7, 28), *TSPAN12* (25, 50), and *ATXN1* (38, 88). The remaining three validated hits—*TNNT2* (36, 34), *TGFA* (112, 184), and *ROBO2* (175, 182)—exhibited broad agreement between CIPHER and log-fold rank. Conversely, only 8 of the 18 high-fold-change rank genes reached CIPHER’s top 100, the remainder ranking as low as 815 (**Table S2**). *PARP8* illustrates the cost of relying on fold-change alone: ranked 119th of 2,000 by fold change yet 1,661st by CIPHER, its knockdown left resistance unchanged, as did *HS6ST3* (469; 1,774) (**Fig. 7H; Fig. S17A**). Notably, these validated drivers were not among the most highly expressed genes in the naive state, spanning a broad range of baseline expression ranks in both the melanoma and pancreatic cancer models (**Fig. S17J,K**). Together, these results demonstrate that CIPHER recovers true resistance drivers across two unrelated cancers, drugs, and resistance mechanisms.

To test CIPHER’s applicability in yet another biological context, we turned to hematopoietic differentiation using LARRY-barcoded datasets^91^ (**Fig. 7K**). LARRY (lineage and RNA recovery) labels cells with a heritable, expressed DNA barcode, so that clonally related cells share a barcode and their initial transcriptional state can be linked to the mature fates their clonal sisters later adopt. This design gives us a covariance matrix with a controlled, known origin: rather than co-fluctuations across an arbitrary cell population, the covariance is computed over undifferentiated cells and used alongside the expression differences and information from lineage committed descendants to infer to what extent early fluctuations in a progenitor population influenced fate decisions. We reasoned that if this covariance structure carries fate information, it should predict the eventual lineage outcome of a barcode from the transcriptional state of its early clonal members. Indeed, CIPHER predicted future lineage outcomes with high accuracy (Pearson and AUROC on held-out future lineages; **Fig. 7L, left and middle**).

Furthermore, the CIPHER-predicted effective differentiation perturbation profiles for each cell type were significantly enriched for canonical cell-type markers (**Fig. S17B,C)**. This signal vanished under two controls: shuffling the fate labels and predicting from the starting population alone (which discards clonal information and asks whether the initial state distribution is sufficient on its own). CIPHER also outperformed both controls on a global measure aggregated across all cell types (AUPRC; **Fig. 7L, right; Fig. S17D,E**), confirming that the predictive signal comes specifically from the clonal covariance structure.

## DISCUSSION

Here, we developed a new conceptual framework combining linear response theory, Bayesian statistics, and regulatory networks to model rich single-cell perturbation experiment outcomes. We tested our framework CIPHER on synthetic networks, demonstrating its capability to predict systemic responses to perturbations. Applied to experimental single-cell perturbation datasets (**Table 1**), CIPHER reliably fit transcriptome-wide responses to gene perturbations—an effect that disappeared when gene-gene fluctuations in unperturbed cells were eliminated. Moreover, gene-gene fluctuations from one dataset performed similarly well in other datasets of the same cell type, highlighting the presence of stereotypic genome-wide fluctuation structures within cell types. Our study also identified the true perturbation with high-fidelity given only the covariance matrix and change in gene expression upon perturbation in both genetic- and chemical-screen datasets, while also revealing that transcriptome-wide responses tend to align with a low-dimensional set of soft eigenmodes. CIPHER matched or exceeded recent deep learning and linear methods while fitting a single parameter, and we validated its top driver predictions experimentally in two cancer models. In sum, our results demonstrate that theoretical approaches drawn from statistical physics offer powerful tools for interpreting the responses to a large class of perturbations, despite the underlying biological complexity. We emphasize that CIPHER is an interpretive framework: it recapitulates and dissects measured responses rather than forecasting responses to perturbations it has not seen.

The success of our study in recapitulating cellular responses reflects the broader power of linear response theory, which has proven effective across a diverse array of complex systems, including physical sciences and protein allostery^24–27^. CIPHER’s ability to accurately fit complex, transcriptome-wide responses using a simple, interpretable linear model suggests that biological systems may operate in regimes where linear response theory remains surprisingly effective despite underlying nonlinear dynamics. The presence of coordinated fluctuations and their important contribution to responses to genetic and environmental perturbations has been reported for diverse microbial and mammalian behaviors across scales including transcriptional, protein, and metabolic cell states^66,76,92,93^. In fact, the authors of one particular experimental study found steady-state single-cell gene correlations could predict the effect of p53 perturbation more accurately than even chromatin immunoprecipitation studies^66^. That same study found that certain correlated gene modules were shared between the U2OS and HEK293T lines while others were line-specific: the p53 module, for instance, was present in one line and absent in the other. This mirrors our own findings: covariance transferred between datasets of the same cell type but not across cell types, while covariances from unrelated cell types still outperformed their mean-field controls—a weaker but universal component. Yet, these observations had not been explicitly rationalized or quantified within a theoretical framework. As such, our study provides a strong theoretical footing toward a fundamental biological principle: complexity can emerge from or be effectively captured by simpler mechanistic rules encoded in the fluctuations of *unperturbed* cell states. Our theoretical model suggests that the specific form of linear response that CIPHER exploits could emerge when the underlying gene regulatory network is approximately low-rank. That CIPHER works across a diverse range of cell types and datasets suggests that such low-rank structures are ubiquitous. How such low-rank structure arises and is maintained in regulatory networks remains an open question for future work.

Linear models can be made uncertainty-aware, for instance through empirical Bayes moderation of coefficient variances as in the method limma,^94^ or by evaluating the Hessian of the likelihood to obtain standard errors on fitted coefficients. Such approaches quantify uncertainty in per-gene effect sizes, and the required curvature computations become costly as dimensionality grows. CIPHER’s Bayesian formulation instead places a posterior over the perturbation vector itself, allowing us to assign each gene a posterior inclusion probability that it has nonzero perturbation effect—a quantity that supports the inverse problem of identifying causal drivers rather than differential expression testing.

In its current linear-response formulation, CIPHER captures the first-order response to single perturbations and is not expected to model epistatic interactions between multi-gene perturbations, which are inherently nonlinear. As larger multi-gene perturbation datasets emerge, extending the framework to learn the nonlinear manifold of combinatorial responses is a promising direction. A related consideration is the interpretation of the scalar u*, best understood not as a perturbation strength but as the proportionality constant scaling covariance onto response. Because current high-throughput screens neither titrate nor measure knockdown strength explicitly, u* cannot yet be assigned from such experiments alone. Controlled technologies such as degron-based systems,^95,96^ which can titrate perturbation magnitude directly, could map how u* scales with perturbation strength, potentially allowing it to be set a priori in future work.

More broadly, given our results within and across cell-types, in principle, our framework could be harnessed to reveal the hierarchical relationships between cell types on underlying genome-wide fluctuation structures. Going further, we imagine linear response theory could be employed across species to probe the evolutionary history of gene-gene correlation. In its present form, CIPHER takes each dataset as input separately. Future studies could couple our findings with existing analytical and deep neural network approaches^97–103^ for joint embeddings of disparate datasets. Where prior knowledge of regulatory identity is available, the framework could also incorporate gene-class weighting or attention-like schemes to emphasize known regulators such as transcription factors, at some cost to its current parameter-free interpretability. Furthermore, incorporating a consistency loss that leverages pre-computed gene-gene covariance matrices could enhance the generative modeling of perturbation effects. By encouraging the model to preserve these covariance structures, this approach may improve generalization to unseen cell types and perturbations.

Our work demonstrated that transcriptome-wide perturbation effects can emerge through different modes: concentrated effects from a small subset of highly responsive genes (u_i_) versus distributed effects where many genes contribute modestly but cumulatively significant changes. These findings, combined with effect size calculations and comparison from TRADE^16^, point to the potential application of our findings in designing efficient attention or, even, tokenization mechanisms^104,105^ to enhance the predictive power of existing deep learning models or design newer, more expressive, cellular foundation models. While we effectively capture responses in many cases, it sometimes fails due to nonlinearities and factors beyond direct gene-gene fluctuations, as well as higher response complexity in some datasets. Incorporating the recently proposed single-cell clone tracing technologies^2,74,106,107^ with pooled CRISPR screening could enhance the accuracy by accounting for clone-specific fluctuations, as we demonstrate with hematopoietic differentiation example.

In a similar spirit to our work, spin-based, Ising-type models from statistical physics have recently been adapted to infer effective gene regulatory networks from single-cell RNA sequencing data and to predict cellular responses to perturbations ^108^. Looking ahead, we anticipate the emergence of technologies capable of capturing time-resolved single-cell perturbation dynamics, which would demand new theoretical frameworks. Response theories for (non)linear time-dependent perturbations^109^, fluctuation theorems^110,111^, and universal thermodynamic bounds and control theories^112–115^ are all derived from the recently developed nonequilibrium statistical physics of entire trajectories, and could be adapted to open new frontiers in the quantitative understanding of cellular regulation through the lens of nonequilibrium fluctuations.

Although beyond the scope of the current work, our work could be harnessed to predict unknown perturbations driving transitions in complex disease traits such as cancer metastasis and drug resistance to targeted therapy^75^, genetic disorders with unknown causal genes^116^, and more, significantly shortening the list of possible causal drivers. In such cases for example, one would compute the covariance of the naive cancer cells before drug as well as the response to drug and apply the inverse problem as we have formulated it here. This could constitute a standalone procedure without necessarily the need for integration of prior perturbation data, as demonstrated by other recent approaches combining trait and perturbation datasets^116^. Scaling up perturbation experiments with AI and statistical physics could enable efficient virtual screens to map genotype to phenotype beyond current single-trait CRISPR studies.

## ACKNOWLEDGEMENTS

We thank members of the Goyal lab for helpful discussions and comments on the manuscript. We thank the Feinberg Information Technology, Northwestern University Information Technology, and Quest High Performance Computing Cluster at Northwestern University Feinberg School of Medicine for their services. We acknowledge support from the National Institute for Theory and Mathematics in Biology (NITMB) through the National Science Foundation (DMS-2235451) and the Simons Foundation (MPTMPS-00005320). YG and SV thank the National Science Foundation through the Center for Living Systems Ignite Award (grant no. 2317138) as well. This research was supported in part by grant NSF PHY-2309135 to the Kavli Institute for Theoretical Physics (KITP) and the Gordon and Betty Moore Foundation Grant No. 2019.02. BKS acknowledges support from NUCATS T32, NITMB, and startup funds to YG from Northwestern University. LS, HS, PP and JJ were supported by funding from YG. MEM acknowledges support from Ryan Fellowship and the International Institute for Nanotechnology, NITMB, and Carcinogenesis NIH T32. NK acknowledges support from CSB postdoc fellowship and the HFSP long-term fellowship. EIG acknowledges support from the National Institute of Allergy and Infectious Disease of the National Institutes of Health (F31AI189214). B.H. was funded from IVADO and the Canada First Research Excellence Fund. We thank the Gene Editing, Transduction and Nanotechnology (GET iN) Core for providing shRNA constructs. Research reported in this publication was supported by the Northwestern University Skin Biology & Diseases Resource-Based Center of the National Institutes of Health under award number P30AR075049. A.G. acknowledges support from the DAE under project no. RTI4001, an Ashok and Gita Vaish Junior Researcher Award, as well as the Centre for Artificial Learning and Intelligence for Biological Research and Education, ICTS-TIFR. S.V. was supported by the National Institute of General Medical Sciences of the NIH under Award No. R35GM147400. YG acknowledges support from the Pew-Stewart Scholars Program and Burroughs Wellcome Fund Career Awards at the Scientific Interface. YG is a CZ Biohub Investigator.

## AUTHOR CONTRIBUTIONS

YG and BKS conceived and designed the study. BKS designed and performed a majority of the computational experiments and analyses with input from YG. LS and MEM collected and harmonized the datasets. HS, BKS, and YG prepared the figures and tables. NK, HS, PP performed the experiments. IS, JJ, HMLR, VRA, CW, QW, EIG, BH, JJL, AG and EP contributed to the analyses. SV and AG assisted BKS and YG in formulating the theoretical framework. BKS and YG wrote the manuscript with input from all authors.

## DATA AND CODE AVAILABILITY

This paper analyzes existing, publicly available genetic perturbation datasets 10/29 of which are available on the scPerturb database, and all the other datasets directly downloaded from the source. See **Table 4** for the information on individual datasets. All 29 datasets can be accessed through google drive (see github README for link). Two drug perturbation datasets are also publicly available: sci-Plex (https://cellxgene.cziscience.com/collections/00109df5-7810-4542-8db5-2288c46e0424) and Tahoe-100M (https://huggingface.co/datasets/tahoebio/Tahoe-100M). The following ATLAS datasets from CellxGene were used in this study: human embryonic meninges (https://datasets.cellxgene.cziscience.com/e2468d96-2b0a-4cfb-9ebe-4fecb8ca2b2c.h5ad), human intestine organoid ATLAS (https://datasets.cellxgene.cziscience.com/a8d79867-3e4d-4026-b2b1-5d0ddd6cb58e.h5ad), and human lung organoid ATLAS (https://datasets.cellxgene.cziscience.com/a489d69e-dd99-4663-9cfd-3d7e0e876103.h5ad). SIGNOR 4.0 Pathway Browser (https://signor.uniroma2.it/) provided the ground truth genetic interaction networks except for hematopoiesis. Hematopoiesis gene-gene interaction network was built by first listing the candidates from Moignard et al. (2013)^47^ and then formulating a transcription factor-gene interaction network based on TRRUST (https://www.grnpedia.org/trrust/). EMT network was obtained from Tripathi et al. (2020)^45^: https://github.com/st35/frustration-biological-networks. Protein-protein interaction network and genetic interaction map were built based on the BioGRID dataset (https://thebiogrid.org/). All code for the analyses in this manuscript has been deposited at: https://github.com/GoyalLab/CIPHER. The full CIPHER code package is downloadable through PyPI as “cipher-perturb”. We present a website to summarize our work: https://goyallab.github.io/CIPHERWebsite/.

## TABLES

Table S1: Melanoma (WM989) resistance-driver ranking: CIPHER posterior score versus log₂ fold change.

Table S2: MIA PaCa-2 resistance-driver ranking: CIPHER posterior score versus log₂ fold change.

**SUPPLEMENTARY CAPTIONS INCLUDED IN SI DOCUMENT**

