## Supplementary Material for "Co-fluctuations of genes in single cells predict transcriptome-wide outcomes to perturbations"

### Supplementary information: Co-fluctuations of genes in unperturbed single cells predict transcriptome-wide outcomes to genetic perturbations

Benjamin Kuznets-Speck[1,2,3,4*], Nitu Kumari[1,2,3,4], Irish Senthilkumar[1,2,3,4], Jaekwon Jung[1,2,3,4], Hector Manuel Lopez Rios[4,5], Leon Schwartz[1,2,3,4,6], Hanxiao Sun[1,2,3,4], Vidyesh Rao Anisetti[4,5], Changhu Wang[7], Qingyang Wang[7], Pornchanan Pholraksa[1,2,3,4], Emanuelle Grody[1,2,3,4], Madeline E. Melzer[1,2,3,4], Benjamin Haley[8], Ekta Prashnani[9], Jingyi Jessica Li[7], Akshit Goyal[10], Suriyanarayanan Vaikuntanathan[4,5], Yogesh Goyal[1,2,3,4,11*†]

[1]Department of Cell and Developmental Biology, Feinberg School of Medicine, Northwestern University, Chicago, IL, USA.
[2]Center for Synthetic Biology, Northwestern University, Evanston, IL, USA.
[3]Robert H. Lurie Comprehensive Cancer Center, Feinberg School of Medicine, Northwestern University, Chicago, IL, USA.
[4]NSF-Simons National Institute for Theory and Mathematics in Biology, Chicago, IL, USA.
[5]Department of Chemistry, University of Chicago, Chicago, IL, USA.
[6]Data Science of Systems Biology, Technische Universität München, Munich, Germany.
[7]Biostatistics Program, Fred Hutchinson Cancer Center, Seattle, WA 98109, United States.
[8]Department of Ophthalmology, Université de Montréal, Quebec, Canada.
[9]NVIDIA, Santa Clara, CA, USA.
[10]International Centre for Theoretical Sciences, Tata Institute of Fundamental Research, Bengaluru, India.
[11]Chan-Zuckerberg Biohub Chicago, Chicago, IL, USA.

†Lead contact

**Table 1**: Specifications for single-perturbation subset of the single-cell perturbation datasets used in this study.

| Alias ID | Sample type | CRISPR type | Perturb. type | Perturb. count | Control count | Gene count | Total cells | Source |
|---|---|---|---|---|---|---|---|---|
| Replogle22_a | RPE1 cells | CRISPRi | single | 654 | 10000 | 2535 | 137927 | [1] |
| Replogle22_b | K562 cells | CRISPRi | single | 1000 | 10000 | 2797 | 221295 | [1] |
| Nadig25_a | Jurkat cells | CRISPRi | single | 788 | 10000 | 2597 | 152211 | [2] |
| Nadig25_b | HepG2 cells | CRISPRi | single | 225 | 4976 | 2999 | 44091 | [2] |
| Norman19 | lymphoblasts | CRISPRa | single | 102 | 10000 | 1834 | 66391 | [3] |
| Marson25_a | $CD4^+$ T-cells | CRISPRi | single | 966 | 10000 | 3166 | 323529 | [4] |
| Marson25_b | $CD4^+$ T-cells | CRISPRi | single | 970 | 10000 | 4645 | 284722 | [4] |
| Marson25_c | $CD4^+$ T-cells | CRISPRi | single | 968 | 10000 | 4688 | 261396 | [4] |
| Marson25_d | $CD4^+$ T-cells | CRISPRi | single | 977 | 10000 | 4449 | 457477 | [4] |
| Marson25_e | $CD4^+$ T-cells | CRISPRi | single | 978 | 10000 | 3576 | 462770 | [4] |
| Marson25_f | $CD4^+$ T-cells | CRISPRi | single | 961 | 10000 | 3490 | 407131 | [4] |
| Marson25_g | $CD4^+$ T-cells | CRISPRi | single | 959 | 10000 | 4828 | 339227 | [4] |
| Marson25_h | $CD4^+$ T-cells | CRISPRi | single | 964 | 10000 | 4311 | 388729 | [4] |
| Marson25_i | $CD4^+$ T-cells | CRISPRi | single | 972 | 10000 | 4054 | 234451 | [4] |
| Marson25_j | $CD4^+$ T-cells | CRISPRi | single | 972 | 10000 | 3493 | 407674 | [4] |
| Marson25_k | $CD4^+$ T-cells | CRISPRi | single | 969 | 10000 | 4605 | 284487 | [4] |
| Frangieh21 | melanocytes | CRISPRi | single | 219 | 10000 | 2143 | 160661 | [5] |
| Tian19_a | iPSC-induced neuron | CRISPRi | single | 26 | 10000 | 92 | 79704 | [6] |
| Tian19_b | iPSC cells | CRISPRi | single | 26 | 10000 | 39 | 139642 | [6] |
| Tian21_a | iPSC-induced neurons | CRISPRi | single | 172 | 437 | 2912 | 30910 | [7] |
| Tian21_b | iPSC-induced neurons | CRISPRa | single | 91 | 434 | 1979 | 20230 | [7] |
| Binan25 | astrocytes | CRISPRi | single | 54 | 9633 | 211 | 4926 | [8] |
| XAtlas25_a | HCT116 cells | CRISPRi | single | 996 | 10000 | 4168 | 375749 | [9] |
| XAtlas25_b | HEK293 T-cells | CRISPRi | single | 994 | 10000 | 5613 | 501023 | [9] |
| Akana26 | Melanoma cells | CRISPRa | single | 57 | 2660 | 4273 | 19722 | [10] |
| Kaden25_a | RPE1 cells | CRISPRa | single | 1000 | 10000 | 2231 | 110000 | [11] |
| Kaden25_b | fibroblast cells | CRISPRa | single | 1000 | 10000 | 2878 | 110000 | [11] |
| Schemidt22 | T-cells | CRISPRa | single | 69 | 3274 | 1241 | 57817 | [12] |

**Table 2**: Specifications for the double-perturbation subset of the single-cell perturbation datasets used in this study.

| Alias ID | Sample type | CRISPR type | Perturb. type | Perturb. count | Control count | Gene count | Total cells | Source |
|---|---|---|---|---|---|---|---|---|
| Norman19 | lymphoblasts | CRISPRa | double | 126 | 11855 | 1000 | 52223 | [3] |
| Tian19_a | iPSC-induced neurons | CRISPRi | double | 322 | 15580 | 1000 | 42365 | [6] |
| Tian19_b | iPSC cells | CRISPRi | double | 322 | 1000 | 10687 | 57350 | [6] |
| Proper_filtered | K562 cells | CRISPRa | double | 104 | 2280 | 1000 | 29970 | [13] |

**Table 3**: Normalization methods and their definitions.

| Normalization name | Definition |
|---|---|
| Raw | $X_g^c$ |
| Log1p | $\log(1 + X_g^c)$ |
| Frequency | $X_g^c / \sum_g X_g^c$ |
| Library Size | $\log\left(1 + 10^4 X_g^c / \sum_g X_g^c\right)$ |
| PFlog | $\log(1/4\alpha + X_g^c) - \frac{1}{G} \sum_g \log(1/4\alpha + X_g^c)$ |

**Table 4:** Data sources and acquisition information for the single-cell perturbation datasets used in this study.

| Data name | General name | Acquisition method | Published year | Link or accession ID |
|---|---|---|---|---|
| Replogle22_a | `ReplogleWeissman2022_RPE1_essential` | scPerturb | 2022 | 7041849 (Zenodo) |
| Replogle22_b | `ReplogleWeissman2022_K562_essential` | scPerturb | 2022 | 7041849 (Zenodo) |
| Nadig25_a | `GSE264667_jurkat_raw_singlecell_01` | scPerturb | 2025 | 13350497 (Zenodo) |
| Nadig25_b | `GSE264667_hepg2_raw_singlecell_01` | scPerturb | 2025 | 13350497 (Zenodo) |
| Norman19 | `NormanWeissman2019_filtered` | scPerturb | 2019 | 7041849 (Zenodo) |
| Marson_a | `Marson2025_D1_Rest_filtered` | Direct download from Biohub | 2025 | SRP643211 (SRA) / GSE314342 (GEO) |
| Marson_b | `Marson2025_D1_Stim8hr_filtered` | Direct download from Biohub | 2025 | SRP643211 (SRA) / GSE314342 (GEO) |
| Marson_c | `Marson2025_D1_Stim48hr_filtered` | Direct download from Biohub | 2025 | SRP643211 (SRA) / GSE314342 (GEO) |
| Marson_d | `Marson2025_D2_Stim8hr_filtered` | Direct download from Biohub | 2025 | SRP643211 (SRA) / GSE314342 (GEO) |
| Marson_e | `Marson2025_D2_Stim48hr_filtered` | Direct download from Biohub | 2025 | SRP643211 (SRA) / GSE314342 (GEO) |
| Marson_f | `Marson2025_D3_Rest_filtered` | Direct download from Biohub | 2025 | SRP643211 (SRA) / GSE314342 (GEO) |
| Marson_g | `Marson2025_D3_Stim8hr_filtered` | Direct download from Biohub | 2025 | SRP643211 (SRA) / GSE314342 (GEO) |
| Marson_h | `Marson2025_D3_Stim48hr_filtered` | Direct download from Biohub | 2025 | SRP643211 (SRA) / GSE314342 (GEO) |
| Marson_i | `Marson2025_D4_Rest_filtered` | Direct download from Biohub | 2025 | SRP643211 (SRA) / GSE314342 (GEO) |
| Marson_j | `Marson2025_D4_Stim8hr_filtered` | Direct download from Biohub | 2025 | SRP643211 (SRA) / GSE314342 (GEO) |
| Marson_k | `Marson2025_D4_Stim48hr_filtered` | Direct download from Biohub | 2025 | SRP643211 (SRA) / GSE314342 (GEO) |
| Frangieh21 | `FrangiehIzar2021_RNA` | scPerturb | 2021 | 7041849 (Zenodo) |
| Tian19_a | `TianKampmann2019_day7neuron` | scPerturb | 2019 | 7041849 (Zenodo) |
| Tian19_b | `TianKampmann2019_iPSC` | scPerturb | 2019 | 7041849 (Zenodo) |
| Tian21_a | `TianKampmann2021_CRISPRi` | scPerturb | 2021 | 7041849 (Zenodo) |
| Tian21_b | `TianKampmann2021_CRISPRa` | scPerturb | 2021 | 7041849 (Zenodo) |
| Binan25 | `Perturb-FISH_data` | Direct download | 2025 | GSE221321 (GEO) |
| XAtlas25_a | `XAtlas2025_HCT116_filtered` | Direct download | 2025 | 29190726 (Figshare) |
| XAtlas25_b | `XAtlas2025_HEK293T_filtered` | Direct download | 2025 | 29190726 (Figshare) |
| Akana26 | `akana_etal_2026_crispra_perturbseq` | Direct download | 2026 | 31119196 (Figshare) |
| Kaden25_a | `kaden25_rpe1_ctrl_10k_min100_greedy_4gb` | Direct download | 2025 | 15215216 (Zenodo) |
| Kaden25_b | `kaden25_fibroblast_ctrl_10k_min100_greedy_4gb` | Direct download | 2025 | 15213597 (Zenodo) |
| Schmidt22 | `Schmidt_etal_2022_crispra_perturbseq` | Direct download | 2022 | 5784651 (Zenodo) |
| Swinderman26 | `Proper_filtered` | Direct download | 2026 | SUB15796302 (SRA) |

*Note:* Marson dataset source: virtualcellmodels.cziscience.com/dataset/genome-scale-tcell-perturb-seq.

### Supplementary Information for CIPHER

Here we detail the methods and procedures for CIPHER. We start by stating the fundamental result and how to use it in the forward and inverse problems. After this, we derive linear response theory in general settings and motivate its use in describing transcriptional changes. Next, we discuss simulations for the synthetic regulatory networks considered in the main text. From here, we consider the 'forward' and 'inverse' problems in more detail, and describe the analysis surrounding our discussion of collective soft modes in propagating the response of gene expression to cellular perturbation. We then make the case for using raw count data, discussing in detail how linear response is most naturally formulated at the absolute counts level, and explicitly derive how linear response changes and what is lost when gene expression is normalized. Finally, we detail our data preprocessing steps and compare to other perturbation-response methods.

### 1 Effective low-rank theory of perturbation response in gene regulatory networks

CIPHER predicts the transcriptome-wide response to a knockdown of gene $j$ from the unperturbed gene–gene covariance, $\Delta X_i \approx \Sigma_{ij}\, u_j^*$. Here we propose a simple theory, based on nonlinear dynamics and random matrix theory, for the origin of this predictability. We argue that CIPHER should work generically for gene regulatory networks (GRNs) which have an approximate rank-1 structure on top of a background disordered network. We show that when the rank-1 coupling is strong enough, both the perturbation response and the gene–gene covariance align along a common dominant direction, which makes the covariance proportional to the response. We demonstrate that this alignment is robust to disorder in the network and to intrinsic gene expression noise, and that CIPHER's predictions will tend to work best near a cell-fate transition. While our analytical calculations are performed in a simple and tractable limit, we confirm with simulations on synthetic and biological networks that a dominant mode in the GRN is sufficient for CIPHER to predict perturbation response. Finally, we discuss several predictions of the theory and how they match observations in the data.

#### 1.1 Setup: noisy nonlinear dynamics of gene regulatory networks (GRNs)

We consider a gene regulatory network (GRN) of $N$ genes whose dynamics of production and degradation are governed by the following chemical Langevin equations

$$\dot{x}_i = \Gamma_i h_i(x, \theta) - \gamma_i x_i + \sqrt{\Gamma_i h_i(x, \theta)}\, \zeta_i^+(t) - \sqrt{\gamma_i x_i}\, \zeta_i^-(t). \tag{S1}$$

where $\Gamma_i$ is the maximal production rate, $\gamma_i$ the degradation rate, and $h_i(x)$ the regulatory input to gene $i$, a product of Hill functions of its activators and repressors. Production and degradation are discrete random events and typically follow a Poisson process, resulting in noise in the dynamics. We model this using the last two terms, where $\zeta_i^+(t)$ and $\zeta_i^-(t)$ represent independent white-noise processes. The first noise term captures production noise while the second captures degradation noise.

At a fixed point production balances degradation, $\Gamma_i h_i(x^*) = \gamma_i x_i^*$, and genes have different mean expression levels $x_i^*$, with noise whose variance scales with the mean. Linearizing around the fixed point $x^*$, we see that the Jacobian is

$$J = -D + K, \tag{S2}$$

with $D = \mathrm{diag}(\gamma_i)$ the degradation rates and $K_{ij} = \Gamma_i\, \partial h_i / \partial x_j |_{x^*}$ the coupling, encoding how gene $j$ influences the production of gene $i$. For simplicity, we consider uniform degradation, $D = \gamma I$. The matrix $K_{ij}$ thus encodes the gene regulatory network.

#### 1.2 Approximate rank-1 structure of GRNs

Following several past works, our theory models the GRN $K$ as a random network. Specifically, we model $K$ as containing a rank-1 mode on top of a random disordered network given by

$$K = c\, \hat{u}\hat{w}^\top + W. \tag{S3}$$

The first term represents a rank 1 regulatory mode, with the number $c > 0$ representing a coupling strength, the vector $\hat{u}$ is the response profile (the combination of genes that respond when the mode is activated) and the vector $\hat{w}$ is the driver profile (the combination of genes that drive it). Generally, we do not expect $\hat{u}$ and $\hat{w}$ to be the same combinations of genes; there will be some overlap $\kappa = \hat{w}^\top \hat{u}$ between them which lies between 0 and 1.

The second term $W$ represents the disordered interaction network across the GRN that operates in addition to the structured rank 1 mode. This is the network that contains cross-talk between pathways and context-dependent couplings. We assume for simplicity that $W$ has independent random entries given by

$$W_{ij} \sim \mathcal{N}(0, \sigma^2/N), \tag{S4}$$

with $W_{ij}$ and $W_{ji}$ being uncorrelated with each other as we would expect in a directed GRN. The scaling with $N$ is assumed merely for mathematical convenience to have a good thermodynamic limit as $N \to \infty$ and does not affect our central results. Note that the disorder $W$ might follow different statistics; here we consider Gaussian interactions only for concreteness.

Several biological observations motivate this choice of modeling $K$ as rank-1 plus disorder. For instance, evolving random GRNs to minimize frustration has been shown to result in such an approximate rank-1 structure [14]. However, minimal frustration is one of many possible sources of a roughly rank-1 interaction matrix. Recent work has shown that selecting networks for robust homeostatic recovery from perturbations also produces a low-rank coupling with a soft mode [15], and there might be other sources or mechanisms as well. Our theory is agnostic to these mechanisms and shows that a rank-1 dominant structure is sufficient to explain how CIPHER may work.

#### 1.3 BBP transition for a rank-1 perturbation to a random matrix

Consider the random matrix

$$A = c\, \hat{u}\hat{w}^\top + W, \tag{S5}$$

where $W$ is an $N \times N$ real Ginibre matrix with iid entries given by the following statistics

$$\langle W_{ij} \rangle = 0, \qquad \langle W_{ij}^2 \rangle = \frac{\sigma^2}{N}, \tag{S6}$$

and $\hat{u}$ and $\hat{w}$ are fixed $N$-dimensional vectors. The parameter $c$ controls the strength of the rank 1 perturbation. Although $W$ is real, its eigenvalues are generally complex. For large $N$, the eigenvalue distribution is given by the circular law, so that the spectrum of $W$ occupies a disk of radius $\sigma$ in the complex plane.

To determine whether the perturbation creates an isolated eigenvalue, i.e., a BBP-style outlier, we consider the characteristic equation

$$\det(zI - A) = 0. \tag{S7}$$

Substituting Eq. (S5) gives

$$\det(zI - W - c\hat{u}\hat{w}^\top) = 0. \tag{S8}$$

For $z \notin \mathrm{spec}(W)$, the matrix $zI - W$ is invertible. Applying the matrix determinant lemma,

$$\det(A + xy^\top) = \det(A)\left(1 + y^\top A^{-1} x\right), \tag{S9}$$

with

$$A = zI - W, \qquad x = -cu, \qquad y = \hat{w}, \tag{S10}$$

yields

$$\det(zI - A) = \det(zI - W)\left[1 - c\,\hat{w}^\top (zI - W)^{-1}\hat{u}\right]. \tag{S11}$$

Hence any eigenvalue of $A$ outside the spectrum of $W$ must satisfy

$$1 = c\,\hat{w}^\top (zI - W)^{-1}\hat{u}. \tag{S12}$$

For $|z| > \sigma$, we can use the expansion

$$(zI - W)^{-1} = \frac{1}{z}\sum_{k=0}^{\infty}\left(\frac{W}{z}\right)^k. \tag{S13}$$

Therefore,

$$\hat{w}^\top (zI - W)^{-1}\hat{u} = \frac{\hat{w}^\top \hat{u}}{z} + \sum_{k=1}^{\infty}\frac{\hat{w}^\top W^k \hat{u}}{z^{k+1}}. \tag{S14}$$

Since $W$ is isotropic and $\hat{u}$ and $\hat{w}$ are random directions, we expect the overlap $\hat{w}^\top W^k \hat{u} \to \mathcal{O}(1/\sqrt{N})$ for all $k \geq 1$. Consequently,

$$\hat{w}^\top (zI - W)^{-1}\hat{u} \longrightarrow \frac{\hat{w}^\top \hat{u}}{z}, \qquad |z| > \sigma. \tag{S15}$$

Substituting Eq. (S15) into Eq. (S12) gives

$$z_{\text{out}} = c\,\hat{w}^\top \hat{u}. \tag{S16}$$

Note that we have thus far assumed that the outlier lies outside the support of the circular law. Therefore, to be self-consistent, an isolated eigenvalue exists only if

$$|z_{\text{out}}| > \sigma. \tag{S17}$$

Combining Eqs. (S16) and (S17) gives us the criterion for the transition

$$|c\,\hat{w}^\top \hat{u}| > \sigma. \tag{S18}$$

##### 1.4 With a mode gap, perturbation response occurs along a single direction

Experiments like CRISPRi typically perturb the parameters of a GRN such as the production rates of different genes. In our model, such a perturbation would result in the fixed point changing, and it is this response in the fixed point that we wish to predict. We will first illustrate this for the case of deterministic GRN dynamics, neglecting noise; in the next section we discuss the effects of noise.

Differentiating the fixed-point condition $F(x^*(\theta);\theta) = 0$ (where $F$ is the deterministic part of Eq. S1) with respect to a parameter $\theta_\mu$ (e.g., the production rate of gene $\mu$), we get that

$$\sum_j J_{ij}\frac{\partial x_j^*}{\partial \theta_\mu} + \frac{\partial F_i}{\partial \theta_\mu} = 0 \quad\Longrightarrow\quad A = -J^{-1}B, \qquad A_{i\mu} = \frac{\partial x_i^*}{\partial \theta_\mu}. \tag{S19}$$

where $A$ is the susceptibility matrix capturing how a gene's steady-state expression level changes with changes in the production rate of gene $\mu$. Each gene knockdown perturbs a gene's production rate and thus the perturbation matrix $B$ is diagonal and we may absorb its scale into parameters and assume $B = I$ for simplicity. With uniform degradation $\gamma_i = \gamma$ we have that

$$A = (\gamma I - K)^{-1}. \tag{S20}$$

This susceptibility $A$ is the key object we need to compute, since it is the linear response matrix that captures how gene expressions respond to perturbations in another gene. With only disorder this is easy to compute, i.e., when $K = W$, we can use the cavity method (Appendix B) to show that

$$(\gamma I - W)^{-1} \approx \frac{1}{\gamma}I, \tag{S21}$$

because each row and column of $W$ is independent of the others and of the rest of the matrix, so all corrections average to zero. We can now add the rank-1 mode to this and apply the Sherman–Morrison identity to get

$$A = \frac{1}{\gamma} I + \alpha\, \hat{u}\hat{w}^{\top}, \qquad \alpha = \frac{c}{\gamma(\gamma - c\kappa)}. \tag{S22}$$

The response to any perturbation is a vector in the $N$-dimensional space of gene expression, and in general we might expect it to be complicated: each gene we perturb might push the cell in its own direction, so that capturing the full response would require all of $A$. The response is simple only in the opposite case, where one of these directions dominate the response. Cells then respond roughly along the same direction to a wide variety of perturbations. To see whether $A$ has such a dominant direction, we must look at its eigenvalues.

From Eq. (S22) the eigenvalues of $A$ are easy to read off. The featureless part $\frac{1}{\gamma}I$ gives $N$ equal eigenvalues $1/\gamma$ that prefer no direction, and the mode $\alpha\, \hat{u}\hat{w}^{\top}$ adds another eigenvalue. In Appendix A, we show that when

$$c\kappa > \sigma, \tag{S23}$$

the matrix undergoes a BBP transition and the eigenvalue due to the rank-1 mode "pops out" of the disordered bulk of the eigenvalues of $W$ and becomes an outlier eigenvalue (see 1.3 for full derivation). Below this threshold the mode is buried in the disorder and $A$ has no dominant direction. Above it, as the mode gap $c\kappa - \sigma$ increases, the outlier stands out distinctly from the bulk, and $A$ can be well-approximated by a single mode. We note that the fixed point is stable only when $\gamma > c\kappa$.

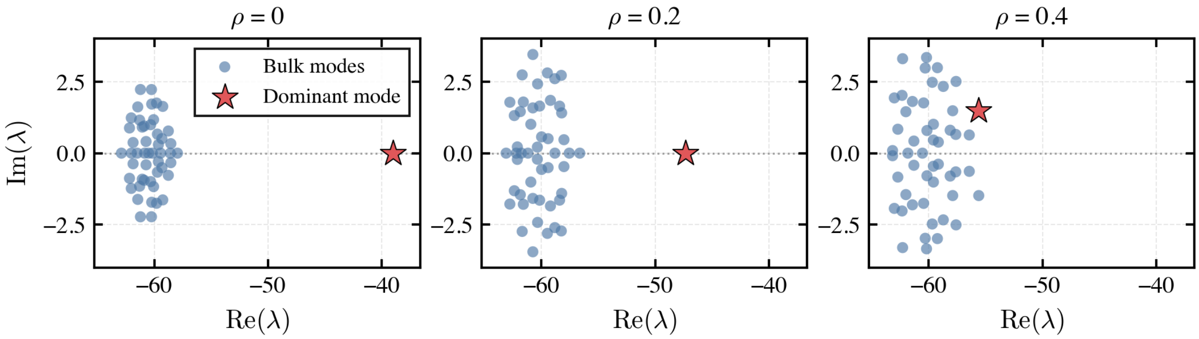


**Fig. S1**: Eigenvalue spectrum of the Jacobian for a random gene regulatory network (GRN) with $N = 50$ genes. The leftmost panel shows the spectrum of random GRN with zero frustration ($\rho = 0$ is our frustration metric, see 1.10 for details of methods and parametrization), highlighting a dominant outlier eigenvalue (red star) clearly separated from the bulk modes (blue circles). As we increase frustration $\rho$ by modifying the interactions in the random GRN, this dominant eigenvalue shifts and is progressively absorbed into the bulk.

This argument lets us predict perturbation response in a simple way. Knocking down gene $j$ lowers its production rate, $\epsilon = \epsilon_j \hat{e}_j$, and $\Delta x = -A\epsilon$. The dominant mode spreads the perturbation across the network: for $i \neq j$,

$$\Delta x_i = -\epsilon_j\, \alpha\, \hat{w}_j\, \hat{u}_i \propto \hat{u}_i. \tag{S24}$$

Whichever gene we knock down, the response points along the same direction $\hat{u}$, with a size set by how strongly that gene drives the mode through $\hat{w}_j$. This is the single dominant direction we sought. Thus, with a rank-1 mode structure and a sufficiently large mode gap, perturbation response in a complex nonlinear GRN becomes simple.

#### 1.5 Gene–gene covariance aligns with the perturbation response

CIPHER does not measure the response directly; it reads the gene–gene covariance across a population of unperturbed cells. To connect the two, note that cells in a population are not identical: they differ in their production and degradation rates because of variation in size, cell-cycle stage, and microenvironment. Writing the parameters of each cell as a population mean plus a small deviation, $\theta = \bar{\theta} + \delta\theta$, and recalling that $A$ maps a change in parameters to a shift in the fixed point, each cell sits at a slightly displaced steady state, $\delta x = A\,\delta\theta$. The gene–gene covariance across the population is then

$$\Sigma = \langle \delta x\, \delta x^\top \rangle = A\, C_\theta\, A^\top, \tag{S25}$$

with $C_\theta = \langle \delta\theta\, \delta\theta^\top \rangle$ the covariance of the parameters across cells. Assuming that this variation is isotropic for simplicity, i.e., $C_\theta = \sigma_\theta^2 I$, we get that $\Sigma = \sigma_\theta^2\, AA^\top$.

Thus, just like the perturbation response, the gene–gene covariance also depends strongly on the susceptibility $A$. If $A$ has a mode gap as discussed previously, $A \approx \frac{1}{\gamma} I + \alpha\, \hat{u}\hat{w}^\top$, and $AA^\top$ is dominated by $\hat{u}\hat{u}^\top$ as follows,

$$\Sigma = \sigma_\theta^2 \Big( \frac{1}{\gamma^2}\, I + \alpha^2\, \hat{u}\hat{u}^\top + \frac{\alpha}{\gamma}\, (\hat{u}\hat{w}^\top + \hat{w}\hat{u}^\top) \Big) \approx \sigma_\theta^2 \Big( \frac{1}{\gamma^2}\, I + \alpha^2\, \hat{u}\hat{u}^\top \Big), \tag{S26}$$

when $\alpha/\gamma$ is sufficiently small. Column $j$ of the covariance therefore points mostly along $\hat{u}$, which is the same direction as the response to knocking down gene $j$ we derived previously (Eq. (S65)). Thus, we see that the unperturbed gene–gene covariance column and the perturbation response are proportional to each other. This immediately leads us to exactly the central equation we use in CIPHER:

$$\Delta x_i \approx \Sigma_{ij}\, u_j^* \tag{S27}$$

with the fitted scalar $u_j^*$ playing the role of the proportionality constant.

#### 1.6 Robustness to adding noise in GRN dynamics

So far the covariance came entirely from cell-to-cell variation in parameters. But cells with identical parameters still fluctuate, because production and degradation are stochastic events. We now show that this intrinsic noise produces a covariance of the same form as perturbations to parameters we showed previously. Consequently, our central result that response is proportional to gene–gene covariance still holds.

Linearizing the noisy dynamics about the fixed point gives,

$$\dot{\delta x} = J\, \delta x + \xi(t), \qquad \langle \xi_i(t) \xi_j(t') \rangle = 2 D_{ij}\, \delta(t - t'), \qquad D = \gamma\, \mathrm{diag}(x^*), \tag{S28}$$

where the Poisson relation $\Gamma_i h_i(x^*) = \gamma_i x_i^*$ sets the diagonal matrix $D$. The steady-state covariance $\Sigma = \langle \delta x\, \delta x^\top \rangle$ obeys a balance equation, which we can obtain by differentiating $\langle \delta x_i \delta x_j \rangle$ in time and using the dynamics as follows:

$$\frac{d}{dt} \langle \delta x_i \delta x_j \rangle = (J\Sigma)_{ij} + (\Sigma J^\top)_{ij} + \langle \xi_i \delta x_j \rangle + \langle \delta x_i \xi_j \rangle. \tag{S29}$$

The first two terms relax fluctuations back to the fixed point through $J$; the last two are the noise that drives them. Writing $\delta x_j$ as the accumulated response to past noise, the delta function contributes at the end of the integration window with half its weight, giving $\langle \xi_i \delta x_j \rangle = D_{ij}$. At steady state the time derivative vanishes, and

$$J\Sigma + \Sigma J^\top = -2D. \tag{S30}$$

This is the Lyapunov equation, which shows that at a stationary state, the variance added by noise balances the attenuation of noise through dynamics. Note that in linearizing the dynamics and deriving this Lyapunov equation we have neglected additional terms in the appropriate expansion. This assumption might not be valid in the limit of low molecule numbers as in GRNs [16]. Nevertheless, we will continue with this approximation for simplicity. A more detailed calculation will involve incorporating more terms, which we leave for future work.

To understand the consequences of noise, we must self-consistently solve this Lyapunov equation. To do so, we need an ansatz for $\Sigma$ which we can solve self-consistently. Since $J$ involves only the directions $\hat{u}$ and $\hat{w}$, a reasonable ansatz is that the covariance $\Sigma$ lies in the span of $I$, $\hat{u}\hat{u}^\top$, $\hat{w}\hat{w}^\top$, and $\hat{u}\hat{w}^\top + \hat{w}\hat{u}^\top$:

$$\Sigma = a\,I + b\,\hat{u}\hat{u}^\top + f\,\hat{w}\hat{w}^\top + d\,(\hat{u}\hat{w}^\top + \hat{w}\hat{u}^\top), \tag{S31}$$

where $a, b, f$ and $d$ are constants to be determined. We now substitute this ansatz into Eq. (S30). We also use the rank-1 approximation to $J = -\gamma I + c\,\hat{u}\hat{w}^\top$, where in doing so we have assumed that we are past the BBP threshold we discussed previously such that there is a sufficient mode gap between the rank-1 mode and the disordered bulk of $W$. After making these substitutions, we see that every product goes through $\hat{w}^\top\hat{u} = \kappa$ and $\hat{w}^\top\hat{w} = 1$, and we can easily obtain the coefficients by matching the four matrices:

$$a = D_0/\gamma, \qquad f = 0, \qquad d = \frac{c\,D_0}{\gamma(2\gamma - c\kappa)}, \qquad b = \frac{c^2 D_0}{\gamma(\gamma - c\kappa)(2\gamma - c\kappa)}. \tag{S32}$$

This is the same structure as the parameter covariance we found above. It contains a diagonal, a dominant term $b\,\hat{u}\hat{u}^\top$ along $\hat{u}$, and a mixed term along $\hat{u}\hat{w}^\top + \hat{w}\hat{u}^\top$. The only difference between this case with noise and the previous case with parameter perturbations in Eq. (S26) is that we obtain different coefficients for the covariance.

Thus, using the same logic as the previous section, we can see that when there is a mode gap between the rank-1 mode and the rest of the spectrum of $J$, the perturbation response $\Delta x$ is proportional to gene–gene covariance $\Sigma$, just as it did for parameter perturbations. This response also points along the $\hat{u}$ direction. This suggests that including intrinsic gene expression noise should not qualitatively change the predictability of CIPHER.

Rather, just like the previous section, the cross-term with strength $d$ may slightly rotate the covariance column away from $\hat{u}$ toward $\hat{w}$. This merely affects the relative size of the contaminant compared to the signal, which we can easily compute as

$$\frac{d}{b} = \frac{\gamma - c\kappa}{c} = \frac{1}{\eta}, \qquad \eta = \frac{c}{\gamma - c\kappa}, \tag{S33}$$

where the recycling number $\eta$ is the ratio of rank-1 mode's coupling strength to the intrinsic relaxation rate of the dynamics. The more times a fluctuation passes through the mode, the more the output direction $\hat{u}$ dominates the input direction $\hat{w}$. We can show that we obtain the same relative size of the contaminant even in the absence of any intrinsic Poisson fluctuations in expression levels. Finally, the diagonal term $a = D_0/\gamma$ merely increases every gene's variance without rotating the column away from $\hat{u}$.

#### 1.7 Robustness to shifted CLR normalization

So far we have treated the expression levels $x_i$ as directly observed. In single-cell RNA-seq, however, these quantities are affected by cell-specific capture efficiency and sequencing depth. A commonly used preprocessing step is shifted centered log-ratio (shifted CLR) normalization, which reduces the strong dependence of count variance on expression level and attenuates sensitivity to cell-to-cell variation in sequencing depth [17]. We now ask how such a preprocessing step affects the covariance–response alignment that underlies CIPHER. The main point is that shifted CLR acts locally as a change of coordinates. Therefore, in the ideal rank-1 limit, both the perturbation response and the covariance are transformed by the same local linear map. This preserves CIPHER's logic in principle, but only if the transformed rank-1 mode remains dominant after normalization.

Let $s = \sum_i x_i$ denote the total count in a cell. Shifted CLR is

$$z_i = \log\left(1 + \tau\frac{x_i}{s}\right) - \frac{1}{N}\sum_k \log\left(1 + \tau\frac{x_k}{s}\right). \tag{S34}$$

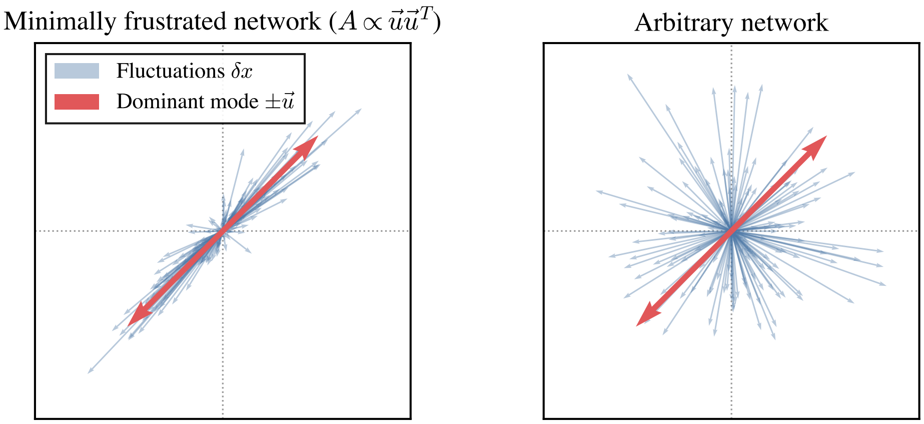


**Fig. S2**: Schematic representation of expression-level fluctuations ($\delta x$) driven by genetic perturbations (e.g., knock-downs) and intrinsic gene expression noise. **(Left)** In a minimally frustrated network, the interaction network and thus susceptibility is approximately rank-1 ($A \propto \hat{u}$). Consequently, the steady state fluctuations primarily relax along a dominant direction $\hat{u}$ (red arrows). **(Right)** In a highly frustrated network there is generally no dominant collective mode, and the susceptibility $A$ is generally high-rank. Thus, fluctuations relax along several directions in gene expression space, and in these cases, we do not expect CIPHER to work.

Equivalently, this can be written as

$$z = P\log(x + x_0\mathbf{1}), \qquad P = I - \frac{1}{N}\mathbf{1}\mathbf{1}^\top, \tag{S35}$$

where $x_0 = s/\tau$ is the effective count-scale pseudocount. The shifted logarithm reduces the influence of large absolute counts, while the matrix $P$ subtracts the mean log expression within each cell. Thus, shifted CLR approximately converts raw count changes into relative changes in expression while removing the common direction associated with sequencing depth or total measured RNA.

Although shifted CLR is nonlinear, we may linearize it about the fixed point $x^*$. Let

$$s^* = \sum_i x_i^*, \qquad x_0^* = \frac{s^*}{\tau}, \qquad q^* = \frac{x^*}{s^*}. \tag{S36}$$

For a small perturbation $\delta x$, the total count changes by

$$\delta s = \mathbf{1}^\top \delta x. \tag{S37}$$

Differentiating the relative abundance $x_i/s$ gives

$$\delta\left(\frac{x_i}{s}\right) = \frac{1}{s^*}\left(\delta x_i - q_i^*\delta s\right). \tag{S38}$$

Therefore,

$$\delta\log\left(1 + \tau\frac{x_i}{s}\right) = \frac{1}{x_i^* + x_0^*}\left(\delta x_i - q_i^*\delta s\right). \tag{S39}$$

In vector form, this gives

$$\delta z \simeq T\,\delta x, \qquad T = PL\left(I - q^*\mathbf{1}^\top\right), \qquad L = \mathrm{diag}\left(\frac{1}{x_i^* + x_0^*}\right). \tag{S40}$$

Eq. (S40) shows how shifted CLR acts locally on raw-count fluctuations. Reading from right to left, the factor $I - q^*\mathbf{1}^\top$ removes the direction that simply scales all expressions by a factor without affecting gene frequencies. Indeed,

$$\left(I - q^* \mathbf{1}^\top\right) x^* = 0. \tag{S41}$$

We can now apply the same logic as in the previous sections. In raw-count coordinates, the dominant-mode response to knockdown of gene $j$ is

$$\Delta x^{(j)} \simeq -\epsilon_j \alpha \hat{w}_j \hat{u}, \tag{S42}$$

where $\hat{u}$ is the raw-count direction along which the dominant regulatory mode moves the transcriptome around $x^*$. Applying Eq. (S40), the corresponding response in shifted CLR coordinates is

$$\Delta z^{(j)} \simeq -\epsilon_j \alpha \hat{w}_j T \hat{u}. \tag{S43}$$

Thus the response remains one-dimensional in the ideal rank-1 limit, but the observed response direction changes from $\hat{u}$ to

$$\hat{u}_{\mathrm{CLR}} = T\hat{u} = PL\left(I - q^* \mathbf{1}^\top\right) \hat{u}. \tag{S44}$$

Likewise, using the covariance structure derived in the previous section,

$$\Sigma = aI + b\,\hat{u}\hat{u}^\top + f\,\hat{w}\hat{w}^\top + d(\hat{u}\hat{w}^\top + \hat{w}\hat{u}^\top), \tag{S45}$$

the covariance in shifted CLR coordinates is

$$\Sigma^{(z)} = T\Sigma T^\top = a\,TT^\top + b\,(T\hat{u})(T\hat{u})^\top + f\,(T\hat{w})(T\hat{w})^\top + d\left[(T\hat{u})(T\hat{w})^\top + (T\hat{w})(T\hat{u})^\top\right]. \tag{S46}$$

The term proportional to $b$ is the transformed rank-1 signal. If this term dominates the others, then

$$\Sigma^{(z)} \simeq b\,(T\hat{u})(T\hat{u})^\top = b\,\hat{u}_{\mathrm{CLR}}\hat{u}_{\mathrm{CLR}}^\top, \tag{S47}$$

and the covariance column remains aligned with the shifted-CLR response direction $T\hat{u}$. Thus, in the ideal rank-1 limit, shifted CLR does not by itself break the covariance–response alignment that underlies CIPHER. It changes the coordinate representation of the dominant mode, but both the response and covariance are changed in the same way.

This analysis also provides a simple interpretation of why normalization can reduce CIPHER's performance. The raw response direction $\hat{u}$ can contain both an absolute-expression component, which changes raw expression amplitudes, and a relative-expression component, which changes expression composition after total-count or scale changes are removed. A broad program of the form $\hat{u}_i \sim g_i x_i^*$ is an absolute-expression component: it need not be a trivial high-expression artifact if the gene-specific weights $g_i$ encode a real biological program. Frequency normalization, library-size correction, and shifted CLR attenuate such absolute-expression components and retain only the relative-expression part $T\hat{u}$. Therefore, comparing CIPHER in raw and normalized coordinates can help decompose the dominant mode into two components. If normalized counts remain predictive, the mode has a relative-expression component; if raw counts are much more predictive, the dominant mode also has a substantial absolute-expression component.

Consistent with this interpretation, CIPHER remains more predictive than shuffled controls after shifted CLR and related normalizations, but its performance is reduced relative to raw counts (Fig. S13, S16). Together with the persistence of predictability after removing highly expressed genes, this argues against the simplest artifacts based on a few large-count genes or sequencing-depth variation. Instead, the results suggest that CIPHER's signal comes from a broad biological mode with both relative and absolute-expression components, most visible in raw expression coordinates rather than in purely compositional coordinates.

#### 1.8 Analytical expression for CIPHER's prediction quality

CIPHER fits one scalar, so one measure of its accuracy for a knockdown of gene $j$ is the squared cosine between the off-diagonal covariance column and the response, also referred to the $R^2$ for the fit between CIPHER predictions and observations. Writing the column as $\Sigma_{ij} = a_j \hat{u}_i + d_j \hat{w}_i$ for $i \neq j$, with $a_j = b\hat{u}_j + d\hat{w}_j$ the coefficient of $\hat{u}_i$ and $d_j = d\hat{u}_j$ the coefficient of $\hat{w}_i$, and using the normalization conditions $\sum_i \hat{u}_i^2 = \sum_i \hat{w}_i^2 = 1$ and $\sum_i \hat{u}_i \hat{w}_i = \kappa$, we get

$$R_j^2 = \frac{(a_j + d_j\kappa)^2}{a_j^2 + d_j^2 + 2a_j d_j \kappa}. \tag{S48}$$

For a gene with $\hat{u}_j \approx \hat{w}_j$ (taken for simplicity), the relative contamination is $\rho = d_j/a_j = 1/(1+\eta)$, and this reduces to

$$1 - R^2 = \frac{\rho^2(1-\kappa^2)}{(1+\rho\kappa)^2 + \rho^2(1-\kappa^2)}, \qquad \rho = \frac{1}{1+\eta}. \tag{S49}$$

This expression shows that two quantities compete to set CIPHER's accuracy: the asymmetry $\kappa$ between drivers and responders, which amplifies a secondary direction along which perturbations respond, and the recycling number $\eta$, which suppresses it. We can see this most clearly in two limits. Near a cell-fate transition or bifurcation, $\eta \to \infty$, the secondary direction is washed out and $1 - R^2 \approx (1-\kappa^2)/\eta^2$, and CIPHER works exactly. Far from it, $\eta \to 0$, recycling no longer helps and $R^2 \approx (1+\kappa)/2$, set by the asymmetry alone. In the special case $\kappa = 1$ the secondary direction actually aligns with the primary direction $\hat{u}$, and $R^2 = 1$ for every $\eta$. Below the threshold $c\kappa > \sigma$, finally, there is no outlier and hence no structured off-diagonal covariance, so $R^2 \sim 1/N$.

Thus a high $R^2$ requires two things. First, a mode gap, so that a single outlier along $\hat{u}$ exists and dominates the covariance. Second, given that gap, proximity to a bifurcation through a large $\eta$, which suppresses the asymmetry contaminant. Both conditions hold even for strongly directed networks with $\kappa$ well below one, so a high $R^2$ does not imply $\kappa \approx 1$: a large $\eta$ drives $R^2 \to 1$ at any $\kappa$. CIPHER working well therefore suggests that there might a dominant soft mode in the GRN.

#### 1.9 Predictions of the theory match CIPHER's observed trends

**Differentiating cells perform best.** Our theory predicts that $\Delta R^2$ is highest when $\eta$ is large, i.e., when $\gamma \approx c\kappa$ and cells are near a fate transition. As cells approaches a fate transition, the leading eigenvalue approaches the stability edge, its susceptibility diverges, and a single mode dominates the response. Consistent with this, we find that CIPHER shows highest $\Delta R^2$ in differentiating stem cell systems such as Tian19 (Fig. 6G; Fig. S12A,B).

**Shuffling counts across cells destroys the prediction.** Our prediction relies entirely on the off-diagonal structure of the covariance. Shuffling each gene's counts independently across cells preserves every gene's marginal distribution but the $\Delta R^2$ collapses to $O(1/N)$. We indeed find that CIPHER's predictability drops drastically after shuffling, confirming that the prediction comes from gene–gene correlations and not from any single-gene statistics.

**Covariance transfers within a cell type but not across.** In our model the rank-1 mode $c\,\hat{u}\hat{w}^\top$ represents a cell-type-specific regulatory program. Two experiments on the same cell type therefore share the same mode, and hence the same response direction $\hat{u}$, so a covariance measured in one should predict perturbation responses in the other. A different cell type has a different mode and a different $\hat{u}$, so covariance should not transfer across cell types. Consistent with this, we find that using CIPHER predictability transfers from one dataset to another as long as it is from the same cell type.

**Gene–gene correlations should typically be less predictive than gene–gene covariances.** CIPHER uses the unnormalized covariance rather than the correlation. Normalizing to a correlation divides gene $i$ by $\sqrt{\Sigma_{ii}}$, which reweights the covariance column away from $\hat{u}$, and the more unevenly genes vary in expression, the larger this distortion. The proportionality between covariance column and response holds only without this rescaling, so we expect correlation to predict perturbation response worse than covariance.

**Low participation ratio accompanies high $\Delta R^2$.** The participation ratio measures the effective dimensionality of the response. As the cell approaches a bifurcation, the covariance sharpens toward the single $\hat{u}$ outlier, which raises $\Delta R^2$ and lowers the participation ratio together; both follow from the same mode gap. When instead several modes sit above threshold, the response spreads across them, the participation ratio rises, and a single scalar can no longer capture all of them, so $\Delta R^2$ falls. CIPHER reports exactly this anti-correlation between participation ratio and $\Delta R^2$ across its datasets.

**A single highly expressed gene cannot produce a spurious prediction by itself.** Suppose one gene is expressed far above the others. Could it dominate the covariance and inflate $\Delta R^2$ on its own? A single gene's expression magnitude is a property of its marginal distribution, which shuffling preserves while destroying correlations. If the prediction rested on that one gene's expression level, it would survive shuffling — but it does not, since shuffling collapses $\Delta R^2$. We conclude that the prediction comes from the correlation structure across genes, not from any single gene's expression. Conversely, in systems with no dominant mode, where $c\kappa < \sigma$ and there is no mode gap in the spectrum of $K$, our theory predicts that CIPHER should fail, $\Delta R^2 \sim 1/N$.

##### 1.10 Simulated GRNs with minimal frustration show approximate rank-1 collective mode

Let the directed network be described by the continuous-time ODE system:

$$\dot{\vec{x_i}} = F_i(\vec{\mathbf{x}}) = h \prod_{j \in U(i)} \phi_i(x_j) - \gamma x_i, \tag{S50}$$

where $x_i$ represents the concentration of transcription factor $i$, $\gamma$ is the uniform degradation rate, and $h$ is the basal production rate of gene $i$.

The term $\phi_i(x_j)$ is the regulation function defining the influence of an upstream node $j \in \mathcal{U}(i)$ on the target node $i$. This is modeled using a Shifted-Hill function based on fractional binding site occupancy:

$$\phi_i(x_j) = 1 + (\beta_{ij} - 1)\frac{\kappa_a x_j^n}{1 + \kappa_a x_j^n} \tag{S51}$$

We define the kinetic parameters dictating this regulation are defined as follows. $\kappa_a$ is the binding affinity (association constant) of the transcription factor to the target gene's promoter region. $n$ is the Hill coefficient, which dictates the cooperativity of the binding interaction (e.g., dimer formation). $\beta_{ij}$ is the fold change modifier to the basal production rate caused by the bound transcription factor. Node $j$ acts as an activator if $\beta_{ij} > 1$, and as a repressor if $0 \leq \beta_{ij} < 1$.

In Fig. S1, we generate the frustration free network by first assigning an arbitrary vector of $\pm 1$s to all genes and setting the interaction between them to be activating if the two nodes it connects have the same sign, and inhibiting otherwise. This network has assigned "spins" for each node (gene) and is guaranteed to be free of frustration [14]. We measure frustration through the metric $\rho$ defined as the fraction of edges (interactions) that deviate from this case; thus frustration-free networks have $\rho = 0$. Starting from this network, we then increase frustration by flipping the signs of a fraction $\rho$ of edges. Kinetic parameters were chosen as: binding affinity $\kappa_a = 1$, Hill coefficient $n = 2$, degradation rate $\gamma = 60$, and basal production $h = 5$. $\beta_{ij}$ were sampled from identical Gaussian distributions with means of 1.5 for activators and 0.5 for repressors and standard deviation 0.1 for both.

#### 2 Application to synthetic gene regulatory networks

##### 2.1 Random linear network

Having explained the theoretical foundations of the CIPHER framework, we now test it on a number of synthetic gene regulatory networks. We first consider the case of a linear system with random regulatory structure. That is, around a stationary fixed point centered at the origin, so that $\mathbf{x}^* = 0$ and $\delta\mathbf{x} = \mathbf{x}$, the Jacobian becomes constant and we parameterize it as $-J \equiv A = \mathbf{1} + \epsilon\mathbf{K}$, where $\mathbf{K}$ is an off-diagonal matrix of interactions with elements drawn from a standard normal distribution. As Robert May showed in his seminal paper on the stability of random ecosystems, $A$ is almost surely stable when the dimension of the system $N$-genes satisfies the following criteria from random matrix theory: $\epsilon < \epsilon_c = 1/\sqrt{N}$. As the network of genetic interactions becomes more and more dense, the coupling strength approaches its critical value and the smallest eigenvalue of $A \to 0$. The system bifurcates at this critical coupling, becoming globally unstable. We simulate this system using an Euler-Maryama integrator with timestep $dt = 0.1$. For the correlations and linear response estimates in FIG 2, we use $N = 300$ genes and integrate from the stationary state $\delta\mathbf{X} = 0$ for a total time of $T = 300$ with a constant diffusion coefficient $D = 0.5$ for all genes. For this system, the critical coupling is $\epsilon_c = 1/\sqrt{N} = 1/20$, and we simulate one system at $\epsilon = 0.95\epsilon_c$, one system at criticality $\epsilon = \epsilon_c$ and one above the threshold $\epsilon = 1.05\epsilon_c$. Linear response estimates are averaged over 500 independent simulation runs.

The emergence of gene teams past criticality is a direct result of the dynamics begin projected (according to center manifold theory [18]) onto the null-space of $A$, i.e. the direction with zero or negative eigenvalue [19]. This effective dimensionality reduction is not a result of gene teams, but the approach to and past criticality. This insight sheds light on the natural emergence of gene teams in the vicinity of a phenotypic transition; as the Jacobian becomes unstable as it would when passing through a short lived unstable fixed point, or barrier, in phase-space, mutual activation within large groups of genes and inhibition between groups becomes inevitable. This along with the fact that such gene teams have recently been posited to induce low-dimensional phenotypic structure in the epithelial to mesenchymal transition is why we study an explicit teams system below.

#### 2.2 Nonlinear Hill function network

We now move on to test CIPHER on a prototypical nonlinear dynamical gene expression system comprised of a network of hill functions [20]. Specifically, the deterministic force on the $i^{th}$ gene $x_i$ is $\mathbf{F}_i(\mathbf{x}) = -\gamma x_i + \sum_j G_{ij}x_j^n/(K^n + x_j^n)$. We apply an additional constant external force to the first gene with magnitude $u$ and compare the true average response (from solving the stationary fixed point equations $\mathbf{F}_i(\langle\mathbf{x}\rangle_\mathbf{u}) + \mathbf{u} = 0$ and $\mathbf{F}_i(\langle\mathbf{x}\rangle_0) = 0$ to that predicted by linear response (S27), where again $J$ is evaluated at the unperturbed fixed point $\langle\mathbf{x}\rangle_0$. For this system, we calculate the original fixed point (with $\mathbf{u} = 0$) and the Jacobian evaluated at that fixed point, and relate it to the new driven fixed point after perturbation. We perform these calculations using deterministic gradient flow $\dot{\mathbf{x}}_i = \mathbf{F}_i$ (towards the fixed point) for a maximum of 2000 iterations with time-step $dt = 0.01$, stopping if the maximum change in $\mathbf{x}$ over an iteration is less than a fixed tolerance of $10^{-12}$. We consider a relatively small system $N = 10$ to sweep over many parameters and fix $K = 10, \gamma = 1, G_{ij} \sim \mathcal{N}(1, 0.1)$, starting from an initial guess where all genes are at abundance $N\langle G_{ij}\rangle/\gamma = 10$. When sweeping over the other parameters, we fix the Hill coefficient to $n = 4$.

For this system, we can explain the sudden jumps in the linear response error as a function of the system parameters by examining its stability. We assume a homogeneous fixed point at zero and follow what happens when we perturb around it by an amount $\delta x$, so that, averaging over the noise and assuming fixed $G_{ij} = G$,

$$\delta\dot{x} = -\gamma\delta x + \frac{GN\delta x^n}{K^n + \delta x^n}. \tag{S52}$$

The system goes from stable to perturbation to unstable when $\delta\dot{x} = 0$, so that

$$\frac{G^*N}{\gamma^*} = \frac{\delta x^{n-1}}{(K^*)^n + \delta x^n}. \tag{S53}$$

Solving for $G^*, \gamma^*$, or $K^*$ yield the critical parameter values past which the system has finite average gene expression and the linear response error estimates jump.

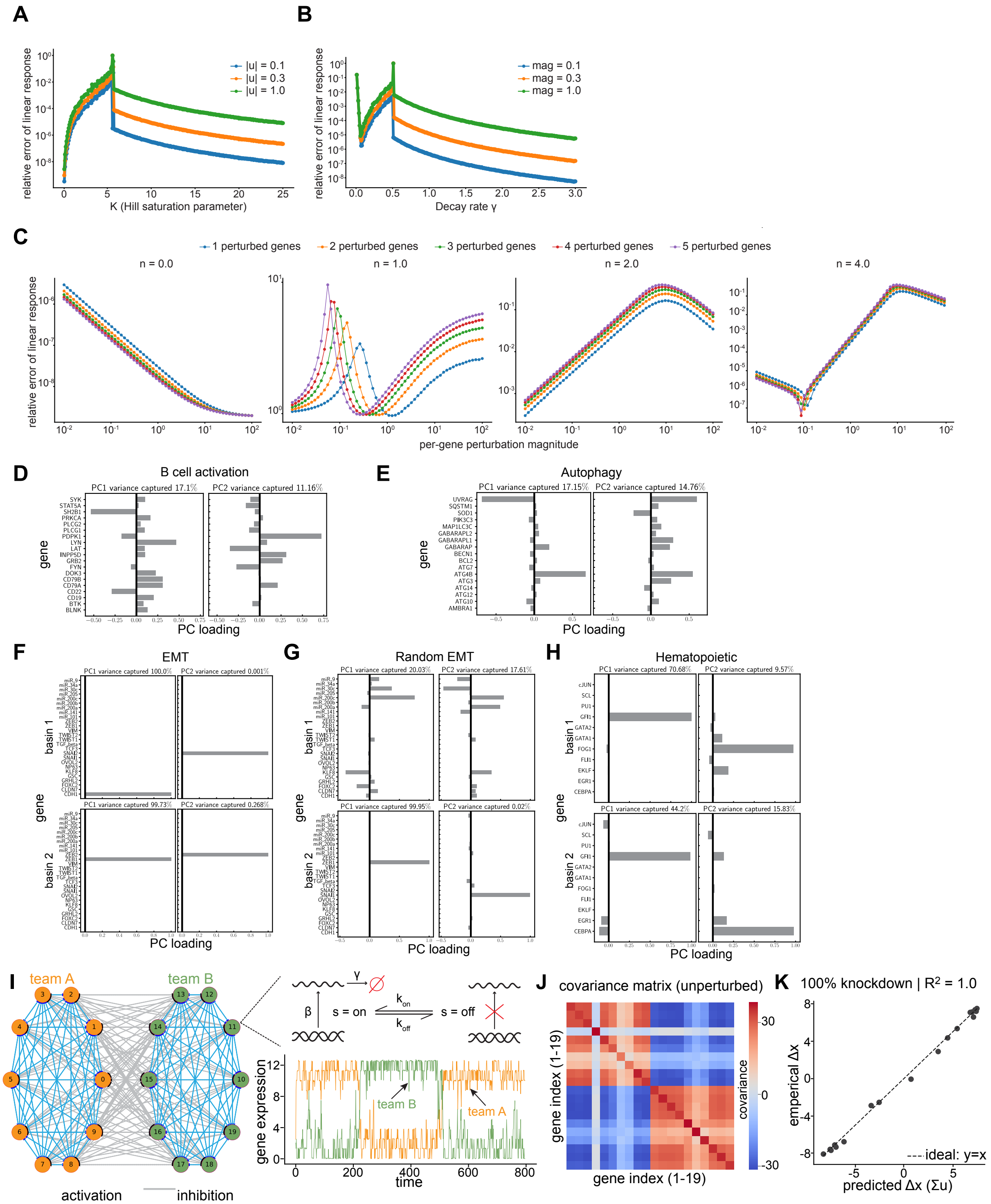


**Fig. S3**: **Additional numerical toy examples.** A) Relative error in linear response as a function of Hill saturation parameter K for three perturbation magnitudes. B) Relative error in linear response as a function of decay rate for three perturbation magnitudes. C) Relative error in linear response with multiple genes perturbed. For increasing Hill coefficient (left to right) we show relative error as a function of perturbation magnitude for 1-5 genes perturbed with equal magnitude. Relative error in linear response (single gene perturbed) as a function of Hill saturation parameter K for three perturbation magnitudes. Each point represents a single simulation run and lines guide the eye. D)-H) First 2 principal components for the biologically informed networks in Fig. 2. I)Teams network illustration J) Teams network covariance matrix K) Teams network response upon single-gene knockdown.

#### 2.3 Testing CIPHER using numerical simulations

##### 2.3.1 Shifted Hill regulation

For all biological networks we probed, regulatory interactions are modeled as multiplicative shifted Hill functions following the RACIPE framework [21, 22]. For a target gene $i$ in the gene regulatory network (GRN), with activating regulators indexed by $\mathcal{R}_i^+$ and inhibiting regulators indexed by $\mathcal{R}_i^-$ (both determined by the network topology), the governing ordinary differential equation is

$$\dot{y}_i \;=\; g_i \prod_{j\in\mathcal{R}_i^+} H^s(y_j;\Theta_{ij},n_{ij},\lambda_{ij}^+) \prod_{j\in\mathcal{R}_i^-} H^s(y_j;\Theta_{ij},n_{ij},\lambda_{ij}^-) \;-\; k_i\, y_i, \tag{S54}$$

where $g_i$ is the production rate and $k_i$ the degradation rate of gene $i$. The shifted Hill function $H^s$ takes a single functional form for both activation and inhibition; the sign of regulation is encoded entirely in the fold-change parameter $\lambda_{ij}$:

$$H^s(y;\Theta,n,\lambda) \;=\; \lambda + (1-\lambda)\frac{1}{1+(y/\Theta)^n}. \tag{S55}$$

Here $\Theta$ is the half-saturation threshold, $n$ is the Hill coefficient, and $\lambda$ is the fold-change. At $y = 0$ we have $H^s = 1$; at saturation ($y \gg \Theta$) we have $H^s \to \lambda$. Thus $\lambda > 1$ encodes activation (expression increases at saturation) and $\lambda < 1$ encodes inhibition.

##### 2.3.2 RACIPE parameter sampling

For each parameter set $\{\mathbf{g},\mathbf{k},\boldsymbol{\Theta},\lambda,\mathbf{n}\}$, Equation (S54) is integrated from multiple random initial conditions to identify stable fixed points. The parameter vector for a network with $N$ nodes and $E$ edges has $2N + 3E$ entries: a production rate $g_i$ and degradation rate $k_i$ for each gene, and $(\Theta_{ij}, n_{ij}, \lambda_{ij})$ for each edge. Sampling an ensemble of parameter sets is intended to represent the variability of steady expression states arising from biological and cellular heterogeneity. Default sampling ranges follow Huang et al. [21], drawn as continuous uniform distributions as implemented in GRiNS [22]. In the RACIPE framework the threshold ranges for $\Theta_{ij}$ are not preset but are estimated numerically for each network to satisfy the half-functional rule: across the model ensemble, each regulator should have roughly a 50% chance of being functional (i.e., of its expression exceeding the threshold) [21]. We solve the GRN equations with the RACIPE framework using the GPU implementation GRiNS [22].

##### 2.3.3 Steady-state filtering

For each (parameter set, initial condition) pair, the ODE is integrated to $t = 800$ using Diffrax's adaptive Dopri5 solver [23] as implemented in GRiNS [22], with relative tolerance $10^{-6}$ and absolute tolerance $10^{-7}$. A candidate steady state $\mathbf{y}^*$ is accepted if the Jacobian of Equation (S54) evaluated at $\mathbf{y}^*$ has eigenvalues with strictly negative real parts (i.e., $\mathbf{y}^*$ is a stable fixed point).

##### 2.3.4 Deduplication of steady states

Different (parameter set, initial condition) pairs may converge to the same steady state; leaving these duplicates in place would bias the steady-state statistics toward more easily reached attractors. We therefore deduplicate steady states by rounding the raw steady-state values to $d = 2$ decimal places and removing repeats. Deduplication yields an unbiased set of distinct attractors from which basins associated with the network topology can be constructed.

##### 2.3.5 Clustering via $K$-means with biological-meaningfulness criteria

After deduplication, steady states are $z$-scored per gene and projected onto the leading $n_{\mathrm{PC}} = 5$ principal components in $\log(1 + p)$ space. For each candidate cluster number $k \in \{2, \ldots, K_{\max}\}$ we run $K$-means and evaluate three criteria:

1. **Silhouette score**: $s(k) \geq 0.25$, where $s(k)$ is the mean silhouette coefficient in PC space.
2. **Balance**: the smallest cluster contains at least 5% of the distinct attractors. This rejects splits driven by isolated outliers.
3. **Differentiation**: for at least one gene, the $\log(1+p)$ difference between two cluster means exceeds 2 (a four-fold change). This rejects splits along noise dimensions.

The smallest $k$ satisfying all three criteria is selected. If no $k > 1$ passes, the network is taken to have a single basin ($k = 1$).

###### 2.3.6 Knockdown perturbation protocol

For each basin of a biological network we test CIPHER on 1200 of its steady states. For each such steady state and each gene $i$ in the network, we perform a fractional knockdown by reducing the production rate,

$$g_i \to f\, g_i, \tag{S56}$$

where $f = 0.2$ is the knockdown fraction, corresponding to an 80% reduction (20% residual production). Starting from the unperturbed steady state $\mathbf{y}^*_\theta$ as the initial condition, the ODEs are integrated with the perturbed parameter vector to the new fixed point,

$$\mathbf{y}^*_{\theta,g} = \lim_{t\to\infty} \mathbf{y}(t;\ \mathbf{y}(0) = \mathbf{y}^*_\theta,\ \theta_{\text{pert}})\,. \tag{S57}$$

We integrate to $t_{\max} = 1000$ using the same solver and tolerances as in Section 2.3. Perturbed trajectories that fail to reach a stable fixed point (non-convergence, or a Jacobian with any eigenvalue of non-negative real part) are excluded. The observed perturbation response is

$$\boldsymbol{\Delta}\mathbf{y}^{\text{obs}}_{\theta,g} \;=\; \mathbf{y}^*_{\theta,g} - \mathbf{y}^*_\theta, \tag{S58}$$

computed using raw values. Responses whose raw-coordinate norm $\|\mathbf{y}^*_{\theta,g} - \mathbf{y}^*_\theta\|$ falls below a threshold, i.e. 10.0 are discarded, as their direction is dominated by numerical noise rather than a resolvable biological response.

###### 2.3.7 The CIPHER prediction and its evaluation

CIPHER predicts that the response to a partial knockdown of gene $g$ is proportional to the corresponding column of the within-basin gene–gene covariance matrix,

$$\boldsymbol{\Delta}\mathbf{y}^{\text{CIPHER}}_g \;\propto\; \boldsymbol{\Sigma}_{:,g}, \tag{S59}$$

where $\boldsymbol{\Sigma}$ is the covariance of the (deduplicated) steady states across the parameter ensemble within a basin, computed using raw values. We stress that $\boldsymbol{\Sigma}$ is the ensemble covariance of steady states and is estimated independently of the perturbation responses.

To quantify agreement between the predicted and observed responses we compute, for each steady state $\theta$ and each knocked-down gene $g$, the Pearson correlation between the observed response and the predicted covariance column:

$$r_{\theta,g} = \frac{\sum_i \left(\Delta y_i^{\text{obs}} - \overline{\Delta y^{\text{obs}}}\right)\left(\Sigma_{i,g} - \overline{\Sigma_{:,g}}\right)}{\sqrt{\sum_i \left(\Delta y_i^{\text{obs}} - \overline{\Delta y^{\text{obs}}}\right)^2}\ \sqrt{\sum_i \left(\Sigma_{i,g} - \overline{\Sigma_{:,g}}\right)^2}}, \tag{S60}$$

where the sum runs over all genes $i$.

Because both $\boldsymbol{\Delta}\mathbf{y}^{\text{obs}}_g$ and $\boldsymbol{\Sigma}_{:,g}$ generally have nonzero means—knockdown responses typically shift many genes in a coherent direction—we use the mean-centered (Pearson) correlation, which removes this shared offset and matches the convention used in single-cell perturbation studies.

For each gene $g$ we summarize performance as the median of $r_{\theta,g}$ over all tested steady states $\theta$ that pass the filtering criteria. Per-steady-state correlations are computed individually and then aggregated, rather than averaging the response vectors before correlating; this avoids cancellation between steady states whose responses point in opposite absolute directions.

Finally, we performed a randomized-covariance control or null test. For each basin we generated 1000 surrogate covariance matrices by permuting the off-diagonal entries of $\mathbf{\Sigma}$ among themselves while leaving the diagonal unchanged. Each surrogate was substituted for $\mathbf{\Sigma}$ in Eq. (S59) and scored against the same observed knockdown responses using Eq. (S60), yielding for each gene a median $r$ across steady states exactly as for the true covariance. We calculate, per gene, the median of these 1000 surrogate values across permutations. Because both the diagonal and the off-diagonal values are preserved for all permutations, any loss of correlation relative to the true $\mathbf{\Sigma}$ is attributable specifically to the arrangement of off-diagonal covariances rather than to their magnitudes or to gene-level variances.

#### 2.4 Boolean dynamics and network frustration

For each biological network we also performed Boolean dynamics. Each node $s_i$ was assigned a random initial value in $\{-1, +1\}$ and updated at discrete time steps according to

$$s_i(t+1) = \begin{cases} +1 & \text{if } \sum_j J_{ij}\, s_j(t) > 0, \\ -1 & \text{if } \sum_j J_{ij}\, s_j(t) < 0, \\ s_i(t) & \text{if } \sum_j J_{ij}\, s_j(t) = 0, \end{cases} \tag{S61}$$

where $J_{ij} = +1$ for an activating edge and $J_{ij} = -1$ for an inhibiting edge, as defined by the network topology. The update is asynchronous in which a single randomly selected node is updated per step. Each initial condition was evolved for 700 steps, and 100 independent initial conditions were used per network.

For a given final state, the frustration is the fraction of edges that are unsatisfied, i.e., edges for which $J_{ij}\, s_i\, s_j < 0$ [24]. We take the network's frustration to be the mean frustration over all sampled initial conditions.

To generate a randomized EMT network we followed the edge-swapping procedure of Tripathi et al. [14]. Pairs of edges are selected at random and their targets exchanged, so that edges $(i \to j)$ and $(k \to l)$ become $(i \to l)$ and $(k \to j)$. Each edge retains its activating or suppressing character, and because only targets are exchanged, the in- and out-degree of every node is preserved exactly. The randomized networks therefore share the degree sequence and the activation-to-suppression ratio of the original network, and differ from it only in the wiring of the edges. We performed at least 50 swaps per network, by which point the frustration value had plateaued, indicating that the topology had decorrelated from the original.

#### 2.5 Teams network

The last synthetic system we explore here is a stochastic hybrid model of team dynamics, where genes on the same team mutually activate each other and there is inhibition between genes on different teams. Each gene has a promoter that transitions from off $s = 0$ to on $s = 1$, and vice versa, with rates $k_{on}$ and $k_{off}$. These rates take in input from all the other genes in the system which is modulated by a hill function. That is $k_{on/off} = 1/(1+(K/\sum_j B_{ij}^{on/off} x_j)^n)$ where $B^{on/off}$ is a matrix of activating/inhibitory interactions, respectively. The force on each gene is then $\mathbf{F}_i(\mathbf{x}) = \beta s_i - \gamma x_i$ where $\beta$ and $\gamma$ are transcription and decay rates. This nonlinear model features collective transitions wherein all the genes on a team will cooperatively transition from off to on while the genes on the other team transition from on to off. Even in such a complex simulated system, gene expression changes upon knock-out of a single gene (setting $\beta = 0$ for some gene $i$) are well captured by linear response. For our simulations, we fix $B_{ij}^{on} = 0.3(1+\mathcal{N}(0, 0.45))$, $B_{ij}^{off} = 0.25(1 + \mathcal{N}(0, 0.45))$, $K = 25$ and $n = 2$. We set $\beta = 50$ and $\gamma = 4$ and simulate $N = 20$ genes (10 on each team) for a total time of $T = 2 \times 10^6$ with timestep $dt = 0.1$. We set the sparsity to $S = 0.7$ by randomly setting elements of $B^{on/off}$ to zero with probability $S$. At the end of the long simulation, we calculate the gene-gene covariance and the optimal single-gene perturbation according to Eq. (S66) after setting the production rate of the first gene to zero, knocking it out.

### 3 The forward problem: Transcriptome-wide linear response and how to use it

We first state the main statement of CIPHER's linear response and briefly cover how to use it on real data before showing general arguments and examples in support of it. Denoting the state of the unperturbed system as $\mathbf{x} = (x_1, x_2, ...)$, the perturbation strength as $\mathbf{u}$ and the covariance as $\Sigma$ so that $\Sigma_{ij} = cov(x_i, x_j)$, our main result is

$$\langle \mathbf{x} \rangle_{\mathbf{u}} = \langle \mathbf{x} \rangle_0 + \Sigma \mathbf{u}. \tag{S62}$$

or

$$\langle \Delta \mathbf{x}_i \rangle = \sum_j \langle \delta \mathbf{x}_i \delta \mathbf{x}_j \rangle_0 \mathbf{u}_j. \tag{S63}$$

Here, we briefly describe how we use this linear response relationship to apply it to real data.

#### 3.1 Fitting single-gene response

We can judge how well correlations explain the expression difference $\Delta X$ by computing the coefficient of determination

$$\Delta R^2 = 1 - \frac{||\Delta X - \Sigma \mathbf{u}||^2}{||\Delta X||^2} \leq 1. \tag{S64}$$

As $\Delta R^2$ approaches 1, the correlations perfectly transport the initial vector of gene expression $\langle \mathbf{x} \rangle_0$ to the final measured target $\langle \mathbf{x} \rangle_{\mathbf{u}}$. When $\Delta R^2 > 0.5$, for example, the initial distance between the control and perturbed gene expression has been halved.

If the cellular perturbations causing phenotypic change are known, as they are in Perturb-seq experiments, wherein it is typical for a single gene to be upregulated or knocked down per cell by CRISPRa or CRISPRi respectively, we can directly test the extent to which linear response to that single-gene perturbation effects the expected expression change. That is, for any known single-gene perturbation $i$, we can solve for the optimal $u_i$ with which to force the system towards the terminal state $\langle \mathbf{x} \rangle_{\mathbf{u}}$. Setting all other elements of $\mathbf{u}$ equal to zero, the $u_i$ that solves the optimization problem

$$\mathbf{u}_i^* = \text{argmin}_{\mathbf{u}} ||\Delta X - \Sigma \mathbf{u}||^2, \Delta X = \langle \mathbf{x} \rangle_{\mathbf{u}} - \langle \mathbf{x} \rangle_0 \tag{S65}$$

is

$$\mathbf{u}_i^* = (0, ..., u_i^*, ..., 0), \; u_i^* = \frac{\Sigma_i^T \cdot \Delta X}{\Sigma_i^T \cdot \Sigma_i} \tag{S66}$$

where $\Sigma_i$ is the ith column of $\Sigma$. We use such optimal single-gene solutions in the 'forward' problem below and can derive it explicitly as follows. Assuming we know which gene is perturbed we set all other elements of $\mathbf{u}$ to zero. Now, $\Sigma \mathbf{u} = \Sigma_{1i} u_i + ... + \Sigma_{Ni} u_i \equiv \Sigma_i u_i$ where $\Sigma_i$ is the $i^{th}$ column of $\Sigma$. This means that the optimization in Eq. (S65) reduces to

$$\mathbf{u}_i^* = \text{argmin}_{u_i} ||\Delta X - \Sigma_i u_i||^2 \tag{S67}$$

which implies

$$\partial_{u_i^*} (\Sigma_i^T \; \Sigma_i u_i^2 - 2 \Sigma_i^T \Delta X u_i) = 0. \tag{S68}$$

Solving this equation for the scalar $u_i$ gives the desired result. We note that the effects of higher order multi-gene perturbations can be solved for with a similar though higher dimensional linear optimization framework.

In practice, we compute the optimal single gene perturbation $\mathbf{u}_i^*$ from Eq. (S65) for each single and double-gene perturbation in each dataset, after cleaning it (see Data Processing below). We then use $\mathbf{u}_i^*$ to calculate an $\Delta R^2$ value for 1) the real $\Sigma$ matrix, 2) the 'mean-field' matrix, shuffled only over cells, not genes, and 3) the fully-'shuffled' matrix. First we regularize the denominator in both the $\Delta R^2$ expression and the expression for $\mathbf{u}_i^*$ by adding a small number $10^{-8}$ to avoid division by zero. In order to suppress spurious small and noisy correlations from artificially driving down $\Delta R^2$, in the sum what uninteresting case that the expression of a given gene does not change ($\Delta \mathbf{x}_i = 0$), we only compute $\Delta R^2$ over genes that have nonzero response to the perturbation.

When comparing distributions and means of $\Delta R^2$ across datasets, we calculate p-values using Kolmogorov-Smirnoff tests and Wilcoxon signed rank t-tests (one-sided), as implemented in sciPy, respectively.

#### 3.2 Visualization of CIPHER-predicted perturbation fields

To visualize how CIPHER-predicted transcriptome-wide responses propagate across heterogeneous control-cell states, we generated counterfactual displacement fields on a shared UMAP embedding. Briefly, we selected the 2,000 most variable genes among control cells (forcing inclusion of perturbed targets where necessary) and fit UMAP directly on the resulting raw-count expression vectors without normalization, log transformation, or principal-component analysis.

For a perturbation targeting gene $g$, CIPHER predicts the transcriptome-wide mean response

$$\Delta x = \Sigma u = u_g \Sigma_{:g}, \tag{S69}$$

where $\Sigma$ is the control-cell covariance matrix and $u_g$ is the inferred perturbation amplitude. For every control cell with expression vector $x_0$, we generated a counterfactual perturbed state

$$x_0^{\mathrm{cf}} = x_0 + \Sigma_{:g} u_g. \tag{S70}$$

Negative values resulting from this additive approximation were truncated to zero before projection. Both the original and counterfactual cells were then embedded using the same pre-fit UMAP transformation

$$U : \mathbb{R}^G \to \mathbb{R}^2, \tag{S71}$$

giving the predicted displacement

$$v_g(x_0) = U(x_0 + \Sigma_{:g} u_g) - U(x_0). \tag{S72}$$

Eq. ((S72)) represents the central quantity visualized in the blue vector fields. Importantly, although every control cell receives the same transcriptomic perturbation $\Sigma_{:g} u_g$, the displacement in UMAP space generally depends on the initial state because the embedding is nonlinear. For sufficiently small perturbations,

$$U(x_0 + \Sigma_{:g} u_g) - U(x_0) \approx J_U(x_0) \Sigma_{:g} u_g, \tag{S73}$$

where $J_U(x_0)$ is the local Jacobian of the UMAP mapping. Thus, the displayed vector field is the nonlinear pushforward of the CIPHER response through the learned manifold geometry.

To reduce noise, the control embedding was partitioned into a regular grid, and displacement vectors were averaged within each occupied bin. If $B_k$ denotes the set of control cells in bin $k$, the displayed vector is

$$\bar{v}_k = \frac{1}{|B_k|} \sum_{i \in B_k} \left[ U(x_{0,i} + \Sigma_{:g} u_g) - U(x_{0,i}) \right]. \tag{S74}$$

Equivalently, the arrows estimate the conditional expectation

$$v_g(y) = \mathbb{E} \left[ U(x_0 + \Sigma_{:g} u_g) - U(x_0) \,|\, U(x_0) \approx y \right]. \tag{S75}$$

These vector fields should not be interpreted as RNA velocity or inferred temporal dynamics. Rather, they visualize the predicted displacement of cells occupying different regions of the control-state manifold under a common transcriptome-wide perturbation. Across diverse datasets, for high-performing CIPHER perturbations, the predicted fields consistently pointed toward the observed perturbed populations and frequently exhibited smooth curved trajectories. This behavior reflects the geometry of the nonlinear embedding rather than nonlinear response dynamics: a globally linear perturbation in gene-expression space becomes state dependent after projection through the nonlinear map $U$.

#### 3.3 Covariance versus correlation formulations of the forward problem

CIPHER predicts the transcriptome-wide response to a perturbation using the linear-response model

$$\Delta x \approx \Sigma u, \tag{S76}$$

where $\Sigma$ is the control-cell covariance matrix and $u$ denotes the perturbation vector. An alternative representation uses the correlation matrix,

$$R = D^{-1}\Sigma D^{-1}, \tag{S77}$$

where $D = \mathrm{diag}(\sigma_1, \ldots, \sigma_p)$ contains the gene standard deviations. Defining standardized expression coordinates $\Delta z = D^{-1}\Delta x$ and transformed perturbation amplitudes $v = Du$ gives the equivalent linear-response model

$$\Delta z \approx Rv. \tag{S78}$$

Although these expressions describe the same linear-response relationship in different coordinate systems, they lead to different optimization problems depending on the metric used to measure prediction error. The covariance formulation minimizes residuals in the original expression units,

$$\min_u \|\Delta x - \Sigma u\|_2^2, \tag{S79}$$

where every gene contributes equally in raw expression space. By contrast, performing ordinary least squares directly in correlation coordinates minimizes

$$\min_v \|\Delta z - Rv\|_2^2 = \min_u (\Delta x - \Sigma u)^T D^{-2} (\Delta x - \Sigma u), \tag{S80}$$

which weights each gene inversely by its variance. Consequently, highly variable genes contribute less to the objective, while genes with small intrinsic variance are emphasized.

We compared these two formulations across all Perturb-seq datasets using identical train/test splits and evaluation procedures (Fig. S7A,B). The covariance formulation consistently achieved higher median Pearson correlations between predicted and observed transcriptome-wide responses than the ordinary correlation formulation, both when using the true covariance matrix and when using the mean-field covariance approximation. This result indicates that preserving the natural covariance scale of gene expression yields more accurate forward predictions than standardizing each gene to unit variance before fitting. In practice, the dominant transcriptional responses are carried by high-variance collective modes of the covariance matrix, and down-weighting these modes through correlation normalization reduces predictive performance.

#### 3.4 Pre-processing of datasets

We apply a simple filtering procedure to the raw count matrices from the datasets considered in this study. We begin by loading single-cell perturbation datasets in .h5ad format using ScanPy [25]. We then standardize perturbation names by stripping replicate or guide suffixes (e.g., g1, g2) to extract a base perturbation label. As we only consider single gene perturbations in this study, we discard combinatorial perturbations from all datasets.

To retain biologically meaningful genes and limit the amount of noise in the data due to extremely low expression counts, we apply an average expression filter, keeping only genes with a mean expression above a specified threshold (1 count) and ensuring that all measured perturbed genes are preserved regardless of their expression level. For the perturbations, we retain only those with at least a specified minimum number of cells (100). As certain perturbation prediction methods require gene embeddings for model training, we further filter the dataset to retain only the genes present in the GenePT 3.5 gene embedding set [26]. We then retain only the top 1,000 perturbations by cell count. For each perturbation across these filtered datasets, we randomly sample 10,000 control cells and an even split of training and testing cells. This filtration regime ensures a fair comparison across perturbation prediction methods as all methods train and test on identical datasets.

To compare CIPHER across different preprocessing strategies, we formulate the forward problem in the transformed expression coordinates rather than directly in raw counts. Let $x \in \mathbb{R}^G$ denote the raw count vector for a cell and let

$$y = f(x), \tag{S81}$$

be an element-wise preprocessing transformation. The covariance matrix is then estimated in the transformed space,

$$\Sigma_y = \mathrm{Cov}(y), \tag{S82}$$

and CIPHER predicts perturbation responses according to

$$\Delta y \approx \Sigma_y u. \tag{S83}$$

Thus, each normalization defines a different coordinate system and corresponding covariance matrix while leaving the linear-response formalism unchanged.

For raw counts,

$$f(x) = x. \tag{S84}$$

For log1p normalization,

$$f(x) = \log(1 + x), \tag{S85}$$

while frequency normalization uses relative transcript abundances,

$$f(x) = \frac{x_j}{\sum_j x_j}. \tag{S86}$$

The standard library-size representation is obtained by

$$f(x) = \log\left(1 + 10^4 \frac{x_j}{\sum_j x_j}\right). \tag{S87}$$

Finally, the PFlog transformation is given by,

$$f(x) = \log(x + c) - \frac{1}{G}\sum_{j=1}^{G} \log(x_j + c), \tag{S88}$$

where the pseudocount is chosen as

$$c = \frac{1}{4\alpha}, \tag{S89}$$

with $\alpha$ estimated on the level of dataset from the negative-binomial mean–variance relationship

$$\mathrm{Var}(X) \approx \mu + \alpha\mu^2. \tag{S90}$$

Apart from the preprocessing transformation itself, all train/test splits, covariance estimation procedures, and evaluation metrics were identical across normalization schemes.

#### 3.5 Perturbation filtering for benchmarking

The in-distribution held out test perturbations were subject to further filtering. We iteratively split each perturbation's gene expression change over the control into two independent subsets, drawn randomly without replacement (n=20). For each iteration, we verify that CIPHER's fitting coefficient maintains the same sign, and that the Pearson correlation between splits is greater than 0.85. These criteria serve as proxies for determining whether a perturbation's expression signature is reproducible from the data. Only the perturbations that pass these criteria form the testing set.

##### 3.6 Benchmarking against other methods

We compare CIPHER against a simple linear mean baseline and five published perturbation prediction methods. All methods were trained and evaluated on identical datasets. For all methods excluding CIPHER and linear mean, the datasets are preprocessed by normalizing each cell by total counts over all genes, followed by log normalization. The methods were trained and evaluated individually on each dataset using a compute node with 256 GB of memory and a single NVIDIA H100 GPU.

In the linear mean baseline, the predicted perturbation effect for each gene is simply the mean expression of cells receiving that perturbation in the training set.

We benchmarked the GenePert model [27], which represents each perturbation by its GenePT 3.5 gene embedding. These pretrained gene embeddings effectively encode biological knowledge, especially for gene-gene interactions and gene function. The regression target for each perturbation is the mean expression profile of the training cells for that perturbation. We fit a Ridge regression ($\alpha = 0.1$) mapping perturbation embeddings to these mean expression profiles, so the prediction for a test perturbation comes from applying the fitted model to perturbed gene embedding.

scLAMBDA applies a deep latent variable approach based on variational autoencoders (VAEs), where a low-dimensional embedding with the prior standard normal distribution $z \sim N(0, I_d)$ captures the control state of a cell [28]. Perturbation information is encoded with gene embeddings, $s = E_{perturb}(p) \in \mathbb{R}^d$. These two embedding components are combined and decoded through a decoder network to reconstruct gene expression, reflecting the joint contributions of basal cell variability and perturbation effects to changes in gene expression, while mutual information neural estimation allows for the disentanglement of control and perturbation information. We retain the default hyperparameters in scLAMBDA. The model is trained over 200 training epochs with a latent dimension $d = 30$. The predicted mean expression profile is computed by decoding $z + s$ averaging the predictions for 10,000 synthetic cells.

Unlike scLAMBDA, Scouter conditions on real control cells rather than a sampled latent prior [29], providing a useful contrast for evaluating if a deterministic function of an observed control cell can adequately predict perturbation response without the need for generative sampling. Control cells gene expressions are directly condensed into a latent representation $z$ with dimension $d = 64$ through an encoder. These cell embeddings are then combined with the GenePT gene embedding $s$ for the perturbed gene, and decoded to directly predict the perturbed expression of the control cell. This autoencoder model is trained over 20 epochs, keeping other hyperparameters as default. The predicted mean expression profile for each gene perturbation is computed from averaging the predictions from 300 randomly sampled control cells.

We also benchmarked the scGPT foundation model [30], which contains stacked transformer layers with multi-head attention that jointly learn cell and gene embeddings. Foundation models leverage the knowledge learned from a large corpus of scRNA-seq data and apply it on specific downstream tasks. We fine-tune the model from the pretrained scGPT human checkpoint for each dataset, following the standard perturbation-prediction fine-tuning recipe provided by scGPT, with architecture dimensions matching the configuration of the checkpoint. The foundation model is fine-tuned over 15 epochs, while all other hyperparameters are kept as default. As with Scouter, predictions are averaged over 300 randomly sampled control cells.

In contrast to gene embedding-based methods, GEARS represents genes and perturbations using graph neural networks constructed from prior biological knowledge graphs [31]. By evaluating perturbation predictions with GEARS, we can determine if explicitly encoding gene-gene relationships, rather than treating each gene independently, can improve prediction accuracy. We widen the GO-similarity candidate pool beyond the default essential gene set to also include the measured genes and perturbation targets in each dataset. This allows genes without a match within their own dataset-specific gene panel to be matched against the broader candidate pool. The GEARS model is trained over 10 epochs, while all other hyperparameters are maintained in the default configuration. The predicted mean expression profile for each gene perturbation is computed by averaging the predictions from 300 randomly sampled control cells which are reused across all perturbations within a dataset.

Our benchmarking analysis shows that the computational overhead associated with each method increases with the number of training cells (Fig. S8A-C). We find that simpler methods such as CIPHER and GenePert (Fig. S8D) require significantly less computational resources and training time compared to more complex methods such as scGPT, whilst still providing comparable performance with regards to perturbation prediction (Fig. S8E-G).

### 4 Two-Layer Model for Linear Response in RNA sequencing

We consider a two-layer model in which the observed count vector $X \in \mathbb{Z}_{\geq 0}^{G}$ is generated from a latent biological expression vector $\Lambda \in \mathbb{R}_{\geq 0}^{G}$. In the Poisson case, the model for gene $g$ and the gth element of $X$ is

$$X_g \mid \Lambda_g \sim \text{Poisson}(\Lambda_g), \qquad \Lambda_g \sim F(m, \Sigma). \tag{S91}$$

Here we have suppressed the ensemble labels, but

$$m_{u/0} = \langle \Lambda \rangle_{u/0}, \tag{S92}$$

and

$$\Sigma = \left\langle (\Lambda - \langle \Lambda \rangle_0)\, (\Lambda - \langle \Lambda \rangle_0)^T \right\rangle_0 \tag{S93}$$

is the covariance of latent biological fluctuations in the unperturbed state. We imagine that Eq. (S91) is valid for both perturbed and control ensembles in linear response, and any time we write a second moment below, it is in the control ensemble.

More generally, we replace the Poisson observation layer with a mean-preserving observation distribution with a finite second moment that depends on the mean. For each gene $g$,

$$X_g \mid \Lambda_g \sim p_g(x_g \mid \Lambda_g), \tag{S94}$$

with

$$\langle X_g \mid \Lambda_g \rangle = \Lambda_g, \tag{S95}$$

and

$$\text{Var}(X_g \mid \Lambda_g) = v_g(\Lambda_g) < \infty, \tag{S96}$$

where $v_g(\cdot)$ is the conditional variance function of the observation model **in the control ensemble**.

We assume that linear response holds at the latent biological level:

$$\langle \Lambda \rangle_u - \langle \Lambda \rangle_0 = \Sigma u. \tag{S97}$$

Because the observation layer is mean-preserving,

$$\langle X \rangle_u = \langle \langle X \mid \Lambda \rangle \rangle_u = \langle \Lambda \rangle_u, \tag{S98}$$

and

$$\langle X \rangle_0 = \langle \langle X \mid \Lambda \rangle \rangle_0 = \langle \Lambda \rangle_0. \tag{S99}$$

Therefore,

$$\langle X \rangle_u - \langle X \rangle_0 = \langle \Lambda \rangle_u - \langle \Lambda \rangle_0. \tag{S100}$$

Combining Eqs. (S97) and (S100), we obtain

$$\langle X \rangle_u - \langle X \rangle_0 = \Sigma u. \tag{S101}$$

Thus, for any mean-preserving observation layer with finite second moment, the mean count response is governed by the latent biological covariance $\Sigma$.

The observed count covariance is obtained from the law of total covariance in the control ensemble:

$$\text{Cov}(X) = \text{Cov}(\langle X \mid \Lambda \rangle) + \langle \text{Cov}(X \mid \Lambda) \rangle\,. \tag{S102}$$

Using Eq. (S95),

$$\text{Cov}(\langle X \mid \Lambda \rangle) = \text{Cov}(\Lambda) = \Sigma. \tag{S103}$$

If genes are conditionally independent given $\Lambda$, then

$$\operatorname{Cov}(X \mid \Lambda) = \operatorname{diag}\left(v_1(\Lambda_1), v_2(\Lambda_2), \ldots, v_G(\Lambda_G)\right). \tag{S104}$$

Therefore,

$$\operatorname{Cov}(X) = \Sigma + \left\langle \operatorname{diag}\left(v_1(\Lambda_1), v_2(\Lambda_2), \ldots, v_G(\Lambda_G)\right)\right\rangle. \tag{S105}$$

Equivalently, for $g \neq h$,

$$\operatorname{Cov}(X_g, X_h) = \Sigma_{gh}, \qquad g \neq h, \tag{S106}$$

whereas on the diagonal,

$$\operatorname{Var}(X_g) = \Sigma_{gg} + \langle v_g(\Lambda_g)\rangle. \tag{S107}$$

Thus the off-diagonal count covariance directly estimates the latent biological covariance, while the diagonal contains both biological fluctuations and conditional sampling noise.

##### 4.0.1 Poisson observation layer

For the Poisson observation layer,

$$X_g \mid \Lambda_g \sim \operatorname{Poisson}(\Lambda_g), \tag{S108}$$

so that

$$v_g(\Lambda_g) = \Lambda_g. \tag{S109}$$

Substituting Eq. (S109) into Eq. (S105) gives

$$\operatorname{Cov}(X) = \Sigma + \operatorname{diag}\left(\langle\Lambda\rangle_0\right). \tag{S110}$$

Since

$$\langle\Lambda\rangle_0 = m_0, \tag{S111}$$

we have

$$\operatorname{Cov}(X) = \Sigma + \operatorname{diag}(m_0). \tag{S112}$$

Therefore, in the Poisson case,

$$\Sigma = \operatorname{Cov}(X) - \operatorname{diag}(m_0). \tag{S113}$$

##### 4.0.2 Negative-binomial observation layer

Now consider a negative-binomial observation layer:

$$X_g \mid \Lambda_g \sim \operatorname{NB}(\Lambda_g, \alpha_g), \tag{S114}$$

parameterized by

$$\langle X_g \mid \Lambda_g\rangle = \Lambda_g, \tag{S115}$$

and

$$\operatorname{Var}(X_g \mid \Lambda_g) = \Lambda_g + \alpha_g\Lambda_g^2. \tag{S116}$$

Here $\alpha_g \geq 0$ is a gene-specific overdispersion parameter. The Poisson model is recovered when

$$\alpha_g = 0. \tag{S117}$$

Because the negative-binomial observation layer is mean-preserving, the same linear-response relation holds for observed mean counts:

$$\langle X\rangle_u - \langle X\rangle_0 = \Sigma u. \tag{S118}$$

The covariance follows by substituting Eq. (S116) into Eq. (S105):

$$\operatorname{Cov}(X) = \Sigma + \left\langle \operatorname{diag}\left(\Lambda_g + \alpha_g\Lambda_g^2\right)_{g=1}^{G}\right\rangle_0. \tag{S119}$$

Therefore, for $g \neq h$,

$$\mathrm{Cov}(X_g, X_h) = \Sigma_{gh}, \qquad g \neq h. \tag{S120}$$

On the diagonal,

$$\mathrm{Var}(X_g) = \Sigma_{gg} + \langle \Lambda_g \rangle_0 + \alpha_g \langle \Lambda_g^2 \rangle_0. \tag{S121}$$

Using

$$\langle \Lambda_g \rangle_0 = m_g, \tag{S122}$$

and

$$\langle \Lambda_g^2 \rangle_0 = m_g^2 + \Sigma_{gg}, \tag{S123}$$

we obtain

$$\mathrm{Var}(X_g) = \Sigma_{gg} + m_g + \alpha_g(m_g^2 + \Sigma_{gg}). \tag{S124}$$

Equivalently,

$$\mathrm{Var}(X_g) = (1 + \alpha_g)\Sigma_{gg} + m_g + \alpha_i m_g^2. \tag{S125}$$

Thus the observed diagonal variance contains three contributions:

$$\mathrm{Var}(X_g) = \underbrace{\Sigma_{gg}}_{\text{biological variance}} + \underbrace{m_g}_{\text{Poisson sampling noise}} + \underbrace{\alpha_g(m_g^2 + \Sigma_{gg})}_{\text{negative-binomial overdispersion}}. \tag{S126}$$

Solving Eq. (S125) for the latent biological variance gives

$$\Sigma_{gg} = \frac{\mathrm{Var}(X_g) - m_g - \alpha_g m_g^2}{1 + \alpha_g}. \tag{S127}$$

Thus, for negative-binomial counts, subtracting only the Poisson diagonal term $\mathrm{diag}(m)$ is not sufficient. The correct diagonal correction also depends on the overdispersion $\alpha_i$. The off-diagonal covariance remains unchanged because the conditional observation noise is diagonal under conditional independence.

Finally, nonlinear transformations such as $\log(X + 1)$ generally do not preserve the linear-response relation. In general,

$$\langle \log(X_u + 1) \rangle - \langle \log(X_0 + 1) \rangle \neq \Sigma u. \tag{S128}$$

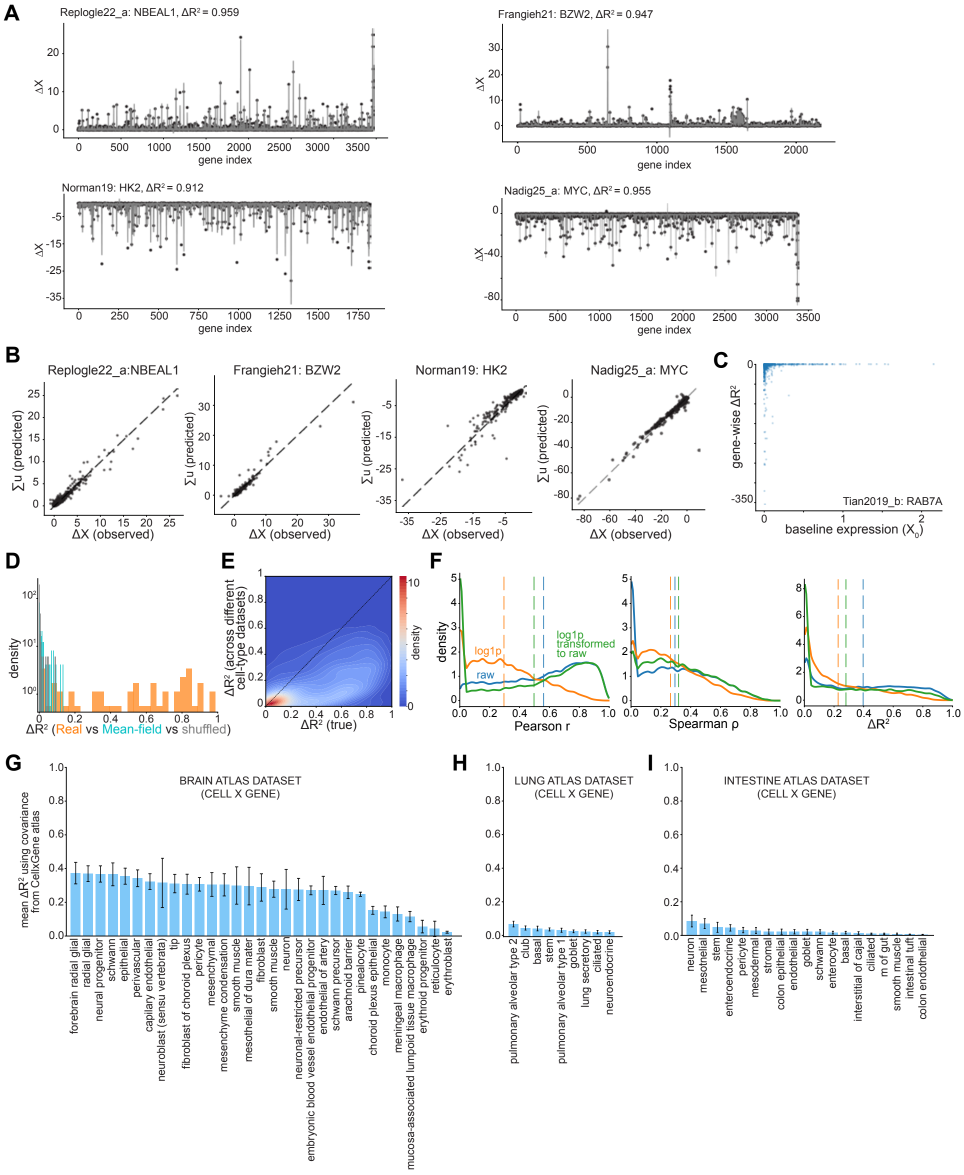


**Fig. S4**: **Perturbation response from covariance structure in single gene perturbation.** A) Forward problem predicted (dots) and experimentally measured (lines) $\Delta X$ for 4 high performing genes across datasets. B) Predicted vs observed $\Delta X$ for the 4 perturbations in A). Each point is a gene. C) Gene-wise $\Delta R^2$ as a function of baseline control expression for a single-gene perturbation. D) Perturb-fish $\Delta R^2$ histogram over perturbations. E) Same vs cross-dataset $\Delta R^2$ for Neurons. F) Metrics in raw gene expression space (blue), log1p space (orange) and fit in log1p space and transformed to raw (green) for 10 representative single-gene perturbation datasets. G)-I) Neuron $\Delta R^2$ from perurbation experiments using cell atlas covariances from brain G), lung H), and intestine I) atlases.

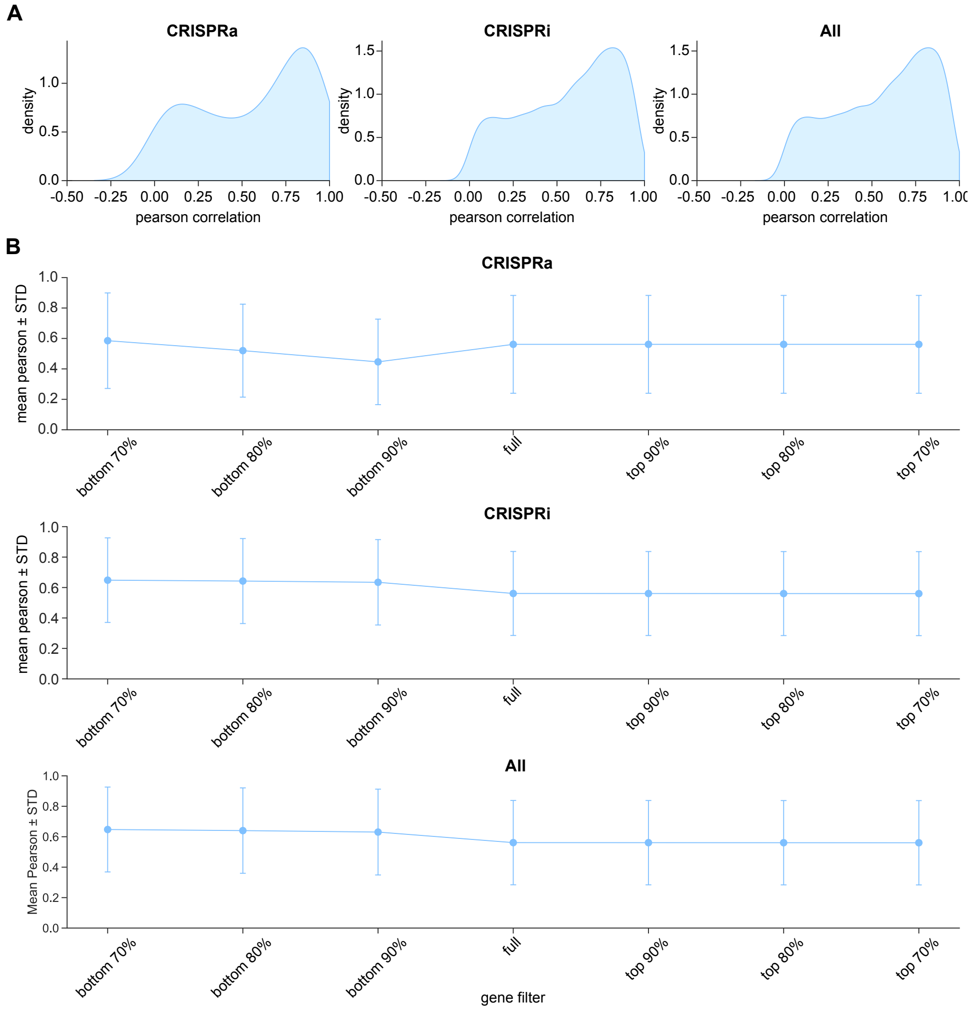


**Fig. S5**: **Sensitivity of the forward problem to highly expressed genes.** A) Distributions of Pearson correlation values over perturbations for selected CRISPRa datasets, CRISPRi datasets, and combined. B) Mean Pearson correlation across perturbations in A) where data has been pre-processed to 1) contain all genes (center), or 2) drop a certain percentage of highly-expressed (left) or lowly-expressed (right) genes.

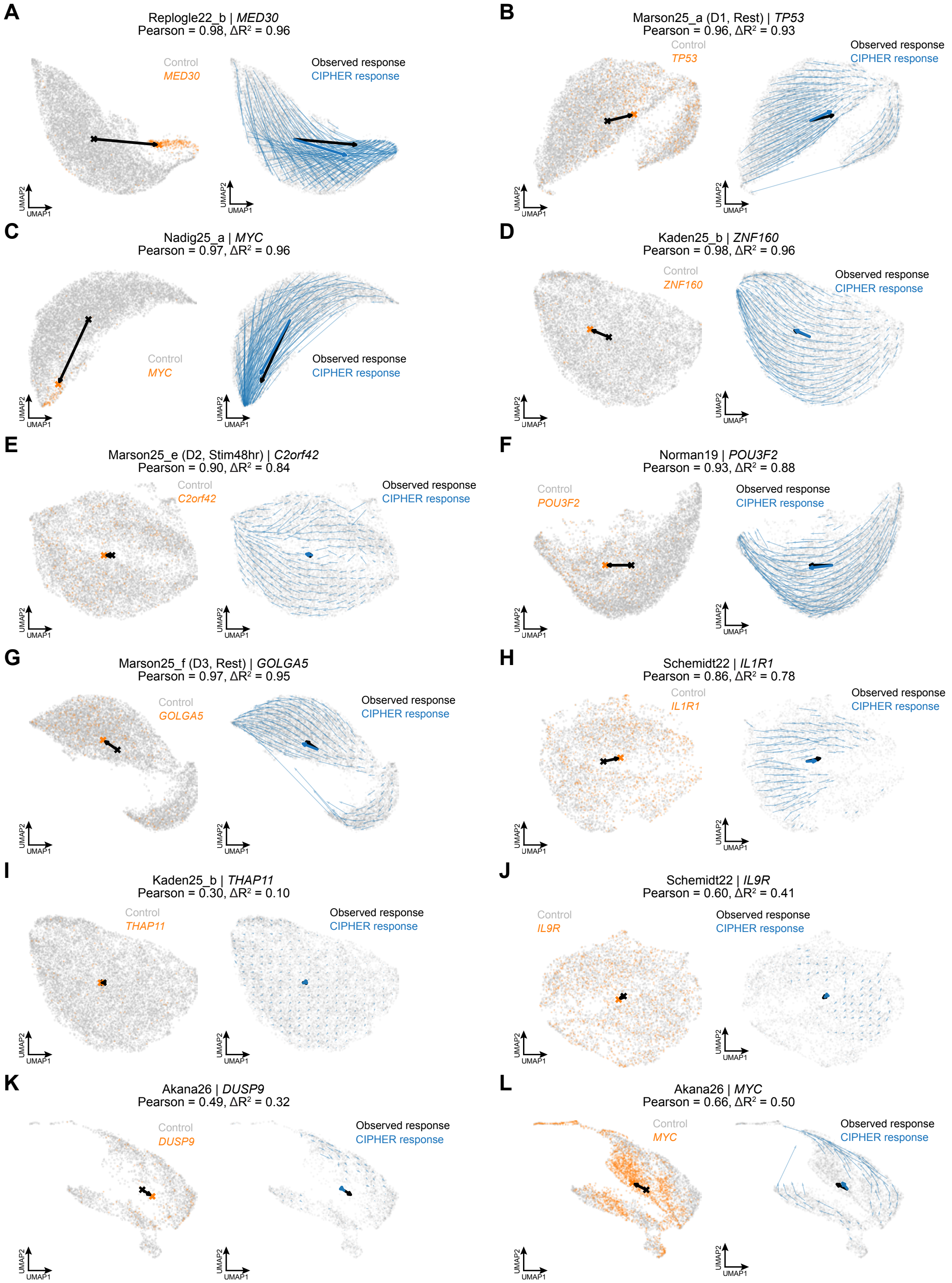


**Fig. S6**: **CIPHER UMAP** A)-L) UMAPS for different gene perturbations across datasets. Left) Control (Gray) and Perturbed (orange) gene expression. Black arrows are average displacement due to the perturbation. Right) CIPHER response flow field (blue arrows) on the control background.

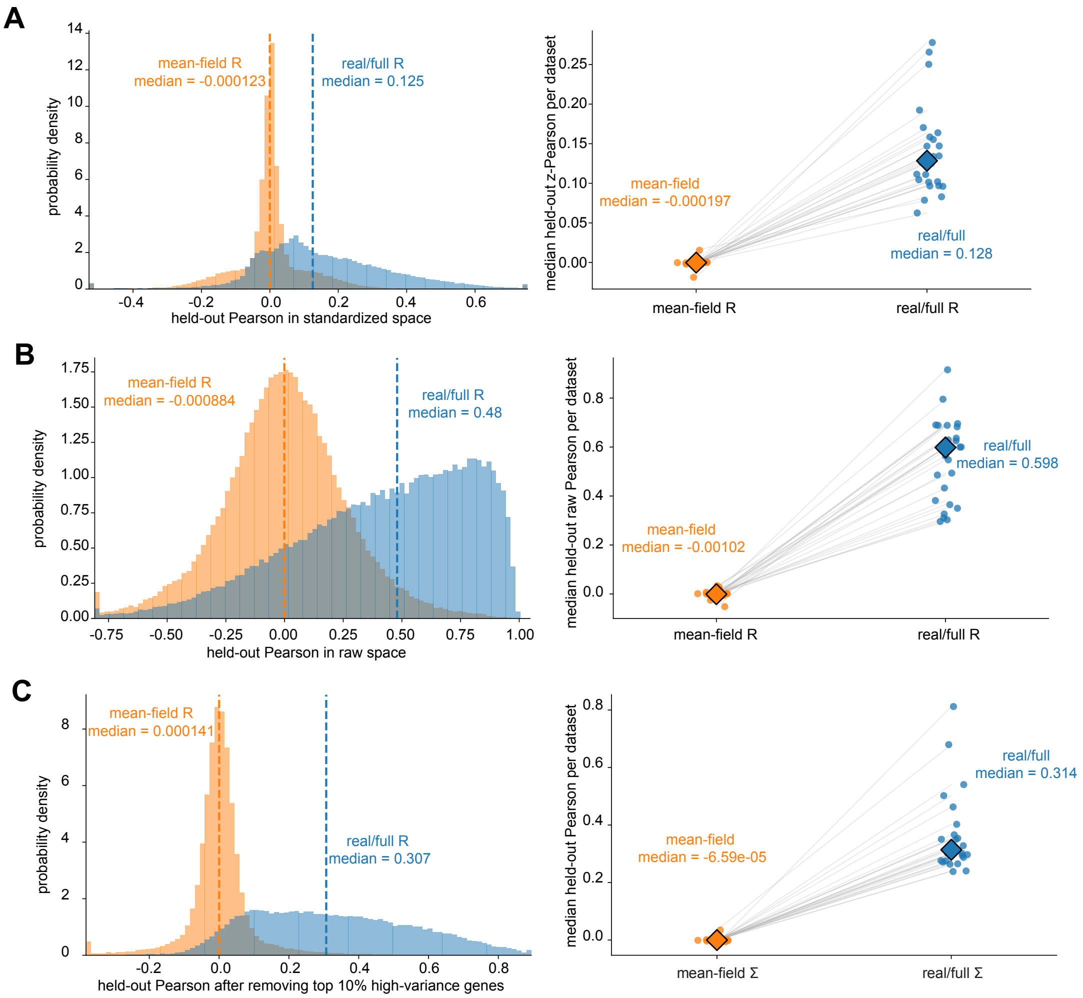


**Fig. S7**: **Additional forward-problem experiments** Left) Distributions of Pearson correlation values over perturbations and Right) dataset stratified dot plots of median scores. A) Correlation (not covariance) coordinate fit. B) Correlation coordinate space fit, transformed to raw space. C) Covariance fit after removing the top 10% of highly variable genes.

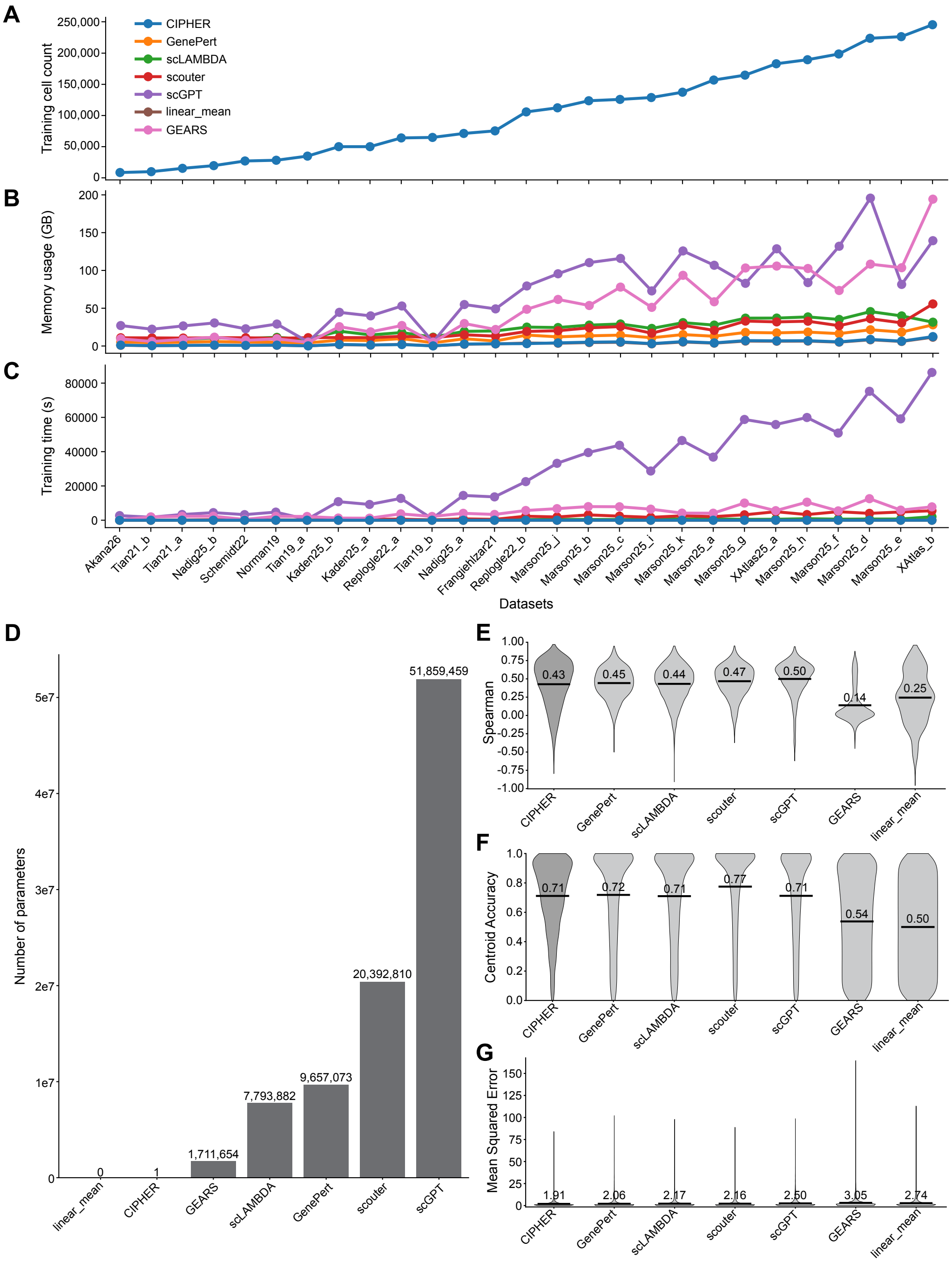


**Fig. S8**: **Computational performance and further comparative metrics of benchmarked methods.** A) The total number of training cells across datasets. B) Total memory usage and C) model training time of the benchmarked methods across datasets. D) Number of model parameters across the benchmarked methods. E) Distributions of Spearman correlation values F) centroid accuracies, and G) mean squared error values over perturbations.

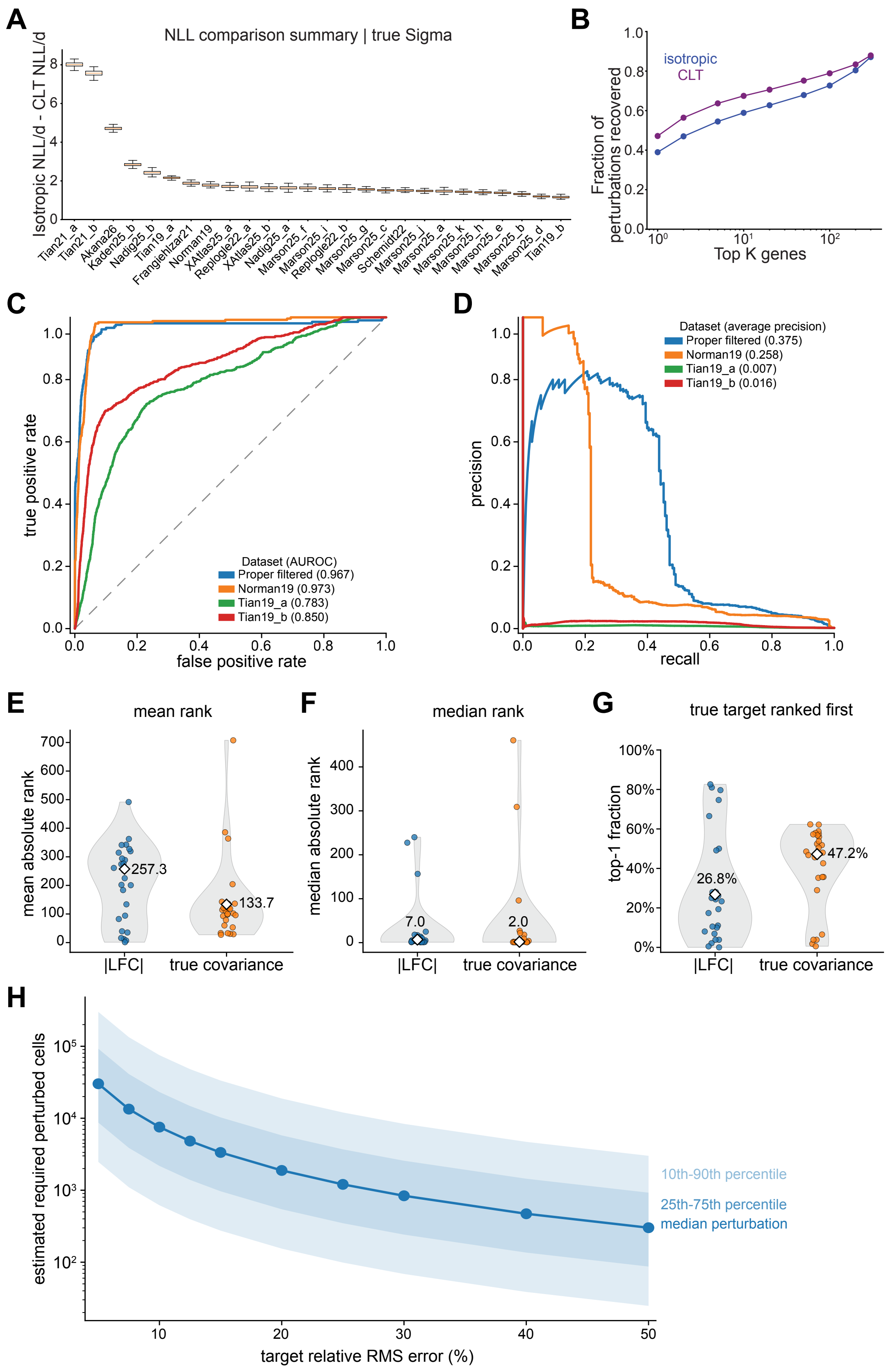

A
NLL comparison summary | true Sigma
Isotropic NLL/d - CLT NLL/d
Tian21_a
Tian21_b
Akana26
Kaden25_b
Nadig25_b
Tian19_a
FrangiehIzar21
Norman19
XAtlas25_a
Replogle22_a
XAtlas25_b
Nadig25_a
Marson25_f
Marson25_i
Replogle22_b
Marson25_g
Marson25_c
Schmidt22
Marson25_j
Marson25_a
Marson25_k
Marson25_h
Marson25_e
Marson25_b
Marson25_d
Tian19_b
B
isotropic
CLT
Fraction of perturbations recovered
Top K genes
C
true positive rate
false positive rate
Dataset (AUROC)
Proper filtered (0.967)
Norman19 (0.973)
Tian19_a (0.783)
Tian19_b (0.850)
D
precision
recall
Dataset (average precision)
Proper filtered (0.375)
Norman19 (0.258)
Tian19_a (0.007)
Tian19_b (0.016)
E
mean rank
mean absolute rank
257.3
133.7
|LFC|
true covariance
F
median rank
median absolute rank
7.0
2.0
|LFC|
true covariance
G
true target ranked first
top-1 fraction
26.8%
47.2%
|LFC|
true covariance
H
estimated required perturbed cells
target relative RMS error (%)
10th-90th percentile
25th-75th percentile
median perturbation

**Fig. S9**: **The inverse problem: Central Limit Theorem noise and double perturbation prediction. (Top) A)** Per-dataset negative log-likelihood gain of the covariance-aware CLT/full-$H$ null relative to a trace-matched isotropic null, evaluated on control-only pseudo-perturbations generated from control cells. Positive values of $\mathrm{NLL}_{\mathrm{iso}} - \mathrm{NLL}_{\mathrm{CLT}}$ indicate that finite-sampling control fluctuations are better described by the empirical covariance structure than by an isotropic approximation. The consistently positive gains across datasets show that control-cell noise is strongly anisotropic even in the absence of perturbation signal. **B)** Practical top-$K$ target recovery for the inverse problem on real perturbation data. For each perturbation, genes were ranked using either the isotropic inverse score or the covariance-aware CLT/full-$H$ inverse score, and the fraction of perturbations for which the true target appeared among the top $K$ ranked genes was computed. The covariance-aware model improves recovery across most $K$, especially at small and intermediate $K$, showing that improved null calibration leads to better practical target identification. **(Bottom)** Each double perturbation contains two known target genes, both of which were treated as positives. Genes were ranked by the full-$H_0$ posterior inverse score after covariance whitening. **C)** ROC curves for pooled target-versus-nontarget discrimination across double perturbations within each dataset. **D)** precision–recall curves for the same recovery task, emphasizing enrichment of the two true targets near the top of the ranked gene list. **E)** Mean absolute rank of the true perturbed gene. **F)** Median absolute rank of the true perturbed gene. **G)** Fraction of perturbations for which the true perturbed gene was ranked first. Violin plots show the distribution across datasets. **H)** Estimated perturbed cell numbers required to achieve different target relative RMS errors under the central limit theorem approximation and linear-response framework. The solid line shows the median across perturbations, while the dark and light shaded regions denote the 25th–75th and 10th–90th percentile ranges, respectively. Estimated cell numbers were computed from the control covariance and observed perturbation responses, in accordance with the Central Limit Theorem and linear response.

### 5 Inverse problem: predicting perturbations from correlations and response

We can also reframe the linear response problem as a Bayesian linear regression [32],

$$\Delta X = \Sigma \mathbf{u} + \xi \tag{S129}$$

where $\Delta X$ is the data, $\Sigma u$ is the model with $\Sigma$ fixed and the elements of $\xi$ are random Normally distributed error terms, with mean zero and variance $\sigma^2$. Here $\xi$ can be thought of as modeling co-variates unaccounted for in linear response. Using Bayes' rule, it can be shown that the posterior probability of observing $\mathbf{u}$ given the data is, up to a constant,

$$\ln p(\mathbf{u}|\Delta X) = -\frac{||\Delta X - \Sigma \mathbf{u}||^2}{2\sigma^2} + \ln p(\mathbf{u}) \tag{S130}$$

where $p(\mathbf{u})$ is a prior on the perturbation.

The likelihood term above assumed i.i.d. Gaussian noise, $\xi \sim \mathcal{N}(0, \sigma^2 I)$, so that the quadratic misfit is Euclidean. More generally, the linear-response residuals are correlated across genes, and it is natural to model

$$\xi \sim \mathcal{N}(0, H), \tag{S131}$$

where $H$ is the noise covariance of $\Delta X$. In this case the Gaussian log-likelihood becomes the generalized least-squares form

$$\ln p(\Delta X \mid \mathbf{u}) \;=\; -\frac{1}{2}\,(\Delta X - \Sigma \mathbf{u})^\top H^{-1} (\Delta X - \Sigma \mathbf{u}) - \frac{1}{2} \ln |H| + \text{const.}, \tag{S132}$$

and therefore (up to an additive constant independent of $\mathbf{u}$),

$$\ln p(\mathbf{u}|\Delta X) \;=\; -\frac{1}{2}\,(\Delta X - \Sigma \mathbf{u})^\top H^{-1} (\Delta X - \Sigma \mathbf{u}) + \ln p(\mathbf{u}). \tag{S133}$$

Two useful "flavors" of $H$ appear repeatedly in practice. The simplest choice is isotropic noise,

$$H \propto I, \tag{S134}$$

which reduces to the scalar-variance model above and treats all gene directions as equally reliable.

A more mechanistic choice follows from how $\Delta X$ is constructed. If $\Delta X$ is a difference of sample means between two conditions,

$$\Delta X \;=\; \bar{X}_1 - \bar{X}_0, \tag{S135}$$

and if cells are (approximately) independent within each condition with within-condition covariances $\mathrm{Cov}(X_0) = \Sigma_0$ and $\mathrm{Cov}(X_1) = \Sigma_1$, then

$$H \;=\; \mathrm{Cov}(\Delta X) \;=\; \frac{\Sigma_0}{n_0} + \frac{\Sigma_1}{n_1}, \tag{S136}$$

where $n_0, n_1$ are the numbers of cells (or effective sample sizes) used to form each mean. Eq. (S136) captures heteroscedasticity and gene–gene correlated noise induced by sampling.

To make Eq. (S136) computationally convenient, we can approximate $H$ in the eigenbasis of $\Sigma_0$. Let $\Sigma_0 = V\Lambda V^\top$ with diagonal $\Lambda = \mathrm{diag}(\lambda_1, \ldots, \lambda_G)$. Then

$$H \;=\; V\Big(\frac{\Lambda}{n_0} + \frac{V^\top \Sigma_1 V}{n_1}\Big)V^\top. \tag{S137}$$

If $\Sigma_1$ is close to $\Sigma_0$ (e.g. perturbations are modest relative to baseline variability), then $V^\top \Sigma_1 V$ is approximately diagonal, and a simple diagonal approximation is

$$H \;\approx\; V\,\mathrm{diag}\Big(\frac{\lambda_i}{n_0} + \frac{\tilde{\lambda}_i}{n_1}\Big)_{i=1}^{G} V^\top, \tag{S138}$$

where $\tilde{\lambda}_i \equiv (V^\top \Sigma_1 V)_{ii}$ are the variances of condition 1 projected onto the condition 0 principal directions. A common further simplification is $\Sigma_1 \approx \Sigma_0$, yielding

$$H \;\approx\; \Big(\frac{1}{n_0} + \frac{1}{n_1}\Big)\Sigma_0, \tag{S139}$$

so that the likelihood effectively measures misfit in the $\Sigma_0^{-1}$-whitened metric. In implementations, inversion of $H$ (or $\Sigma_0$) is typically stabilized by a small ridge term, $H \leftarrow H + \epsilon I$, to control poorly conditioned soft modes.

#### Covariance-aware null calibration improves inverse target recovery

To evaluate calibration of the null noise model used in inverse target recovery, we first performed a control-only likelihood test in which no perturbation signal was present. In this analysis, the applied perturbation was set to zero and the pseudo-perturbation mean was generated by drawing cells from the control population. Thus, the observed mean shift reflects only finite-sampling fluctuations among control cells, rather than a biological response to perturbation. This provides a direct test of whether the null model correctly describes the fluctuations expected when two sample means are drawn from the same underlying control distribution.

We compared two null models for these control-derived mean shifts. The first was a trace-matched isotropic null, which preserves the total variance but assumes that fluctuations are equally likely in all directions of expression space. The second was a covariance-aware CLT/full-$H$ null, which preserves the anisotropic covariance structure estimated from control cells. For a control-derived mean shift $\Delta X$, the relevant quantity is the negative log likelihood under the null model,

$$\mathrm{NLL}(\Delta X) = \frac{1}{2}\Delta X^T H^{-1} \Delta X + \frac{1}{2}\log\det H + \text{constant}, \tag{S140}$$

where $H$ is either the isotropic covariance or the CLT/full-$H$ covariance of the sample mean. We summarized the advantage of the covariance-aware null by the per-dataset gain

$$\Delta\text{NLL} = \text{NLL}_{\text{iso}} - \text{NLL}_{\text{CLT}} \tag{S141}$$

where $d$ is the number of genes used in the inverse problem. Positive values indicate that control fluctuations are more compatible with the covariance-aware null than with the isotropic approximation.

As shown in Fig. S9A, the likelihood gain is consistently positive across datasets. This shows that even in the no-signal setting, control-cell fluctuations are strongly anisotropic and are better described by the empirical covariance structure than by a trace-matched isotropic model. In other words, the covariance-aware null is better calibrated to the actual geometry of control fluctuations.

We then asked whether this improved null calibration translates into better inverse recovery on real perturbation data. For each perturbation, genes were ranked using either the isotropic inverse score or the CLT/full-$H$ inverse score, and we measured the fraction of perturbations for which the known target gene was recovered within the top $K$ ranked genes. The practical top-$K$ recovery curves in Fig. S9B show that the CLT/full-$H$ model improves recovery across most values of $K$, with the clearest gains at small and intermediate $K$, where target identification is most useful experimentally. Thus, better calibration of the control-only null model leads directly to better target recovery in the inverse problem.

##### 5.0.1 Power analysis for perturbation sample size

To estimate the number of perturbed cells required to accurately measure a perturbation response, we performed an analytical power analysis using the precomputed control covariance matrix and observed perturbation mean shifts. Only the precomputed covariance matrices and perturbation summary statistics were required; no raw single-cell data were reloaded and no covariance matrices were recomputed.

We assumed that the number of control cells greatly exceeded the number of perturbed cells, allowing uncertainty in the control mean to be neglected. Furthermore, we assumed that the covariance of perturbed cells was well approximated by that of control cells,

$$\Sigma_{\text{pert}} \approx \Sigma, \tag{S142}$$

where $\Sigma$ denotes the control-cell covariance matrix. Under these assumptions, the estimated perturbation response is

$$\widehat{\Delta x} = \bar{x}_{\text{pert}} - \bar{x}_{\text{control}}, \tag{S143}$$

with expectation equal to the true perturbation response.

Because averaging over a finite number $m$ of perturbed cells introduces sampling error, the expected squared norm of the observed response satisfies

$$\mathbb{E}\left[\|\widehat{\Delta x}\|^2\right] = \|\Delta x\|^2 + \frac{\text{tr}(\Sigma)}{m}, \tag{S144}$$

where $\text{tr}(\Sigma)$ is the total transcriptomic variance. This motivates the unbiased estimator of the true signal magnitude,

$$\widehat{S} = \|\widehat{\Delta x}\|^2 - \frac{\text{tr}(\Sigma)}{m}, \tag{S145}$$

which estimates $\|\Delta x\|^2$. Perturbations for which $\widehat{S} \leq 0$ were considered insufficiently resolved above sampling noise and were excluded from subsequent sample-size estimation.

The relative root-mean-square (RMS) error of the estimated response scales as

$$\epsilon = \sqrt{\frac{\text{tr}(\Sigma)}{m\,\|\Delta x\|^2}}, \tag{S146}$$

which can be inverted to estimate the number of perturbed cells required to achieve a desired target relative error,

$$m_{\text{required}}(\epsilon) = \frac{\operatorname{tr}(\Sigma)}{\epsilon^2 \widehat{S}}. \tag{S147}$$

Required sample sizes were computed independently for each perturbation over target relative RMS errors ranging from 5% to 50%.

Finally, required cell numbers were summarized across perturbations using the median together with the 10th–90th and 25th–75th percentile ranges, and analogous summaries were computed separately for each dataset to quantify the distribution of sampling requirements across experimental systems.

##### 5.0.2 Analytical approach on full perturbation space

A Gaussian prior with variance $\tau^2$,

$$\mathbf{u} \sim \mathcal{N}(0, \tau^2 I), \tag{S148}$$

enforces shrinkage while preserving analytic tractability.

***General correlated noise.***

Instead of assuming i.i.d. noise, we allow a general Gaussian noise covariance

$$\xi \sim \mathcal{N}(0, H), \tag{S149}$$

so that the likelihood is

$$p(\Delta X \mid \mathbf{u}) \propto \exp\Big( -\tfrac{1}{2}(\Delta X - \Sigma\mathbf{u})^\top H^{-1}(\Delta X - \Sigma\mathbf{u})\Big). \tag{S150}$$

Combining this likelihood with the isotropic Gaussian prior yields a Gaussian posterior

$$p(\mathbf{u} \mid \Delta X) = \mathcal{N}(\mu_u, \operatorname{Cov}[\mathbf{u}]), \tag{S151}$$

with

$$\mu_u = \Big(\Sigma^\top H^{-1}\Sigma + \tfrac{1}{\tau^2} I\Big)^{-1} \Sigma^\top H^{-1} \Delta X, \tag{S152}$$

$$\operatorname{Cov}[\mathbf{u}] = \Big(\Sigma^\top H^{-1}\Sigma + \tfrac{1}{\tau^2} I\Big)^{-1}. \tag{S153}$$

The posterior mean $\mu_u$ is also the maximum-a-posteriori (MAP) estimate, which we denote $\hat{\mathbf{u}}$.

***Solving for $\hat{\mathbf{u}}$ in an eigenbasis.***

Directly forming $H^{-1}$ or inverting the $G \times G$ matrix in Eqs. (S152)–(S153) is expensive at genome scale. A convenient approach is to work in a basis that (approximately) diagonalizes the noise metric. Let

$$H = V\Lambda_H V^\top, \qquad \Lambda_H = \operatorname{diag}(h_1, \ldots, h_G), \tag{S154}$$

so that $H^{-1/2} = V\Lambda_H^{-1/2}V^\top$. Define whitened quantities

$$\tilde{y} \equiv H^{-1/2}\Delta X, \qquad \tilde{\Sigma} \equiv H^{-1/2}\Sigma. \tag{S155}$$

Then the MAP estimator becomes standard ridge regression in the whitened space,

$$\hat{\mathbf{u}} = \Big(\tilde{\Sigma}^\top\tilde{\Sigma} + \tfrac{1}{\tau^2} I\Big)^{-1} \tilde{\Sigma}^\top \tilde{y}, \tag{S156}$$

and the posterior covariance is

$$\mathrm{Cov}[\mathbf{u}] = \left(\tilde{\Sigma}^\top \tilde{\Sigma} + \tfrac{1}{\tau^2} I\right)^{-1}. \tag{S157}$$

Operationally, applying $H^{-1/2}$ in the eigenbasis amounts to scaling the coordinates of any vector $z$ by $h_i^{-1/2}$ in the $V$ basis: $H^{-1/2}z = V\Lambda_H^{-1/2}(V^\top z)$. When $H$ is taken from Eq. (S136) and approximated in the $\Sigma_0$ eigenbasis (so that $V$ are the $\Sigma_0$ eigenvectors), the same procedure yields a fast approximate whitening that captures correlated sampling noise.

***Per-perturbation summaries and conservative scores.***

For each perturbation $p$, we summarize inferred effects using the posterior mean $\mu_u^{(p)}$ and posterior standard deviation $\sigma_u^{(p)} \equiv \sqrt{\mathrm{diag}(\mathrm{Cov}[\mathbf{u}])}$. To produce a scalar score per gene that reflects both magnitude and uncertainty, we use

$$s_j^{(p)} = \max\left(\left|\mu_{u,j}^{(p)} + \sigma_{u,j}^{(p)}\right|,\ \left|\mu_{u,j}^{(p)} - \sigma_{u,j}^{(p)}\right|\right), \tag{S158}$$

which prioritizes genes whose inferred effects are large and robust to posterior uncertainty. For evaluation, each perturbation is associated with a binary ground-truth vector $y^{(p)} \in \{0,1\}^G$ indicating which genes were directly targeted; each perturbation therefore yields paired score–label sets $(s^{(p)}, y^{(p)})$ for ranking analyses and ROC evaluation.

***Empirical Bayes hyperparameter selection.***

The posterior depends on the prior scale $\tau^2$ and on the noise model through $H$ (e.g. via sample-size scaling and/or a ridge stabilization $H \leftarrow H + \epsilon I$). Rather than fixing $\tau^2$ a priori (or tuning it for predictive performance), we select it by empirical Bayes: we minimize the negative log marginal likelihood (NLL) of the data under the linear–Gaussian model.

With $\mathbf{u} \sim \mathcal{N}(0, \tau^2 I)$ and $\xi \sim \mathcal{N}(0, H)$, integrating out $\mathbf{u}$ gives the marginal

$$\Delta X \sim \mathcal{N}\big(0,\ C(\tau^2)\big), \qquad C(\tau^2) \equiv H + \tau^2 \Sigma\Sigma^\top. \tag{S159}$$

Up to an additive constant, the corresponding NLL is

$$\mathcal{L}(\tau^2) \equiv -\ln p(\Delta X \mid \tau^2) = \frac{1}{2}\left[\Delta X^\top C(\tau^2)^{-1}\Delta X + \ln\left|C(\tau^2)\right|\right]. \tag{S160}$$

We then choose

$$\hat{\tau}^2 \ = \ \arg\min_{\tau^2 \in [\tau^2_{\min}, \tau^2_{\max}]} \mathcal{L}(\tau^2), \tag{S161}$$

where the bounded interval $[\tau^2_{\min}, \tau^2_{\max}]$ is imposed to prevent degenerate solutions in which the prior either collapses ($\tau^2 \to 0$) or becomes effectively uninformative ($\tau^2 \to \infty$) due to finite-sample noise and imperfect model specification. In practice, we minimize Eq. (S160) with a one-dimensional bracketed search (e.g. over $\log \tau^2$) on the interval Eq. (S161), which is robust and fast.

Having selected $\hat{\tau}^2$, we compute the posterior mean $\hat{\mathbf{u}}$ and posterior covariance using Eqs. (S152)–(S153), with the same $H$ (including any ridge stabilization) used in the marginal likelihood.

#### Inverse recovery of double-perturbation targets

We next evaluated whether the full-$H_0$ posterior inverse model could recover the known target genes in double-perturbation experiments. Each perturbation in this analysis contains two target genes, so both genes were treated as positives in the recovery task. For perturbation $j$, the observed response was computed as the mean expression shift relative to controls,

$$\Delta X_j = \langle X \rangle_j - \langle X \rangle_0\,, \tag{S162}$$

and candidate genes were ranked by the full-$H_0$ posterior inverse score after covariance whitening. Performance was then summarized by pooled ROC and precision–recall curves across double perturbations within each dataset. Known perturbed genes were force-included after expression filtering, so failures of recovery reflect the inverse score itself rather than accidental loss of the true targets during preprocessing.

The analysis used the full-$H_0$ posterior inverse model in covariance-normalized expression space. Specifically, the control covariance was used both as the whitening covariance and as the linear-response covariance, so inference was performed on the covariance-standardized response.

For the double-perturbation analysis, we used substantially weaker shrinkage than in the single-gene inverse experiments. The prior variance $\tau^2$ was fit by empirical Bayes over $\log \tau^2 \in [6, 8]$, corresponding to $\tau^2 \approx e^6$ to $e^8$, or roughly $4 \times 10^2$ to $3 \times 10^3$. For scoring, we selected the largest $\tau^2$ within the marginal-likelihood plateau. This choice places the posterior in a weakly regularized regime where the likelihood, rather than the prior, more strongly determines the inferred perturbation vector. Such a choice is appropriate for multigene perturbations, because the true perturbation is not expected to be concentrated on a single coordinate. Instead, two target genes are directly perturbed, and the measured transcriptional response may distribute posterior mass across both targets as well as correlated downstream directions. A smaller $\tau^2$ would impose stronger shrinkage toward zero and could suppress one or both true targets.

The ROC curves show that the full-$H_0$ posterior score separates true target genes from non-target genes across the tested datasets. Performance is strongest for the Proper filtered and NormanWeissman2019 datasets, and remains above chance for the TianKampmann day-7 neuron and iPSC datasets. The precision–recall curves provide a stricter measure of practical recovery, since each perturbation has only two positives among many candidate genes. Under this metric, the Proper filtered and NormanWeissman2019 datasets show the clearest enrichment of true targets near the top of the ranked list, whereas the Tian datasets show weaker average precision despite non-random ROC performance. Thus, in some datasets the target genes are ranked better than background genes overall, but are not always concentrated at the very top of the posterior ranking. This is particularly true for soft-mode dominated datasets that that exhibit a great deal of covariance column degeneracy.

Together, these results show that the full-$H_0$ posterior inverse score contains information about both components of double perturbations. The large-$\tau^2$ regime is an important part of this analysis: it relaxes the strong shrinkage useful for conservative single-gene attribution and instead allows the posterior to represent multigene perturbation structure. The double-perturbation recovery task therefore probes a more permissive inverse regime, asking whether covariance-normalized linear response can identify multiple directly perturbed genes from a combined transcriptional response.

#### 5.1 Drug response analysis via likelihood-based matching

We next apply the linear-Gaussian response framework to drug perturbation data, treating each drug $d$ as inducing a characteristic transcriptional response vector $\mathbf{u}_d$. For each drug, we split the data into a training set and a held-out test set at the level of single cells (or pseudobulk replicates). Using only the training subset for drug $d$, we estimate its response vector $\hat{\mathbf{u}}_d$ by empirical Bayes linear response, exactly as described above, with the same noise model $H_d$ (including sample-size scaling and any ridge stabilization).

Concretely, let $\Delta X_d^{\text{train}}$ denote the mean expression shift between drug-treated and control cells in the training split. From $\Delta X_d^{\text{train}}$ we infer

$$\hat{\mathbf{u}}_d = \arg\max_{\mathbf{u}} \; p(\mathbf{u} \mid \Delta X_d^{\text{train}}), \tag{S163}$$

together with an associated noise covariance $H_d$ for that drug. This yields a library of drug-specific linear response models $\{(\hat{\mathbf{u}}_d, H_d)\}_{d=1}^{D}$, learned without access to the test data.

To classify held-out samples, we compute the observed transcriptional shift $\Delta X^{\text{test}}$ from the test cells and evaluate its likelihood under each drug-specific model. Conditioned on drug $d$, the predictive distribution is

$$\Delta X^{\text{test}} \;\sim\; \mathcal{N}\big(\Sigma \hat{\mathbf{u}}_d,\; H_d\big), \tag{S164}$$

so that the (unnormalized) log-likelihood is

$$\ell_d(\Delta X^{\text{test}}) = -\frac{1}{2}\big(\Delta X^{\text{test}} - \Sigma \hat{\mathbf{u}}_d\big)^\top H_d^{-1}\big(\Delta X^{\text{test}} - \Sigma \hat{\mathbf{u}}_d\big) - \frac{1}{2}\ln |H_d|. \tag{S165}$$

We rank drugs by $\ell_d$, and assign the test sample to the drug

$$\hat{d} = \arg\max_{d} \; \ell_d(\Delta X^{\text{test}}). \tag{S166}$$

This likelihood-based ranking has a direct mechanistic interpretation: each drug $d$ defines a predicted mean response $\Sigma\hat{\mathbf{u}}_d$ and an uncertainty ellipsoid set by $H_d$, and classification selects the drug whose linear-response model best explains the observed test shift in the appropriate noise-weighted metric. Because $\hat{\mathbf{u}}_d$ is learned only from training cells, this procedure evaluates how well drug-specific transcriptional programs generalize to unseen data, rather than memorizing individual cells.

Beyond top-1 classification accuracy, the full likelihood ranking allows finer-grained analyses, including margins between the best and second-best drugs, confusion patterns between transcriptionally similar compounds, and robustness as a function of test-set size. In all cases, inference and evaluation are performed entirely within the same probabilistic framework, with no additional tuning beyond the empirical Bayes selection of $\tau^2$ and the specification of $H_d$.

### 6 Pharmaceutical perturbations

#### 6.1 Drug perturbations are inferred reproducibly across Tahoe biological replicates

To test whether inverse-response scoring can identify the correct drug from an observed transcriptional response, we applied CIPHER to the Tahoe drug-response dataset in a replicate-held-out design. The key feature of this benchmark is that fitting and testing were performed on independent biological replicates. In one direction, replicate 1 was used to fit drug-response signatures and replicate 2 was used for testing; in the reverse direction, replicate 2 was used for fitting and replicate 1 was used for testing. All analyses were restricted to the top 2,000 highly variable genes. Therefore, successful recovery requires that the inferred drug-response structure generalize across biological replicates rather than simply fitting replicate-specific noise.

For each drug $d$, we computed a training response vector

$$\Delta X_d^{\text{train}} = \langle X \rangle_{d,\text{train}} - \langle X \rangle_{0,\text{train}}\,, \tag{S167}$$

where $\langle X \rangle_{d,\text{train}}$ is the mean expression of cells treated with drug $d$ in the training replicate and $\langle X \rangle_{0,\text{train}}$ is the corresponding control mean. Using the training replicate covariance $\Sigma_{\text{train}}$, we inferred a drug-specific perturbation vector $u_d$ through the linear-response model

$$\Delta X_d^{\text{train}} \approx \Sigma_{\text{train}} u_d. \tag{S168}$$

For a held-out test response,

$$\Delta X_q^{\text{test}} = \langle X \rangle_{q,\text{test}} - \langle X \rangle_{0,\text{test}}\,, \tag{S169}$$

we scored each candidate drug $d$ by the Gaussian linear-response likelihood

$$\log p\left(\Delta X_q^{\text{test}} \mid u_d, \Sigma_{\text{test}}, H_q\right) = -\frac{1}{2}\left(\Delta X_q^{\text{test}} - \Sigma_{\text{test}} u_d\right)^T H_q^{-1} \left(\Delta X_q^{\text{test}} - \Sigma_{\text{test}} u_d\right) + \text{constant}, \tag{S170}$$

analogous to the inverse likelihood in Eq.(S165). Candidate drugs were ranked by this score, and the known applied drug $q$ was treated as the true positive.

We compared the covariance-aware model, LR-TRUE, to a mean-field linear-response baseline, LR-MF, as well as baselines based on log-fold change and classifier-derived scores. The assignment heatmaps summarize the confusion matrix where diagonal enrichment indicates correct recovery of the applied drug. The top-$k$ curves report the fraction of held-out drugs for which the true drug appears among the top $k$ candidates, which is the most relevant metric for experimental prioritization. ROC curves test whether true drug identities tend to score above candidate alternatives overall, while precision-recall curves emphasize enrichment of the true drug among the highest-scoring candidates. Across both

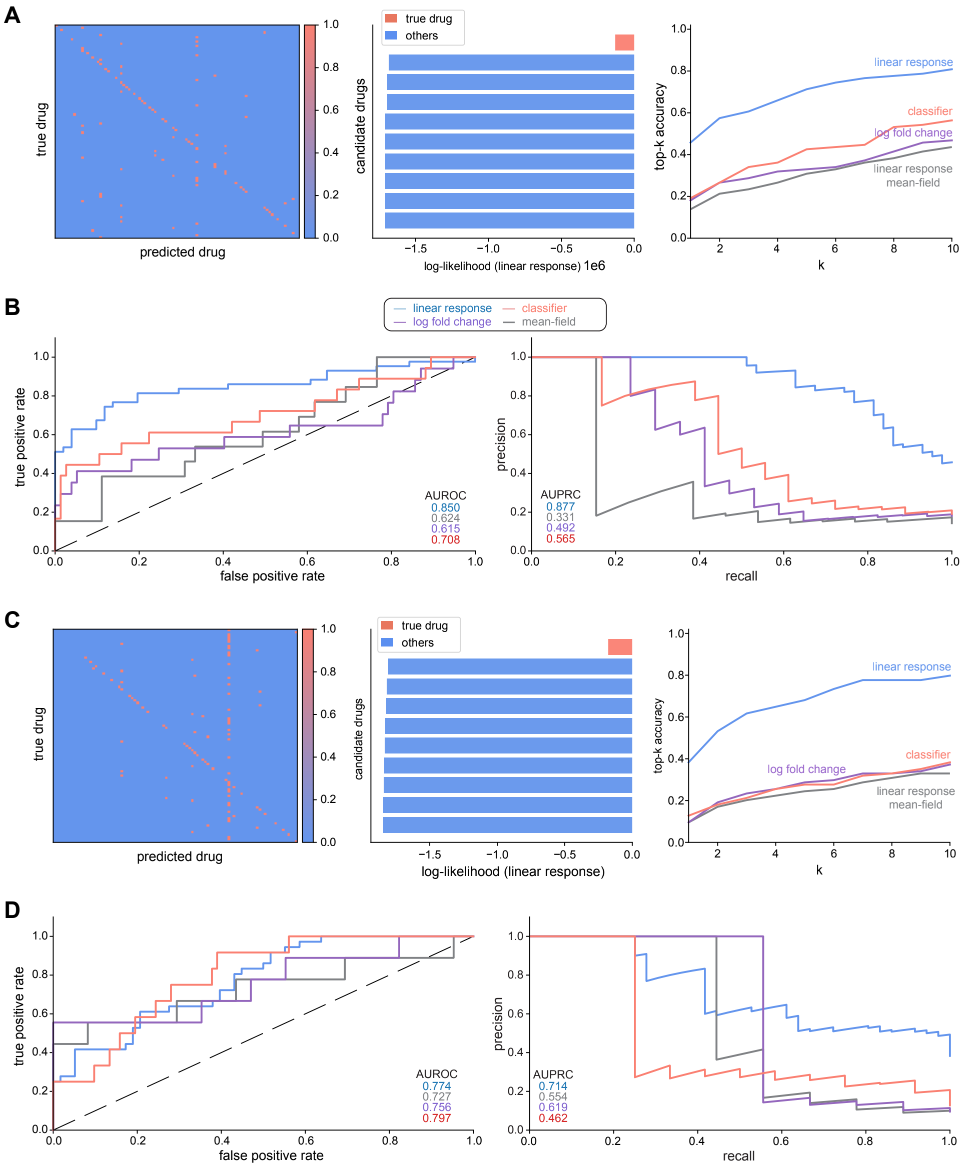


**Fig. S10**: **Tahoe drug-inference benchmark across independent biological replicates.** (A,C) Results for the two train/test replicate splits. Left: heatmaps of true versus predicted drug assignments. Middle: example ranked candidate-drug log-likelihoods with the true drug highlighted. Right: top-$k$ accuracy curves for the linear-response, mean-field, log fold change, and classifier methods. (B,D) Pooled receiver operating characteristic (ROC; left) and precision–recall (PR; right) curves for the corresponding replicate split.

replicate directions, LR-TRUE shows stronger early retrieval and precision-recall performance than the mean-field LR-MF baseline, supporting the conclusion that empirical gene-gene covariance improves drug inference beyond what is obtained from gene-wise variance alone.

#### 6.2 Conservation of inferred perturbation programs across sci-Plex drug classes

To determine whether the inferred dense intervention vectors ($u^*$) captured biologically meaningful mechanisms of action, we asked whether drugs with similar annotated targets exhibited similar inferred intervention programs. For each drug, we used the inferred $u^*$ vector obtained from the inverse CIPHER

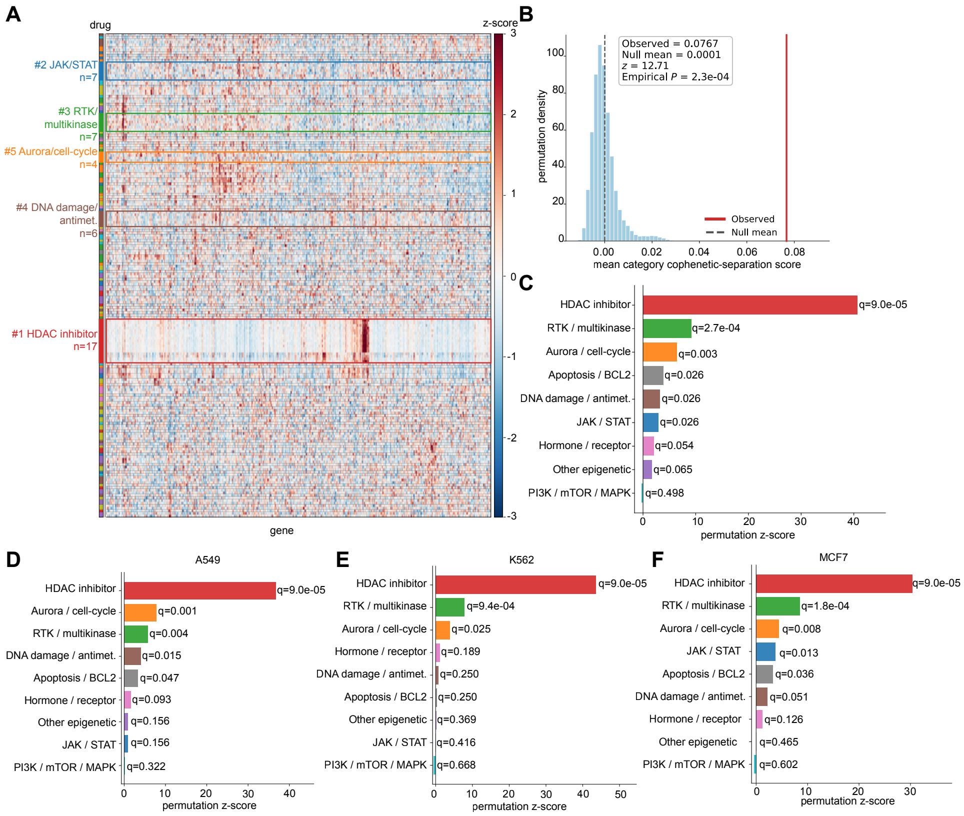


**Fig. S11**: **Conservation of inferred intervention programs across SciPlex drug classes.** (A) Hierarchically clustered heatmap of inferred intervention vectors ($u^*$) averaged across A549, K562, and MCF7 cells after RMS normalization and row-wise $z$-scoring. Rows correspond to drugs, columns to genes, and colored annotations indicate drug mechanism-of-action classes. (B) Permutation null distribution of the mean cophenetic clustering score across drug classes, with the observed value indicated. (C) Permutation $z$-scores for individual drug classes in the aggregated analysis, with Benjamini–Hochberg adjusted $q$-values shown. (D-F) Class-specific permutation $z$-scores computed independently for the A549 (D), K562 (E), and MCF7 (F) cell lines, with corresponding Benjamini–Hochberg adjusted $q$-values.

model. In the aggregate analysis, a single representative $u^*$ was obtained by averaging the inferred vectors across the three SciPlex cell lines (A549, K562, and MCF7). We additionally repeated the entire analysis independently within each cell line to assess the reproducibility of these patterns.

For visualization, each drug was represented by its genome-wide $u^*$ vector after RMS normalization followed by row-wise $z$-scoring. Drugs and genes were independently clustered using hierarchical clustering with correlation distance and average linkage. The resulting clustered drug-by-gene heatmap is shown in Fig. S11A, with drugs annotated according to broad mechanism-of-action classes.

To quantify whether drugs from the same class clustered more closely than expected by chance, we computed the cophenetic distance matrix of the drug dendrogram. For each drug class $c$, we defined a clustering score as

$$S_c = \langle d_{\text{between}} \rangle - \langle d_{\text{within}} \rangle, \tag{S171}$$

where $\langle d_{\text{within}} \rangle$ is the mean cophenetic distance between pairs of drugs within class $c$, while $\langle d_{\text{between}} \rangle$ is the mean distance between drugs in class $c$ and all drugs outside the class. Positive values therefore indicate that drugs within a class cluster more tightly than expected relative to the remainder of the dataset. A global clustering statistic was obtained by averaging $S_c$ across all tested drug classes.

Statistical significance was assessed using 100,000 permutations in which drug-class labels were randomly reassigned while preserving the dendrogram and the number of drugs assigned to each class.

Empirical $p-$values were calculated from the resulting null distributions, converted to $z$-scores, and corrected for multiple testing using the Benjamini–Hochberg procedure.

The aggregate analysis revealed strong organization of inferred intervention programs by drug mechanism (Fig. S11A). Hierarchical clustering produced extended contiguous blocks enriched for individual drug classes, indicating that pharmacologically related compounds exhibit similar inferred dense intervention vectors despite acting on different molecular targets. The global clustering score was highly significant relative to the permutation null (Fig. S11B), demonstrating that the observed organization substantially exceeded that expected from random assignment of drug classes.

Class-specific permutation tests further showed that HDAC inhibitors formed the most coherent group, with the largest enrichment above the null distribution (Fig. S11C). RTK/multikinase inhibitors and Aurora/cell-cycle inhibitors also exhibited significant clustering, while several additional classes, including DNA-damage agents, apoptosis/BCL2 inhibitors, and JAK/STAT inhibitors, showed more modest but significant enrichment. These results indicate that CIPHER infers intervention programs that capture shared regulatory effects among drugs with related mechanisms of action.

Repeating the analysis independently within A549, K562, and MCF7 cells produced highly consistent results (Fig. S11D-F). HDAC inhibitors remained the most strongly clustered class in all three cell lines, while RTK/multikinase inhibitors also showed reproducible enrichment. Aurora/cell-cycle inhibitors were significant in each individual cell line, although with reduced effect sizes compared to the aggregate analysis. Other drug classes exhibited greater variability across cell types, consistent with the presence of cell-type-specific transcriptional responses. Overall, these results demonstrate that the inferred intervention vectors recover biologically meaningful, mechanism-specific programs that are reproducible across diverse cellular backgrounds while remaining sensitive to context-dependent regulatory differences.

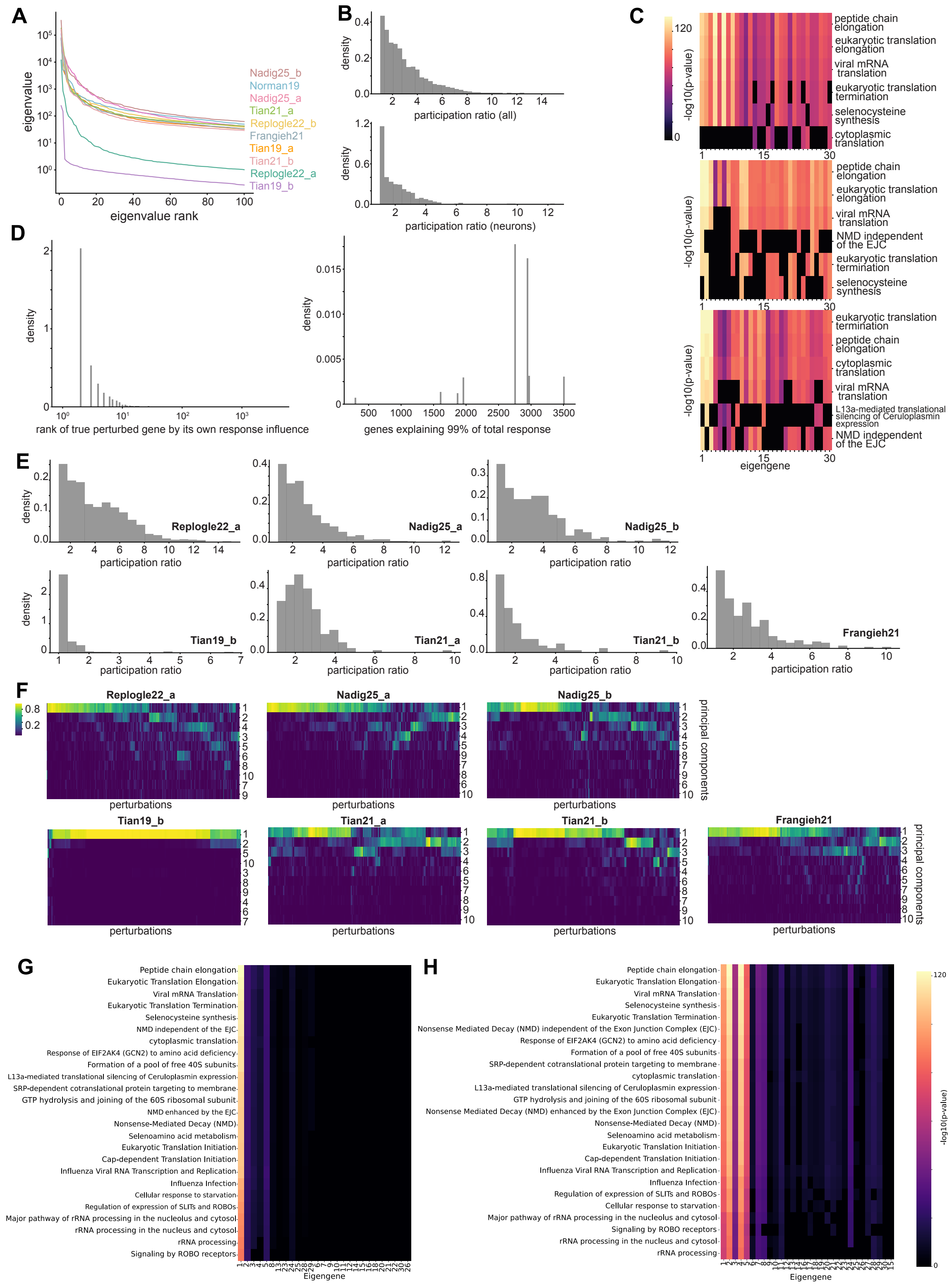
A
eigenvalue
eigenvalue rank
Nadig25_b
Norman19
Nadig25_a
Tian21_a
Replogle22_b
Frangieh21
Tian19_a
Tian21_b
Replogle22_a
Tian19_b
B
density
participation ratio (all)
participation ratio (neurons)
C
-log10(p-value)
peptide chain elongation
eukaryotic translation elongation
viral mRNA translation
eukaryotic translation termination
selenocysteine synthesis
cytoplasmic translation
NMD independent of the EJC
L13a-mediated translational silencing of Ceruloplasmin expression
eigengene
D
density
rank of true perturbed gene by its own response influence
genes explaining 99% of total response
E
density
participation ratio
Replogle22_a
Nadig25_a
Nadig25_b
Tian19_b
Tian21_a
Tian21_b
Frangieh21
F
Replogle22_a
Nadig25_a
Nadig25_b
Tian19_b
Tian21_a
Tian21_b
Frangieh21
perturbations
principal components
G
H
Peptide chain elongation
Eukaryotic Translation Elongation
Viral mRNA Translation
Eukaryotic Translation Termination
Selenocysteine synthesis
NMD independent of the EJC
Nonsense Mediated Decay (NMD) independent of the Exon Junction Complex (EJC)
cytoplasmic translation
Response of EIF2AK4 (GCN2) to amino acid deficiency
Formation of a pool of free 40S subunits
L13a-mediated translational silencing of Ceruloplasmin expression
SRP-dependent cotranslational protein targeting to membrane
GTP hydrolysis and joining of the 60S ribosomal subunit
NMD enhanced by the EJC
Nonsense Mediated Decay (NMD) enhanced by the Exon Junction Complex (EJC)
Nonsense-Mediated Decay (NMD)
Selenoamino acid metabolism
Eukaryotic Translation Initiation
Cap-dependent Translation Initiation
Influenza Viral RNA Transcription and Replication
Influenza Infection
Cellular response to starvation
Regulation of expression of SLITs and ROBOs
Major pathway of rRNA processing in the nucleolus and cytosol
rRNA processing in the nucleus and cytosol
rRNA processing
Signaling by ROBO receptors
Eigengene
-log10(p-value)

**Fig. S12**: **Additional analyses of the effective dimensionality of transcriptome-wide perturbation response.** A) Ranked covariance eigenvalue spectra for the datasets analyzed, illustrating the distribution of variance across eigengenes. B) Distributions of participation ratios across all perturbations (top) and for the neuron datasets (bottom). C) Gene ontology enrichment ($-\log_{10}(p)$) of the top 30 eigengenes for the three neuron datasets. D) Left: distribution of the rank of the true perturbed gene according to its own response influence. Right: distribution of the effective number of genes required to explain 99% of the total predicted response. E) Participation-ratio distributions shown separately for the remaining datasets not included in main text Fig. 6. F) Heatmaps showing the fraction of each perturbation response explained by the first ten principal components for the remaining datasets. G) Gene ontology enrichment ($-\log_{10}(p)$) of the top 30 eigengenes over all datasets in A). H) Gene ontology enrichment ($-\log_{10}(p)$) of the top 30 eigengenes for all neuron datasets.

### 7 Eigengene analysis

Let $\Sigma = V\Lambda V^\top$ be the eigendecomposition [33] of the covariance matrix with $V = [v_1, \ldots, v_G]$. Then, the projection coefficients which quantify the extent to which the response to perturbation j lies in the same direction as the eigenvectors are

$$\alpha_{ij} = v_i^\top \delta X_j, \tag{S172}$$

and the normalized contribution of PC $i$ to perturbation $j$ can be estimated as:

$$\tilde{\alpha}_{ij}^2 = \frac{\alpha_{ij}^2}{\sum_{k=1}^K \alpha_{kj}^2}. \tag{S173}$$

#### 7.1 Participation Ratio

Another measure of effective dimensionality of response to perturbation j is the participation ratio:

$$\mathrm{PR}_j = \frac{\left(\sum_{i=1}^K \alpha_{ij}^2\right)^2}{\sum_{i=1}^K \alpha_{ij}^4}. \tag{S174}$$

#### 7.2 GO enrichment analysis on principle components

We perform gene ontology (GO) enrichment [34] on the dominant axes of transcriptional variation across multiple Perturb-seq datasets. For each dataset, the we begin by computing the top 30 eigenvectors of the control covariance ordered by explained variance, which represent the primary directions of variation in control gene expression. These eigenvectors are interpreted by identifying the top 200 genes with the highest absolute loadings for each. These top genes are then submitted to g:Profiler for functional enrichment analysis against GO Biological Process (BP), Molecular Function (MF), and Reactome pathway (REAC) databases. For each eigengene, the top 5 significantly enriched terms (based on p-value) are retained. The results are aggregated into a matrix where rows represent GO or Reactome terms and columns represent eigengenes, with cell values reflecting enrichment strength (log p-value). This matrix provides a compact, interpretable summary of which biological processes are most associated with the dominant axes of gene expression variance in each dataset. All outputs—including the eigenvectors, gene rankings, enrichment results, and GO enrichment matrices—are saved for downstream analysis and visualization.

#### 7.3 Gene Contribution Fractions

Define the absolute propagated contribution to gene $i$:

$$c_i = \sum_{j=1}^G |\Sigma_{ij}| \cdot |u_j^*|, \tag{S175}$$

which can in general be larger than the true expression change because of possible cancellation of positive and negative correlations. The normalized global contribution vector is then

$$f_i^{(\text{global})} = \frac{c_i}{\sum_k c_k}, \tag{S176}$$

and the true perturbed target gene-specific contribution vector is

$$f_j^{(\text{target})} = \frac{|\Sigma_{gj}| \cdot |u_j^*|}{\sum_k |\Sigma_{gk}| \cdot |u_k^*|} \quad \text{where } g = \text{perturbed gene index.} \tag{S177}$$

##### 7.4 Effective Number of Contributing Genes (Entropy)

For a normalized distribution $f \in \mathbb{R}^G$, we define the entropy-based [35] effective number of genes as:

$$\text{EffSize} = \exp\left(-\sum_{i=1}^{G} f_i \log f_i\right). \tag{S178}$$

##### 7.5 Self Rank

We rank the perturbed gene's index $g$ in its own contribution vector $f^{(\text{target})}$, with genes sorted by descending $f_j$.

#### 8 Negative effects of normalizing gene expression by total counts

In short, normalization of gene expression vectors by their total mRNA count projects high-dimensional count space onto the probability simplex. This nonlinear constraint removes the "total" mRNA count degree of freedom, alters covariances, induces spurious negative correlations, and suppresses dynamical eigenmodes—particularly slow, global "soft modes." This section provides a concise, self-contained derivation of these effects.

##### 8.1 Linearizing the Change of Variables $\boldsymbol{x} \mapsto \boldsymbol{y}$

Let $x \in \mathbb{R}_+^n$ denote unnormalized gene expression for a single cell,

$$S = \sum_{k=1}^{n} x_k, \qquad y_i = \frac{x_i}{S}, \qquad \sum_i y_i = 1,$$

and let $\mu = \langle x \rangle$, $M = \sum_i \mu_i$, and $\bar{y}_i = \mu_i / M$.

Consider small perturbations $\delta x = x - \mu$ and $\delta y = y - \bar{y}$. Linearizing around $\mu$,

$$\delta y_i = \sum_j \left.\frac{\partial y_i}{\partial x_j}\right|_{x=\mu} \delta x_j.$$

Since

$$\frac{\partial}{\partial x_j}\left(\frac{x_i}{S}\right) = \frac{\delta_{ij} S - x_i}{S^2},$$

we obtain at $x = \mu$:

$$\frac{\partial y_i}{\partial x_j} = \frac{1}{M}\left(\delta_{ij} - \bar{y}_i\right).$$

Thus,

$$\boxed{\delta y_i = \frac{1}{M}\left(\delta x_i - \bar{y}_i \sum_k \delta x_k\right)} \tag{S179}$$

or in matrix form

$$\delta y = A\delta x, \qquad A = \frac{1}{M}\left(I - \bar{y}\,\mathbf{1}^\top\right).$$

#### 8.2 Total Expression Changes Are Nullified

Summing Eq. (S179) over $i$:

$$\sum_i \delta y_i = \frac{1}{M}\left(\sum_i \delta x_i - \sum_i \bar{y}_i \sum_k \delta x_k\right) = \frac{1}{M}\left(\delta S - \delta S\right) = 0.$$

Thus,

$$\boxed{\sum_i \delta y_i = 0, \quad \text{i.e. the normalization removes all fluctuations in total mRNA.}}$$

An infinitesimal global scaling $\delta x = \varepsilon\mu$ gives

$$\delta y_i = \frac{\varepsilon}{M}\left(\mu_i - \bar{y}_i M\right) = 0,$$

so the entire "radial" direction proportional to $\mu$ is annihilated.

Hence normalization eliminates any mode (dynamical or statistical) that acts primarily by changing total expression.

#### 8.3 Spurious Negative Correlations on the Simplex

Let $\Sigma_x = \langle \delta x \delta x^\top \rangle$ and $\Sigma_y = A\Sigma_x A^\top$. Even if the $x_i$ are independent:

$$\Sigma_x = \mathrm{diag}(\sigma_1^2, \ldots, \sigma_n^2),$$

normalization induces off-diagonal structure.

From Eq. (S179):

$$\delta y_i = \frac{1}{M}(\delta x_i - \bar{y}_i \delta S), \qquad \delta S = \sum_k \delta x_k,$$

so for $i \neq j$:

$$\boxed{\mathrm{Cov}(\delta y_i, \delta y_j) = \frac{1}{M^2}\left(-\bar{y}_i\sigma_j^2 - \bar{y}_j\sigma_i^2 + \bar{y}_i\bar{y}_j\sum_k \sigma_k^2\right).} \qquad \text{(S180)}$$

For example, If all means and variances are equal:

$$\bar{y}_i = \frac{1}{n}, \qquad \sigma_i^2 = \sigma^2,$$

then Eq. (S180) becomes

$$\mathrm{Cov}(\delta y_i, \delta y_j) = -\frac{\sigma^2}{nM^2} < 0.$$

Thus, normalization alone produces negative correlations even when the unnormalized genes fluctuate independently.

##### 8.3.1 Separating Total and Composition Directions

Introduce an invertible linear change of variables:

$$z = Tx = \begin{pmatrix} S \\ w \end{pmatrix},$$

where $S = \mathbf{1}^\top x$ and $w \in \mathbb{R}^{n-1}$ spans the orthogonal complement of $\mathbf{1}$.

The linearized dynamics in $x$-space

$$\delta\dot{x} = J\delta x + \eta$$

becomes

$$\delta\dot{z} = J_z \delta z + \eta_z, \qquad J_z = TJT^{-1}, \qquad \eta_z = T\eta.$$

In block form:

$$J_z = \begin{pmatrix} J_{SS} & J_{Sw} \\ J_{wS} & J_{ww} \end{pmatrix}.$$

#### 8.4 Correlations Between Total mRNA and Compositional degrees of freedom

Using Eq. (S179), the cross-covariance is

$$\mathrm{Cov}(\delta S, \delta y_i) = \frac{1}{M^2}\left(\sigma_i^2 - \bar{y}_i \sum_k \sigma_k^2\right).$$

***Symmetric system.***

If all $\mu_i$ and $\sigma_i^2$ are equal, this evaluates to 0:

$$\mathrm{Cov}(\delta S, \delta y_i) = 0.$$

Total and composition are uncorrelated at linear order.

***Heterogeneous system.***

If $\sigma_i^2$ and $\bar{y}_i$ vary—which is always the case in real data—the cross-covariance is generically nonzero and often negative:

$$\sigma_i^2 < \bar{y}_i \sum_k \sigma_k^2 \quad \Rightarrow \quad \mathrm{Cov}(\delta S, \delta y_i) < 0.$$

As a consequence, highly expressed or high-variance genes absorb a disproportionate fraction of the total mRNA fluctuations. The normalization term $-\bar{y}_i \delta S/M$ therefore pushes their relative frequencies downward when the total expression $S$ is high and upward when $S$ is low. This mechanism induces a genuine statistical coupling between fluctuations in total expression $\delta S$ and compositional shifts $\delta y_i$, even in the absence of any underlying regulatory interaction.

#### 8.5 The Total-Direction Eigenvector

In typical transcriptional systems, one eigenvector is closely aligned with the "total expression" direction $\mathbf{1}$ or equivalently with the mean expression vector $\mu$. Under normalization, the projection operator onto the probability simplex,

$$P = I - \bar{y}\,\mathbf{1}^\top,$$

annihilates this direction exactly, since

$$P\mathbf{1} = 0.$$

As a result, the dynamical mode associated with fluctuations in total expression is removed entirely from the observable dynamics in the normalized variables $y$.

#### Loss of dominant collective modes

If the total-direction eigenvector corresponds to a small-magnitude eigenvalue—that is, to a global dominant collective mode—then this mode does not survive the projection induced by normalization, because it does not exist on the simplex. Moreover, any dynamical mode with substantial overlap with the total-expression direction is strongly attenuated by the projection. Consequently, normalized data systematically under-represent slow global modes of transcriptional dynamics. The effective Jacobian governing compositional fluctuations,

$$J_{\mathrm{comp}} \approx PJP,$$

therefore exhibits a spectrum with one fewer eigenvalue, reduced variances along the retained directions, and typically more negative (stiffer) inferred relaxation rates.

#### 8.6 PFlog variance stabilization

To further test whether CIPHER performance depends on the representation of expression values, we repeated the forward-prediction analysis using a PFlog-transformed expression space. PFlog is a shifted, centered log transform designed to reduce the strong mean–variance dependence of raw single-cell count data while preserving covariance structure among genes. For each dataset, the transformation was fit directly from raw control cells, and all downstream quantities used for CIPHER forward prediction were recomputed in the transformed space.

For each gene $g$, we first estimated the raw control-cell mean and variance,

$$\mu_g = \langle X_g \rangle_0, \qquad v_g = \mathrm{Var}_0(X_g), \tag{S181}$$

where $\langle \cdot \rangle_0$ denotes an average over control cells. We then fit a negative-binomial-style mean–variance trend,

$$v_g \approx \mu_g + \alpha \mu_g^2, \tag{S182}$$

using binned median mean–variance trend points rather than individual genes. This fit estimates the typical control-cell dispersion structure while reducing sensitivity to outlier genes. The fitted dispersion parameter defines a dataset-specific pseudocount,

$$p_c = \frac{1}{4\alpha}. \tag{S183}$$

Raw counts were then transformed as

$$Y_{ig} = \log\left(X_{ig} + p_c\right) - \frac{1}{G}\sum_{h=1}^{G} \log\left(X_{ih} + p_c\right), \tag{S184}$$

where $i$ indexes cells and $G$ is the number of genes. Thus, $Y_i$ is a shifted, centered-log expression vector for each cell. The shift stabilizes low-count genes, while the centering removes cell-wide multiplicative effects such as differences in capture efficiency or library size.

After transforming the data, we recomputed the control-cell covariance matrix in PFlog space,

$$\Sigma^{\mathrm{PFlog}} = \mathrm{Cov}_0(Y). \tag{S185}$$

For each single-gene perturbation targeting gene $g$, the observed PFlog response was

$$\Delta Y_g = \langle Y \rangle_g - \langle Y \rangle_0, \tag{S186}$$

where $\langle Y \rangle_g$ is the mean PFlog expression vector among cells assigned to perturbation $g$. CIPHER forward prediction was then evaluated using the $g$-th column of the PFlog control covariance matrix,

$$\widehat{\Delta Y}_g = a_g \Sigma^{\mathrm{PFlog}}_{:g}, \qquad a_g = \frac{\left(\Sigma^{\mathrm{PFlog}}_{:g}\right)^T \Delta Y_g}{\left\| \Sigma^{\mathrm{PFlog}}_{:g} \right\|^2}. \tag{S187}$$

Prediction accuracy was quantified by the Pearson correlation between the observed response $\Delta Y_g$ and the predicted response $\widehat{\Delta Y}_g$ across genes.

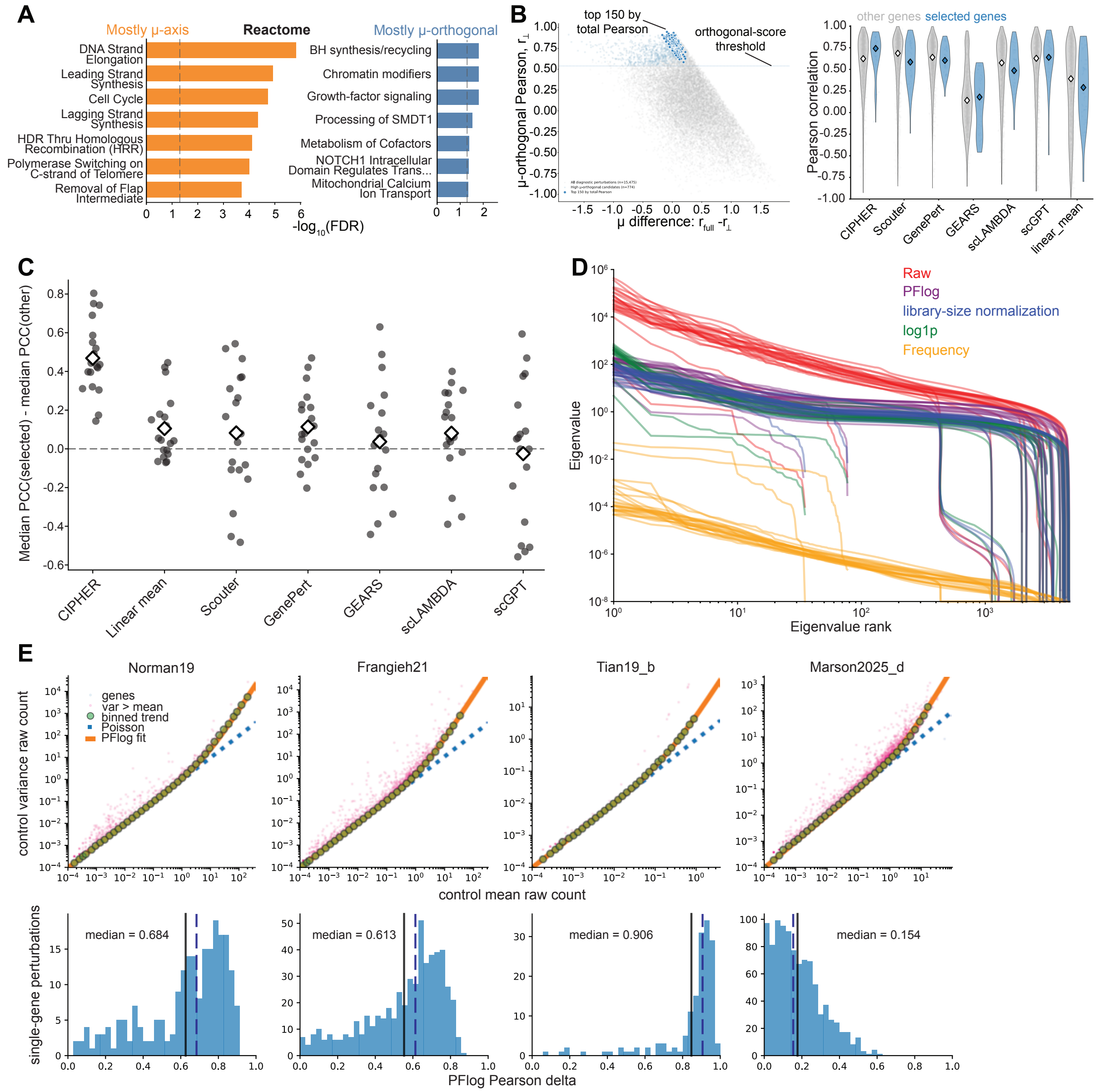


**Fig. S13**: **Effects of normalization methods.** A) Reactome gene-set enrichment analysis for perturbations whose responses are primarily aligned with the mean-expression ($\mu$) axis (left) or primarily orthogonal to the $\mu$ axis (right). The top seven enriched pathways are shown for each group. B) Distributions of Pearson correlations between predicted and observed responses for perturbations targeting genes with strong $\mu$-orthogonal responses (blue) compared with all other perturbations (gray), shown for each method. C) Difference between the median Pearson correlation of the $\mu$-aligned perturbations and that of the remaining perturbations within each dataset. Points correspond to datasets and open diamonds denote dataset means. D) Covariance eigenspectra across normalization methods (colors) and datasets. E) Top row: control-cell mean–variance relationships for four representative datasets. Each point is a gene, green points denote binned median mean–variance trends, the blue dashed line is the Poisson expectation ($v = \mu$), and the orange curve is the fitted negative-binomial mean–variance relation ($v = \mu + \alpha\mu^2$) used to determine the PFlog pseudocount. Bottom row: distributions of PFlog-space CIPHER forward-prediction performance across single-gene perturbations, measured by the Pearson correlation between predicted and observed PFlog responses. Dashed blue lines indicate the median Pearson correlation and solid black lines indicate the mean. Scores have not been converted back to raw.

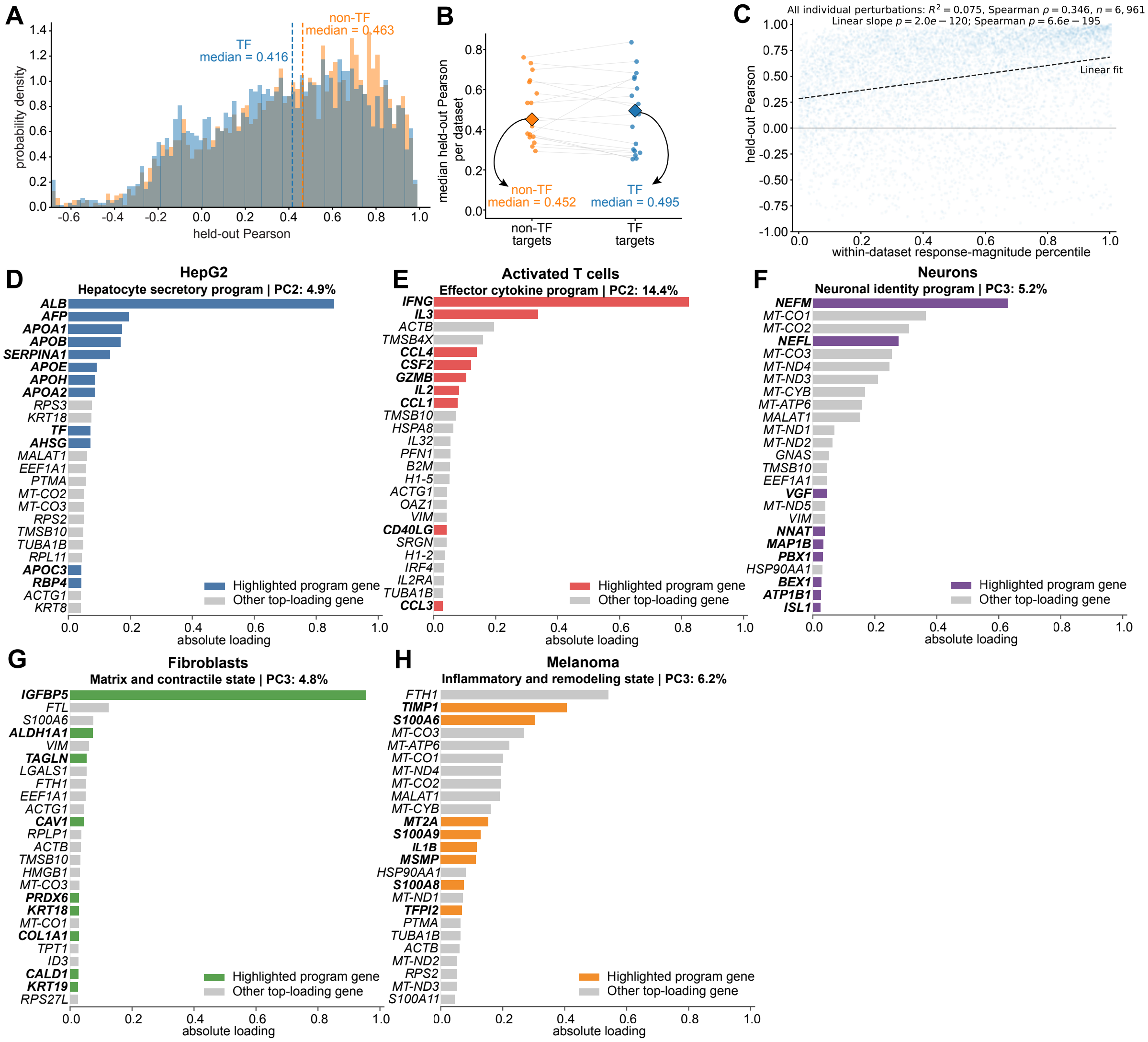


**Fig. S14**: **Transcription factor perturbations and covariance-derived cell programs.** (A) Distribution of held-out Pearson correlations for transcription factor (TF) and non-transcription factor (non-TF) perturbations. Dashed lines indicate the median of each group. (B) Median held-out Pearson correlation for TF and non-TF perturbations across datasets, with paired datasets connected by gray lines. (C) Held-out Pearson correlation as a function of within-dataset response-magnitude percentile for all perturbations. he panel shows the stats ($R^2$=0.075, Spearman =0.346, n=6,961, slope/Spearman p-values) (D–H) Top absolute gene loadings for representative covariance principal components from HepG2, activated T cells, neurons, fibroblasts, and melanoma cells, with genes associated with the highlighted transcriptional program shown in color. Titles indicate the inferred program, principal component, and fraction of covariance variance explained.

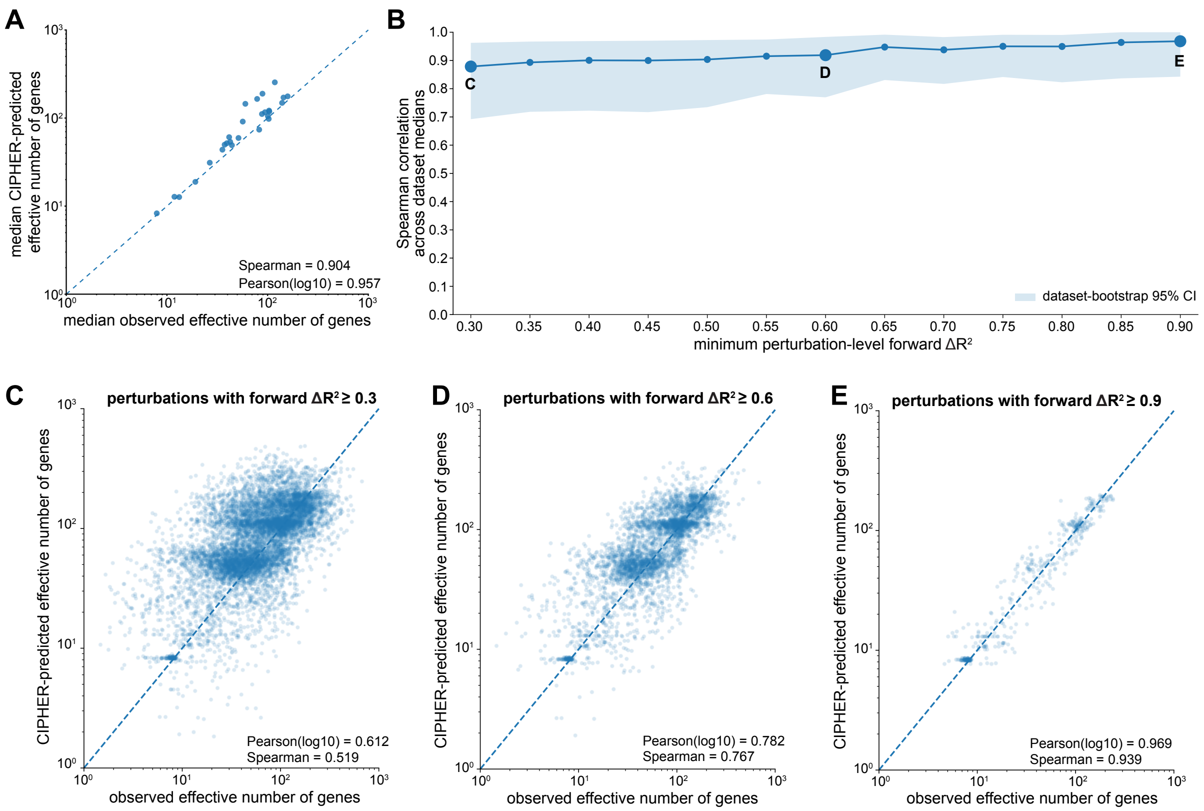


**Fig. S15**: **Prediction of the effective dimensionality of perturbation responses.** A) Median CIPHER-predicted effective number of genes versus the median observed effective number of genes across datasets for perturbations with minimum $\Delta R^2 = 0.5$. Each point corresponds to a dataset. B) Spearman correlation between predicted and observed median effective numbers of genes across datasets as a function of the minimum forward-prediction $\Delta R^2$ threshold used to include perturbations. Shaded region denotes the bootstrap 95% confidence interval across datasets. C-E) Predicted versus observed effective number of genes for individual perturbations passing minimum forward-prediction thresholds of $\Delta R^2 \geq 0.3$ (C), $\Delta R^2 \geq 0.6$ (D), and $\Delta R^2 \geq 0.9$ (E). Dashed lines indicate equality between prediction and observation.

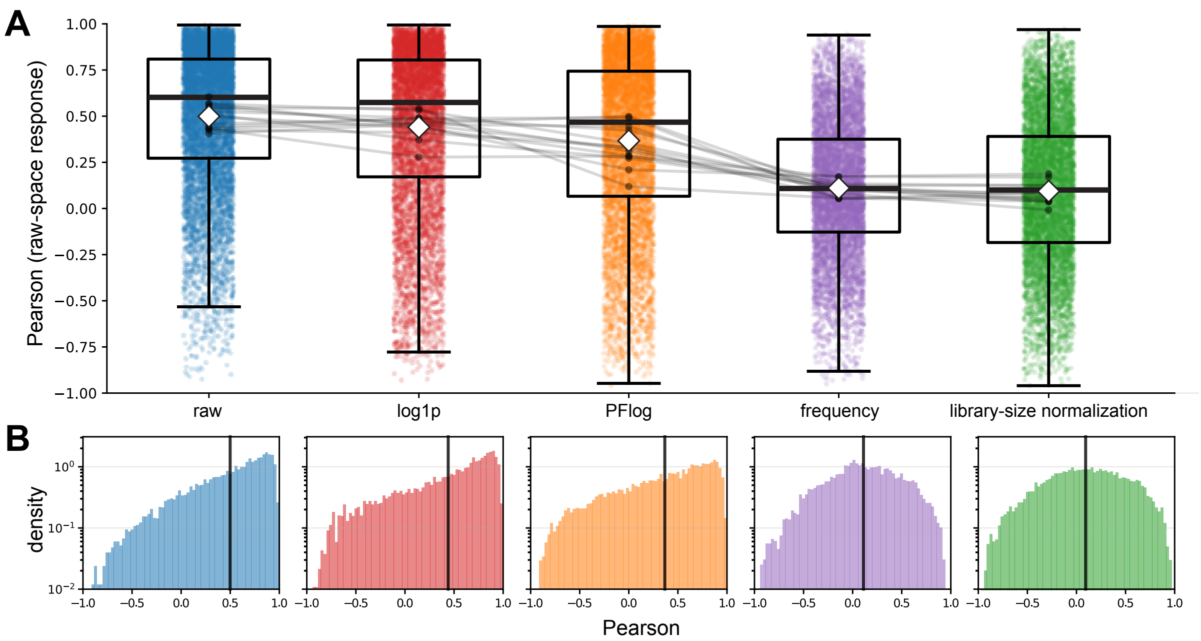


**Fig. S16**: **Effects of normalization on linear-response prediction after mapping back to raw expression space.** (A) Held-out Pearson correlations between predicted and observed perturbation responses after transforming predictions from each normalization back to raw expression space using a linear approximation to the inverse Jacobian. Points denote individual perturbations, gray lines connect the same dataset across normalization methods, boxplots summarize the distributions, and white diamonds indicate the mean. (B) Corresponding Pearson correlation distributions for each normalization method. Vertical black lines denote the mean Pearson correlation.

The top row of Fig. S13 verifies the PFlog fitting procedure. In all four representative datasets, the raw control-cell mean–variance relationship is strongly non-Poisson and is well captured by the fitted quadratic trend over the expression range used for analysis. This confirms that the PFlog pseudocount is estimated from dataset-specific count statistics rather than imposed as a fixed global constant. The bottom row shows that PFlog-space covariance retains predictive information about perturbation responses, although the strength of prediction remains dataset-dependent. The TianKampmann2019 iPSC dataset shows especially high PFlog-space predictability, whereas the Marson2025 stimulated T-cell dataset shows weaker performance. Overall, these results show that the covariance-based CIPHER signal is not restricted to raw-count coordinates, but persists after a variance-stabilizing, centered-log transformation of the data.

### 9 Bayesian inverse CIPHER identifies candidate drivers of resistance to targeted cancer therapy

We next applied the Bayesian inverse form of CIPHER to cancer drug-resistance gene expression, where the goal is not to predict the known perturbation in a Perturb-seq experiment, but to infer a parsimonious set of candidate driver genes whose covariance-propagated effects can explain the observed transition from a drug-naive state to a drug-resistant state. For each naive-resistant expression difference, we defined

$$\Delta X = \langle X \rangle_{\text{Resistant}} - \langle X \rangle_{\text{Naive}}\,, \tag{S188}$$

and estimated the covariance matrix $\Sigma_0$ from the naive population. We then used the linear-response observation model

$$\Delta X = \Sigma_0 u + \epsilon, \qquad \epsilon \sim \mathcal{N}(0, H), \tag{S189}$$

with a Gaussian prior on the latent perturbing force,

$$u \sim \mathcal{N}(0, \tau^2 I). \tag{S190}$$

The resulting posterior is Gaussian, with precision and mean

$$\Lambda_{\text{post}} = \Sigma_0^T H^{-1} \Sigma_0 + \tau^{-2} I, \qquad \mu_{\text{post}} = \Lambda_{\text{post}}^{-1} \Sigma_0^T H^{-1} \Delta X. \tag{S191}$$

The posterior mean $\mu_{\text{post},g}$ is therefore a covariance-corrected driver score for gene $g$: it measures the inferred force on that gene after accounting for how perturbations propagate through the naive-state covariance structure. This differs from log fold change, which measures only the marginal expression shift of each gene between resistant and naive cells.

For the KRAS pancreatic MP2 analysis, we used raw expression counts and selected the top 2,000 genes satisfying BH-FDR $< 0.05$ and $|\log_2 \text{FC}| \geq 0.5$, ranked by absolute log fold change. Ribosomal, mitochondrial, heat-shock, and translation-initiation-factor genes were removed before inference. We used the naive covariance matrix with shrinkage $10^{-6}$, a naive sample-mean noise model $H$, ridge stabilization $10^{-6}$, and $\tau^2 = 1$. This value was chosen as a weakly regularizing prior variance: it is large enough to allow a multigene resistance force, but still prevents the inverse problem from becoming completely unregularized. In this setting, $\tau^2 = 1$ represents a predictive-plateau compromise: larger values can improve direct reconstruction of $\Delta X$, but they also increase posterior force magnitudes and make the model more capable of fitting null or non-specific expression differences.

For the melanoma naive-to-resistant expression difference, we used a more conservative prior variance, $\tau^2 = 10^{-6}$. This value was chosen deliberately for high-specificity driver attribution, not because it maximized the marginal likelihood. The marginal likelihood favored much larger $\tau^2$ values, approaching an almost unregularized reconstruction regime in which the posterior can pass nearly all selected-gene modes and closely interpolate the observed mean shift. Although useful as a reconstruction limit, that regime is less appropriate for identifying robust biological drivers because it can also fit naive-vs-naive null expression differences. In contrast, $\tau^2 = 10^{-6}$ acts as a strong minimum-intervention prior: it suppresses spurious high-dimensional forces and restricts inference to the dominant covariance-aligned melanoma resistance axis. Thus, the melanoma analysis prioritizes specificity and null calibration, whereas the KRAS MP2 analysis uses a less restrictive prior to allow a broader multigene resistance program.

The posterior mean was compared with conventional differential expression by plotting $\log(1+\mu_{\text{post},g})$ against $\log_2 \text{FC}_g$. Genes in the upper-right portion of this plot are both strongly upregulated in resistant cells and assigned high covariance-corrected posterior driver scores. In the case of KRAS MP2, genes such as *ANKRD1*, *IGFN1*, *SAMD9*, *PSG9*, and *PSG4* appear as high-scoring candidates, indicating agreement between marginal resistance-associated expression changes and the inverse CIPHER posterior. Genes that deviate from the main log-fold-change trend are also informative, because they represent cases where covariance structure either amplifies or suppresses the inferred driver score relative to what would be expected from differential expression alone.

#### 9.1 Experimental validation in melanoma and MIA PaCa-2

#### Cell lines and culture

The melanoma line WM989 A6-G3 (BRAF-V600E), described in (https://pmc.ncbi.nlm.nih.gov/articles/PMC10628994/) was obtained from the laboratory of Arjun Raj (University of Pennsylvania) and derived by twice single-cell bottlenecking WM989. MP2-E7 was derived by single-cell bottlenecking the KRAS-G12C-mutant PDAC line MIA PaCa-2 (provided by the laboratory of Beatriz Sosa-Pineda, Northwestern University). The identity of MIA PaCa-2 was verified by ATCC human cell STR profiling.

MP2-E7 was cultured in DMEM (Corning, MT10017CM) supplemented with 10% FBS (Gibco, 16000044) and 1% penicillin-streptomycin (Gibco, 15070063). WM989 A6-G3 was cultured in TU2% medium (80% MCDB 153, 10% Leibovitz's L-15, 2% FBS, 2.4 mM $CaCl_2$, 50 U ml$^{-1}$ penicillin, 50 $\mu$g ml$^{-1}$ streptomycin). MP2-E7 was passaged with 0.25% trypsin-EDTA (Gibco, 25200056) and WM989 A6-G3 with 0.05% trypsin-EDTA (Gibco, 25300054).

#### Single-cell RNA sequencing of naive and resistant populations

To generate resistant populations, cells were seeded sparsely (<5% confluency), allowed to adhere ~16 h, and treated with drug: MP2-E7 with 8 μM sotorasib (AMG510; SelleckChem, S8830), and WM989 A6-G3 with 5 nM trametinib in TU2% medium. Medium and drugs were replaced twice weekly, and treatment continued for 3-4 weeks until resistant colonies emerged. Therapy-naive and resistant populations were dissociated, resuspended in 0.1% BSA in PBS, filtered to a single-cell suspension, and processed on the 10x Chromium platform and sequenced.

#### Lentivirus packaging and transduction

Lentivirus was packaged in-house following a previously described protocol (https://pmc.ncbi.nlm.nih.gov/articles/PMC10628994/). HEK293FT cells were grown to 80-95% confluency in 10-cm plates in DMEM with 10% FBS and 1% penicillin-streptomycin; one day before transfection the medium was replaced with DMEM/10% FBS without antibiotics. Per plate, 80 μl polyethylenimine (Polysciences, 23966) in 500 μl Opti-MEM was combined with 5 μg VSV-G, 7.5 μg psPAX2, and ~7.35 μg transfer plasmid in 500 μl Opti-MEM; solutions were incubated separately for 5 min, mixed, incubated 15 min, and added dropwise. Medium was refreshed after 6-7 h, and virus-laden medium was collected every 9-11 h over ~30 h, filtered (0.45-μm PES), and stored at −80 °C.

#### shRNA knockdown

Individual genes were knocked down using GIPZ lentiviral shRNA constructs (Dharmacon) obtained from the Gene Editing, Transduction and Nanotechnology (GET iN) Core of the Northwestern University Skin Biology & Diseases Resource-Based Center. Plasmid was prepared by endotoxin-free maxiprep (Qiagen EndoFree Plasmid Maxi Kit) and packaged into lentivirus in-house as above. Two independent shRNA constructs were used per gene, with a non-targeting scrambled construct as control. Constructs were generated against *ANKRD1*, *IGFN1*, *SAMD9*, *TSPAN12*, *TNNT2*, *ATXN1*, *PSG4*, *PSG9*, *TGFA*, *ROBO2*, *ATRNL1*, *DACT1*, *CDKN1A*, *GBP3*, *SYBU*, *MAPK4*, *PARP8*, *APOL1*, *STX11*, and *HS6ST3*. Following transduction, GFP-positive cells were isolated by fluorescence-activated cell sorting.

For transduction, cells were seeded at 50,000 per well of a 6-well plate with virus, 4 μg ml$^{-1}$ polybrene, and medium to 2 ml, then spinfected at 1,700 rpm for 25 min. Polybrene-containing medium was removed after ~7 h. Transduced (GFP-positive) cells were isolated by fluorescence-activated cell sorting, expanded, and cryopreserved. Virus volume per transduction was determined by titration on each line.

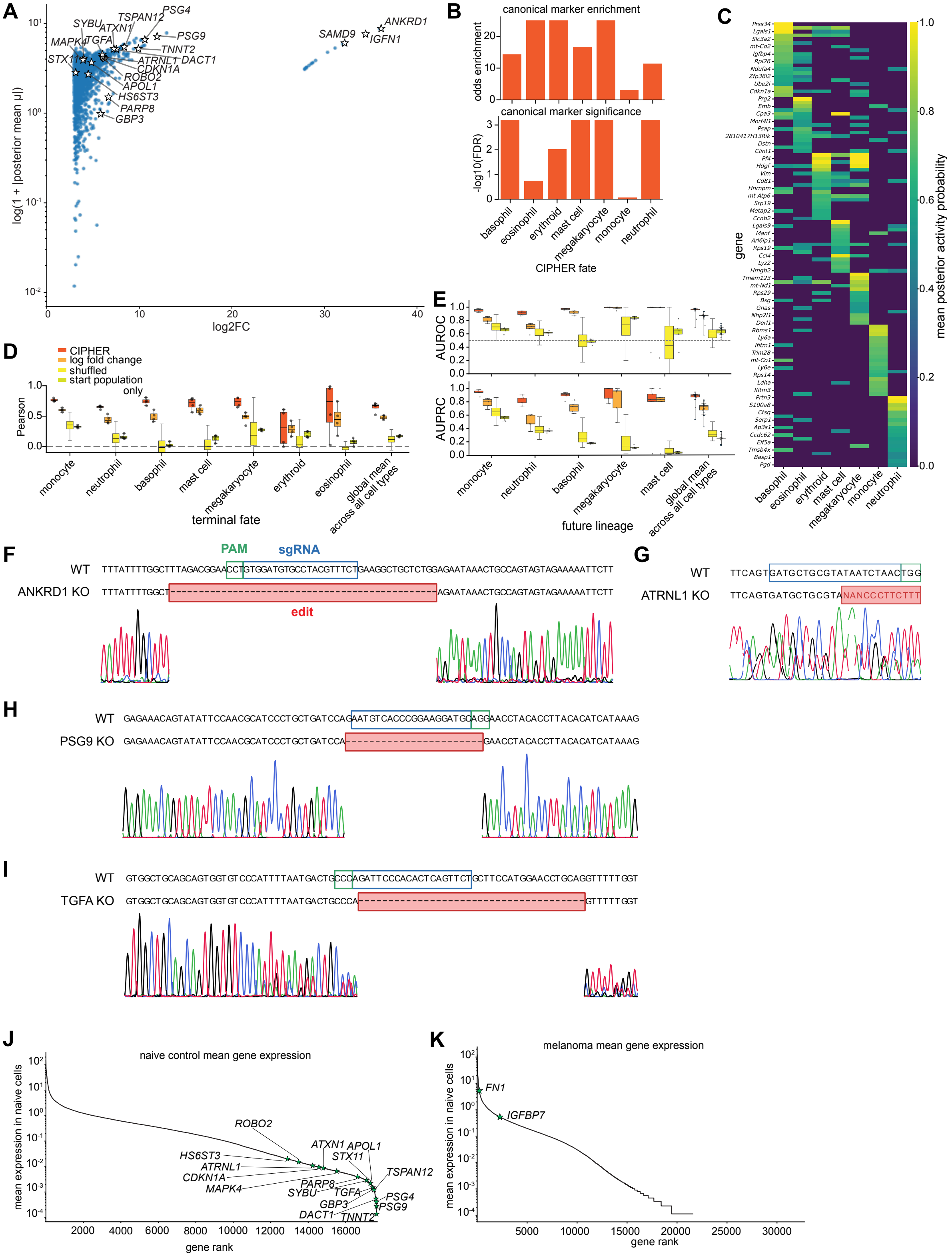
A
log(1 + |posterior mean μ|)
log2FC
SYBU
TSPAN12
PSG4
ATXN1
MAPK4
TGFA
PSG9
TNNT2
STX11
ATRNL1
DACT1
CDKN1A
ROBO2
APOL1
HS6ST3
PARP8
GBP3
ANKRD1
SAMD9
IGFN1
B
canonical marker enrichment
odds enrichment
canonical marker significance
-log10(FDR)
basophil
eosinophil
erythroid
mast cell
megakaryocyte
monocyte
neutrophil
CIPHER fate
C
gene
mean posterior activity probability
D
CIPHER
log fold change
shuffled
start population only
Pearson
terminal fate
global mean across all cell types
E
AUROC
AUPRC
future lineage
F
PAM
sgRNA
WT
ANKRD1 KO
edit
G
ATRNL1 KO
H
PSG9 KO
I
TGFA KO
J
naive control mean gene expression
mean expression in naive cells
gene rank
K
melanoma mean gene expression
FN1
IGFBP7

**Fig. S17**: **Biological applications supplementary figures.** A) Each point represents one selected gene from the Naive-to-Resistant expression difference. The $x$-axis shows the conventional $\log_2$ fold change between resistant and naive cells, and the $y$-axis shows the posterior CIPHER score, $\log(1 + \mu)$, where $\mu$ is the posterior mean of the inferred perturbing force under the Gaussian inverse model. The analysis used the top 2,000 significant high-log-fold-change genes, raw expression counts, the naive-state covariance matrix, a naive sample-mean noise model, and prior variance $\tau^2 = 1$. Highlighted genes mark candidate resistance-associated drivers with large posterior scores. Genes in the upper-right quadrant, including *ANKRD1*, *IGFN1*, *SAMD9*, *PSG9*, and *PSG4*, are both strongly upregulated and assigned high covariance-corrected posterior force. B) Enrichment of canonical lineage marker sets among the top CIPHER-ranked genes for each inferred fate force. The top panel shows the odds enrichment for each lineage marker set, and the bottom panel shows the corresponding FDR-adjusted significance. C) Heatmap of Bayesian CIPHER posterior activity probabilities for top fate-associated gene perturbations. High posterior activity indicates genes with strong evidence for nonzero contribution to a fate-specific early-state force. D) Prediction of terminal clone composition from day 4 undifferentiated clone states. For each held-out clone, CIPHER scores predicted day 6 terminal fate fractions and compared with the observed terminal fate fractions. Performance is shown as the Pearson correlation between predicted and observed clone fractions for each fate, together with the global mean across fates. Shuffled fate-label controls, start-population-only predictions, and terminal marker log-fold-change scores are also shown. E) Dominant future-fate classification from day 4 undifferentiated clone states. For each lineage, clones were scored for future fate commitment and evaluated using AUROC and AUPRC. F-I) Sanger sequencing validation of CRISPR-Cas9 knockouts at endogenous loci in MP2-E7 (MIA PaCa-2) cells. For each gene, the wild-type (WT) reference sequence is aligned above the edited knockout (KO) allele, with the sgRNA target and PAM annotated on the WT sequence and the resulting deletion shown as a boxed gap; chromatograms below show the Sanger traces spanning the cut site. Frameshifting deletions confirm loss-of-function editing for F) *ANKRD1*, G) *ATRNL1*, H) *PSG9*, and I) *TGFA*. J-K) Ranked mean gene expression in naive control cells, with validated resistance-associated genes labeled. J) Naive MIA PaCa2 control cells, highlighting CIPHER-nominated drivers across the expression-rank distribution. K) Naive WM989 (BRAF-V600E) melanoma control cells, highlighting the top-ranked resistance mediators *FN1* and *IGFBP7*.

#### CRISPR-Cas9 knockout

*ANKRD1*, *PSG9*, *ATRNL1*, and *TGFA* were disrupted in MP2-E7 using the pSpCas9(BB)-2A-GFP system (pX458; Addgene #48138). For each gene, three to five sgRNAs targeting distinct exons were designed with CRISPick (https://portals.broadinstitute.org/gppx/crispick/public, SpCas9-NGG), excluding guides with predicted off-target matches; guide oligos (IDT) were annealed and cloned into the BbsI-digested vector, and insert identity was confirmed by Sanger sequencing (U6 primer). Cells were transfected with 2 $\mu$g plasmid per well of a 6-well plate using jetOPTIMUS (Polyplus), co-transfecting multiple guides per gene, with a non-targeting sgRNA as control. At 48 h, GFP-positive cells were single-cell sorted into 96-well plates and expanded into clonal populations.

Clonal knockouts were confirmed by genomic DNA isolation, PCR amplification across the cut site, and Sanger sequencing (GENEWIZ/Azenta Life Sciences and ACGT DNA sequencing services). Where direct sequencing yielded overlapping traces from compound indels, amplicons were TA-cloned (Qiagen pDrive) and individual clones sequenced to resolve each allele. Clones carrying frameshifting deletions were retained for drug-resistance assays.

#### Resistance colony-formation assay

Knockdown or knockout cells were thawed, seeded sparsely and, after 12-16 h, treated with drug (8.5 $\mu$M sotorasib for MP2-E7; [5 nM trametinib] for WM989 A6-G3). Medium and drug were replaced twice weekly, and colony formation was monitored by Incucyte live-cell imaging over [3-4 weeks]. Resistant colonies were quantified [manually in FIJI]. Colony counts were normalized within each experiment to the scrambled (shRNA) or non-targeted (CRISPR) control. Data are from n = 2 biological replicates; bars show the mean and individual points show replicates.

### 10 Cell fate mapping in LARRY barcoding dataset

To evaluate whether early transcriptional states encode future hematopoietic fate commitment, we analyzed the LARRY clonal lineage dataset by separating cells according to experimental day and using clonal barcodes to connect early progenitors to their descendants. Day 4 undifferentiated cells were used as the starting population, and day 6 differentiated cells were used to define terminal lineage outcomes. For each clone, day 4 cells provided the early expression state, while associated day 6 descendants provided either a terminal fate-fraction vector or a dominant future lineage label. We then estimated the covariance structure of the day 4 undifferentiated starting population and used CIPHER to infer covariance-corrected fate scores for each terminal lineage. In contrast to a standard terminal marker score, which projects early cells onto genes differentially expressed in day 6 terminal populations, the CIPHER score identifies early expression directions that are predictive of future fate after accounting for correlated variation in the starting population.

Let $E_c$ denote the set of day 4 undifferentiated cells belonging to clone $c$, and let $T_c$ denote its day 6 terminal descendants. The early expression state of clone $c$ is defined as

$$X_c^{(4)} = \frac{1}{|E_c|} \sum_{i \in E_c} X_i, \tag{S192}$$

where $X_i$ is the gene-expression vector of cell $i$. The observed terminal fate composition of the same clone is

$$Y_{cf} = \frac{N_{cf}}{\sum_{f'} N_{cf'}}, \qquad N_{cf} = \#\{i \in T_c : \text{fate}(i) = f\}. \tag{S193}$$

Thus, $X_c^{(4)}$ is the prospective day 4 measurement, whereas $Y_{cf}$ is the future day 6 outcome.

For each terminal fate $f$, we constructed a fate-associated early expression contrast using only day 4 undifferentiated clone states. In the terminal-composition analysis, this contrast was defined as

$$\Delta X_f = \frac{\sum_{c \in \mathcal{C}_{\text{train}}} Y_{cf} X_c^{(4)}}{\sum_{c \in \mathcal{C}_{\text{train}}} Y_{cf}} - \frac{\sum_{c \in \mathcal{C}_{\text{train}}} (1 - Y_{cf}) X_c^{(4)}}{\sum_{c \in \mathcal{C}_{\text{train}}} (1 - Y_{cf})}. \tag{S194}$$

In the dominant-fate analysis, this reduces to the future-fate-versus-rest contrast

$$\Delta X_f = \left\langle X^{(4)} \right\rangle_{\text{future } f} - \left\langle X^{(4)} \right\rangle_{\text{future } /f}. \tag{S195}$$

This is the lineage-tracing analogue of the perturbation contrast $\Delta X = \langle X \rangle_u - \langle X \rangle_0$ in Eq. (S65). However, unlike a terminal marker contrast, both averages in Eq. (S195) are taken over day 4 undifferentiated cells. The day 6 cells are used only to assign the future fate label or fate-fraction vector.

The covariance matrix was estimated from the day 4 undifferentiated starting population,

$$\Sigma_4 = \text{Cov}\left(X_i : i \in \text{day 4 undifferentiated}\right). \tag{S196}$$

CIPHER then assumes the linear-response relation

$$\Delta X_f \approx \Sigma_4 u_f, \tag{S197}$$

where $u_f$ is a fate-specific force. In the simplest point-estimate form,

$$u_f = \Sigma_4^{-1} \Delta X_f. \tag{S198}$$

In practice, $u_f$ was estimated with regularization or Bayesian posterior shrinkage to stabilize inference in the high-dimensional setting.

Given an inferred force $u_f$, we define the raw CIPHER score of held-out clone $c$ for fate $f$ as

$$R_{cf} = \left(X_c^{(4)}\right)^T u_f. \tag{S199}$$

This score measures the alignment between the previously unseen clone's early transcriptional state and the inferred future-fate direction from the training clones. To obtain a likelihood-based fate score, we used the Gaussian linear-response log enrichment

$$L_{cf} = \left(X_c^{(4)}\right)^T u_f - \frac{1}{2} u_f^T \Sigma_4 u_f. \tag{S200}$$

The first term in Eq. (S200) rewards alignment with the fate-associated early expression direction, while the second term penalizes forces that are large under the day 4 covariance. Thus, $L_{cf}$ is a covariance-corrected prospective fate score, rather than a direct terminal marker score.

To convert fate scores into predicted terminal fate fractions, we normalized the fate logits across lineages using a temperature-calibrated softmax,

$$\widehat{Y}_{cf}(T) = \frac{\exp\left(L_{cf}/T\right)}{\sum_{f'} \exp\left(L_{cf'}/T\right)}. \tag{S201}$$

Here, $T$ is a calibration temperature: larger $T$ produces softer fate-fraction predictions, whereas smaller $T$ produces more decisive dominant-fate predictions. We selected $T$ on training clones by minimizing the multinomial negative log likelihood of the observed day 6 fate counts,

$$\mathcal{L}_{\mathrm{NLL}}(T) = -\sum_{c \in \mathcal{C}_{\mathrm{train}}} \sum_f N_{cf} \log \widehat{Y}_{cf}(T). \tag{S202}$$

Held-out clones were then evaluated either as full fate-composition predictions, using $\widehat{Y}_{cf}$ compared with $Y_{cf}$, or as dominant-fate predictions,

$$\widehat{f}_c = \arg\max_f \widehat{Y}_{cf}. \tag{S203}$$

For the Bayesian CIPHER analysis, we treated the observed fate contrast as a noisy linear-response measurement,

$$\Delta X_f = \Sigma_4 u_f + \epsilon_f, \qquad \epsilon_f \sim \mathcal{N}(0, H_f), \tag{S204}$$

with a Gaussian prior on the fate force,

$$u_f \sim \mathcal{N}(0, \tau^2 I). \tag{S205}$$

This gives an analytic Gaussian posterior over $u_f$. For each gene $g$ and fate $f$, we summarized this posterior by the mean force $\mu_{fg}$, posterior standard deviation $\sigma_{fg}$, and posterior activity probability

$$\mathrm{PIP}_{fg} = P\left(|u_{fg}| > u_{\min} \mid \Delta X_f, \Sigma_4, H_f\right). \tag{S206}$$

Genes with high posterior activity were interpreted as candidate early regulators or markers of the corresponding fate-associated force.

The resulting scores revealed that day 4 transcriptional variation in undifferentiated cells contains prospective information about day 6 fate outcomes. In Fig. S17D, predicted fate fractions were compared with observed terminal clone compositions across held-out clones. CIPHER achieved positive correlations across multiple lineages, including monocyte, neutrophil, basophil, mast cell, megakaryocyte, erythroid, and eosinophil outcomes, and outperformed shuffled-fate and starting-population-only controls, as well as log-fold change terminal marker scores. Fig. S17E shows the complementary dominant-fate classification analysis, where CIPHER generally produced higher AUROC and AUPRC than terminal marker scores or null controls. This indicates that covariance-corrected early-state directions provide stronger prospective fate information than terminal marker programs alone. To assess biological interpretability, we examined genes with high posterior activity in the Bayesian CIPHER model. Fig. S17C shows that fate-associated genes form lineage-specific activity patterns across inferred fate forces, while Fig. S17B shows that the top CIPHER genes are enriched for canonical lineage marker sets with significant FDR-adjusted overlap for several fates. Together, these analyses suggest that CIPHER identifies biologically interpretable pre-commitment programs already present at day 4 that quantitatively predict clonal fate output at day 6.

#### 11 Correlations are enriched for protein and gene interactors

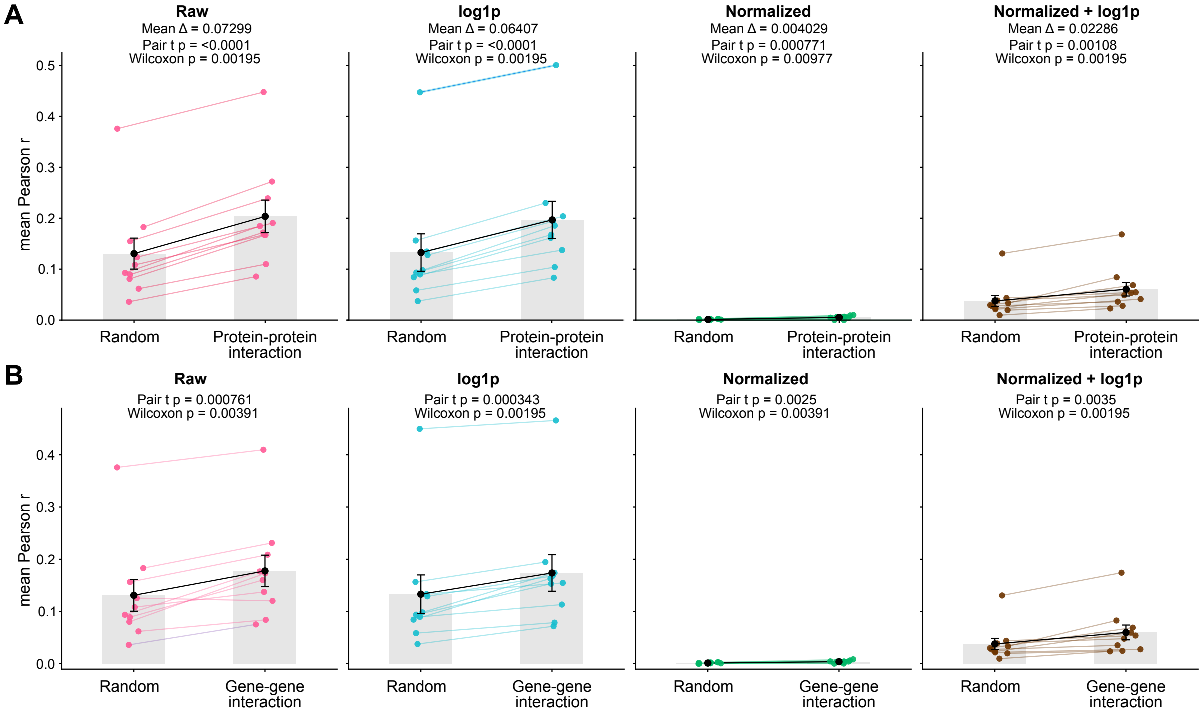


**Fig. S18**: **Known biological interactions produce more coherent gene-gene correlations.** (A) Mean Pearson correlations for gene pairs corresponding to proteins with known protein–protein interactions compared with randomly paired gene-gene correlations, evaluated under four preprocessing methods. Each point corresponds to a dataset, paired lines connect the same dataset, gray bars denote the group means, black points indicate the mean across datasets, and error bars show the standard error of the mean. Reported statistics correspond to paired $t$-tests and Wilcoxon signed-rank tests across datasets. (B) Same analysis for gene pairs with known gene–gene interactions.

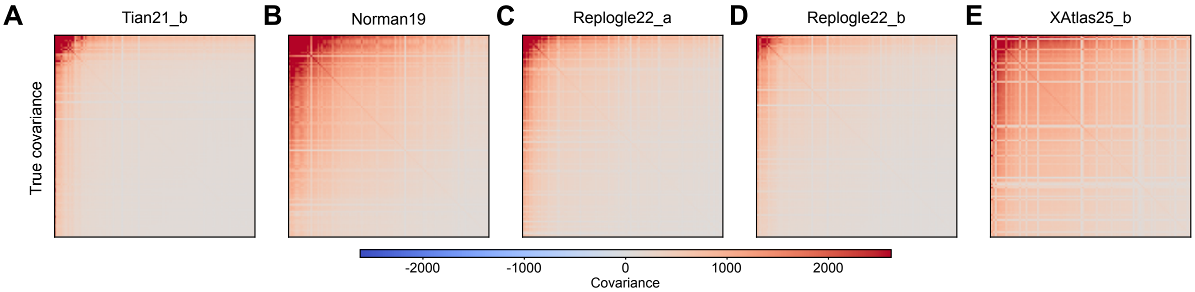


**Fig. S19**: **Representative control covariance matrices across Perturb-seq datasets.** (A–E) Gene–gene covariance matrices computed from control cells for five representative datasets: Tian21_b, Norman19, Replogle22_a, Replogle22_b, and XAtlas25_b. Genes are ordered identically within each dataset, and the color scale denotes covariance, with blue indicating negative covariance, white near-zero covariance, and red positive covariance. The matrices illustrate the heterogeneous covariance structure present across datasets, including broad positive covariance, localized blocks of highly correlated genes, and dataset-specific patterns of gene–gene co-fluctuation.
